# Genetic screens identify multiple negative regulators that additively restrict recruitment of the SWI/SNF chromatin remodeler to the *HO* promoter in *Saccharomyces cerevisiae*

**DOI:** 10.1101/606780

**Authors:** Emily J. Parnell, David J. Stillman

## Abstract

Activation of the *Saccharomyces cerevisiae HO* promoter is highly regulated, requiring the ordered recruitment of activators and coactivators and allowing production of only a few transcripts in mother cells within a short cell cycle window. We conducted genetic screens to identify the negative regulators of *HO* expression necessary to limit *HO* transcription. Known repressors of *HO* (Ash1 and Rpd3) were identified, as well as several additional chromatin-associated factors including the Hda1 histone deacetylase, the Isw2 chromatin remodeler, and the corepressor Tup1. We also identified clusters of *HO* promoter mutations that suggested roles for the Dot6/Tod6 (PAC site) and Ume6 repression pathways. We used ChIP assays with synchronized cells to validate the involvement of these factors and map the association of Ash1, Dot6, and Ume6 with the *HO* promoter to a brief window in the cell cycle between binding of the initial activating transcription factor and initiation of transcription. We found that Ash1 and Ume6 each recruit the Rpd3 histone deacetylase to *HO,* and their effects are additive. In contrast, Rpd3 was not recruited significantly to the PAC site, suggesting this site has a distinct mechanism for repression. Increases in *HO* expression and SWI/SNF recruitment were all additive upon loss of Ash1, Ume6, and PAC site factors, indicating the convergence of independent pathways for repression. Our results demonstrate that multiple protein complexes are important for limiting the spread of SWI/SNF mediated nucleosome eviction across the *HO* promoter, suggesting that regulation requires a delicate balance of activities that promote and repress transcription.

## Introduction

Eukaryotic gene expression is a highly regulated process requiring a variety of protein factors that ensure proper temporal and cell type control. Activated transcription is initiated by DNA-binding factors that recognize specific sequences in promoters and recruit multi-subunit coactivator complexes (Weake and Workman 2010). These coactivators contain enzymes that can post-translationally modify histones, alter the position of nucleosomes, or evict histones from the DNA, and thus positively influence transcription. Multi-subunit corepressors oppose the action of coactivators, also largely through histone modifications and altered nucleosome positioning, in this case to make the chromatin environment less favorable to expression. A delicate balance between activating and repressing processes is therefore a necessary component of complex gene regulation.

The yeast *Saccharomyces cerevisiae* provides a useful model system for studying this balance due to its ease of genetic manipulation. In both yeast and higher eukaryotes, the chromatin state is a key determinant of expression level, and chromatin remodelers and modifiers are evolutionarily conserved (Li *et al*. 2007; Weake and Workman 2010). However, most yeast genes have promoters that are relatively simple compared with those of more complex organisms. A notable exception is the *HO* gene (Stillman 2013), which has been extensively studied because it displays more sophisticated regulatory mechanisms typically seen in higher eukaryotes.

The *HO* gene encodes an endonuclease that initiates mating type interconversion in yeast by cleaving the mating type (*MAT*) locus, initiating *MAT* allele replacement via gene conversion (Strathern *et al*. 1982). Transcription of *HO* is highly regulated both temporally and by cell type, as inappropriate expression could lead to additional double stranded DNA breaks that would be deleterious to the cell (Stillman 2013). Yeast divides asymmetrically, giving rise to a larger mother cell and a smaller daughter cell from each mitotic division. Only haploid mother cells express the *HO* gene and are capable of switching their mating type, an evolutionary adaptation that allows mother/daughter pairs to mate and produce diploid progeny (Jensen *et al*. 1983; Nasmyth 1983). *HO* expression is also restricted to a narrow window in the cell cycle in late G1, with only a few transcripts produced per cell cycle (Nasmyth 1983; Miura *et al*. 2008).

Activation of *HO* expression is highly complex, involving sequential, ordered recruitment of several transcription factors and coactivators (Cosma *et al*. 1999; Bhoite *et al*. 2001; Takahata *et al*. 2009b). The *HO* promoter is unusually large relative to others in yeast, with a distance of 3 kb between the *HO* translation start site and the 3’ end of the upstream ORF. Binding sites for transcription factors have been identified in sequences up to approximately −1800 relative to the *HO* translation start site (Breeden and Nasmyth 1987; Stillman *et al*. 1988; Tebb *et al*. 1993), and a regulatory long noncoding RNA that originates at −2700 impacts *HO* promoter memory under specific conditions (Yu *et al*. 2016). Two upstream regulatory sequences (URS) are required for expression of *HO*, URS1 and URS2 (Nasmyth 1985). The Swi5 activator binds to two sites, A and B, within nucleosome depleted regions (NDRs) of URS1, at positions −1800 and −1300 (Fig 1A) (Stillman *et al*. 1988; Tebb *et al*. 1993; Jiang and Pugh 2009; Brogaard *et al*. 2012). The SCB binding factor (SBF), consisting of Swi4 and Swi6, associates with nine sites in URS2 (−900 to −200) (Taba *et al*. 1991); these sites are occluded by nucleosomes which must be evicted prior to binding of SBF (Takahata *et al*. 2009b; Yarrington *et al*. 2016).

**Figure 1.**
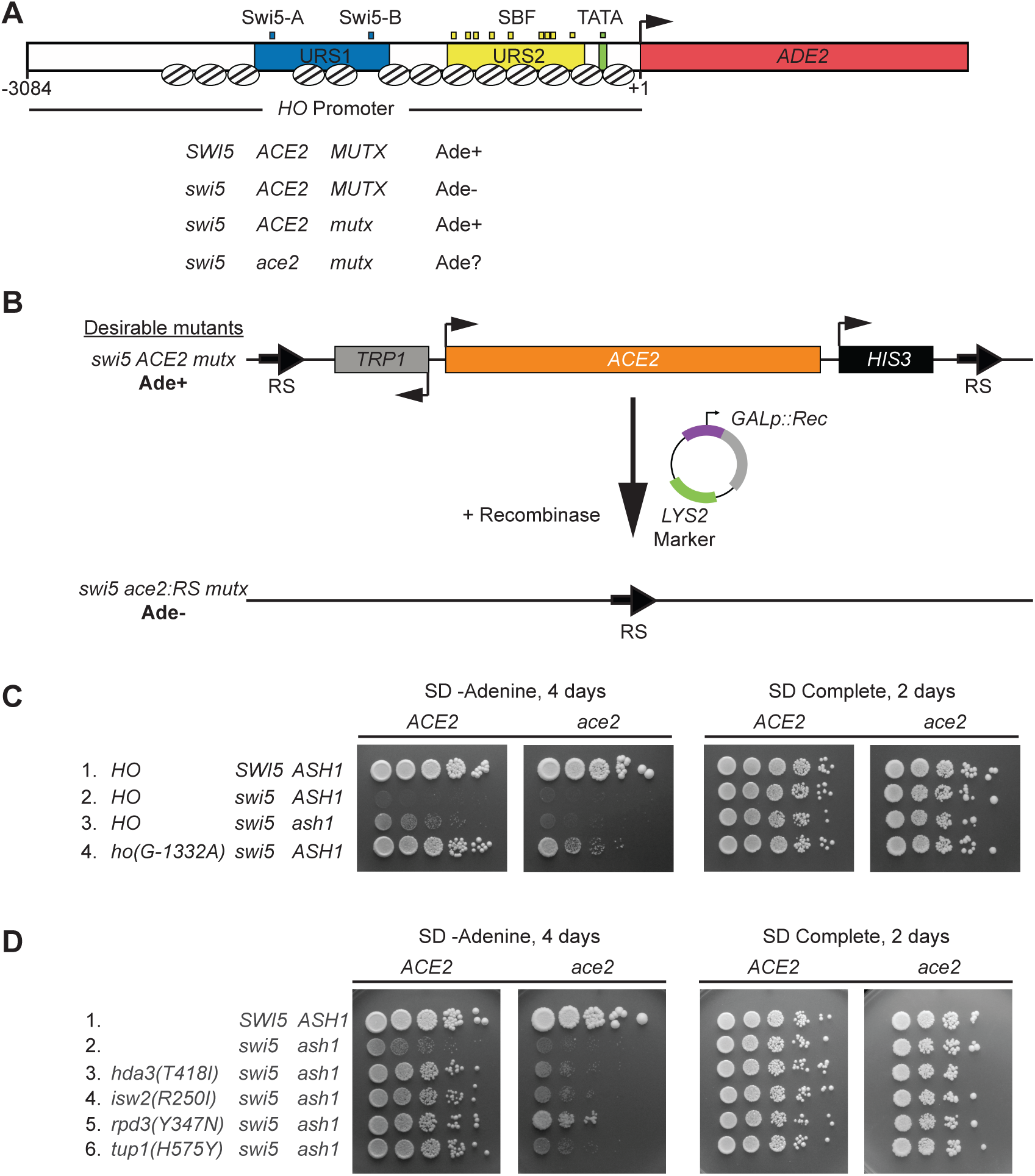
Genetic screen for mutants that allow activation of *HO-ADE2* in the absence of Swi5. (A) A schematic of the *HO-ADE2* reporter is shown. The *ADE2* ORF (red) replaced the *HO* ORF, preserving the 3084 possible nucleotides of the *HO* promoter, including Upstream Regulatory Sequences 1 (URS1; blue) and 2 (URS2; yellow). Small blue boxes = Swi5 binding sites; Small yellow boxes = SBF binding sites; Small green box = TATA. The arrow indicates the position of the ATG (+1). Well-defined nucleosome positions are shown throughout the region from approximately −2700 to +1 as ovals with lines. A strain that is *SWI5 ACE2* and wild type for other factors involved in *HO* expression (*MUTX*) grows on media lacking adenine (Ade+), due to expression of the *HO-ADE2* reporter. The strain used in the initial screen was *swi5* mutant and phenotypically Ade-, since *HO-ADE2* is very weakly expressed (*swi5 ACE2 MUTX*). Mutants identified in the screen (*mutx*) allowed *HO-ADE2* to be expressed in the *swi5* strain, making them Ade+ (*swi5 ACE2 mutx*). The *mutx* mutants identified in the screen were also tested for their ability to grow in the absence of adenine when lacking *ACE2 (swi5 ace2 mutx)*. (B) A schematic of the *ACE2* popout strategy is shown. The endogenous *ACE2* locus was modified to introduce RS recombinase target sites as well as *TRP1* and *HIS3* markers flanking *ACE2*. Mutant strains from the screen (*swi5 ACE2 mutx*) were transformed with a plasmid encoding the recombinase under the control of a galactose inducible promoter (*GAL*p::Rec). Subsequent growth on media containing galactose and selection for Trp-cells with 5-FAA allowed for isolation of strains lacking the *ACE2* gene (*swi5 ace2:RS mutx*). Assessment of the growth on -Ade was used to determine dependence of the Ade+ phenotype on the *ACE2* gene. Strains that were Ade+ with *ACE2* but Ade-with *ace2:RS* were kept for further analysis. (C, D) Shown are ten-fold serial dilutions, demonstrating the level of growth of wild type and mutant strains on media lacking adenine (SD -Adenine, left) and complete media (SD Complete, right). Strain genotypes are listed at the far left, and both the original strains (*ACE2*) and *ace2:RS* popout strains (*ace2*) are shown.

The initiating event for *HO* activation is the binding of Swi5 to URS1, which occurs when Swi5 enters the nucleus during anaphase (Nasmyth *et al*. 1990; Cosma *et al*. 1999). Swi5 recruits three coactivator complexes, the SWI/SNF chromatin remodeler, the SAGA complex containing the Gcn5 histone acetyltransferase, and Mediator; these coactivators are interdependent upon one another for full recruitment to *HO* (Cosma *et al*. 1999; Bhoite *et al*. 2001; Mitra *et al*. 2006). Following recruitment of coactivators, Swi5 is rapidly degraded (Tebb *et al*. 1993; Cosma *et al*. 1999). Remodeling of nucleosomes by SWI/SNF proceeds as a cascade of nucleosome evictions, first at URS1, then at the left end of URS2, and finally at the right half of URS2, allowing SBF to bind to its sites within URS2 (Takahata *et al*. 2009b; Yarrington *et al*. 2015). SBF also recruits the SWI/SNF, SAGA and Mediator coactivator complexes to extend the region of nucleosome eviction to the TATA region, ultimately allowing association of RNA polymerase. However, the final steps of activation are delayed until very late G1 by the association of inhibitors Whi5 and Stb1 with SBF, and their recruitment of the Rpd3 histone deacetylase complex to URS2 (Costanzo *et al*. 2004; De Bruin *et al*. 2004; Takahata *et al*. 2009a). At the end of G1, the Cdc28 cyclin dependent kinase (CDK) hyper-phosphorylates Whi5; Whi5 then leaves the nucleus and no longer represses *HO* transcription.

The unusually complex and regulated nature of *HO* promoter activation suggests it has a high barrier to transcription, maintained by potentially multiple repressive mechanisms. Early genetic screens identified the Sin3 and Rpd3 subunits of a histone deacetylase complex as negative regulators of *HO* (Nasmyth *et al*. 1987; Sternberg *et al*. 1987; Dorland *et al*. 2000), and subsequently Ash1 was identified as a critical determinant of daughter-specific repression of *HO* (Bobola *et al*. 1996; Sil and Herskowitz 1996). Ash1, a GATA-family DNA-binding protein that associates with the *HO* promoter, recruits the Rpd3 deacetylase to URS1, opposing the action of the Gcn5 coactivator and making the nucleosomes more restrictive to transcription (Maxon and Herskowitz 2001; Mitra *et al*. 2006; Takahata *et al*. 2011). We reasoned that the *HO* promoter likely has other mechanisms of repression to ensure that activation is terminated quickly, allowing only a few transcripts to be produced prior to re-binding of Swi5 in the next cell cycle. To identify other proteins that negatively affect *HO* expression, we employed a series of genetic screens designed to uncover mutations that would increase *HO* expression. We wanted to design a strain in which *HO* expression was driven by a very weak activator, such that it would be possible to easily identify increases in *HO* expression using a reporter system. We could then look for suppressor mutations that would allow the weak activator to function better, reasoning that these mutations may be in the types of negative factors we wished to identify. To design such a screen, we made use of the fact that the Swi5 pioneer transcription factor at *HO* has a paralogue, Ace2, that under normal circumstances fails to activate *HO* expression at a level that allows mating type switching.

The Swi5 and Ace2 proteins have nearly identical DNA-binding domains and bind to the same sequences *in vitro*, but *in vivo* they activate different genes (Dohrmann *et al*. 1992; Dohrmann *et al*. 1996; Voth *et al*. 2007). The *HO* gene is one example that is activated by Swi5 but not by Ace2 (Dohrmann *et al*. 1992). The similarity between Swi5 and Ace2 is low outside the zinc finger DNA-binding domain, and thus it is likely that the two factors interact differently with proteins, including coactivators. Swi5 may have the ability to recruit multiple factors that do not associate with Ace2 but are critical for overcoming the repression complexes at *HO*. We reasoned that by screening for mutants that would allow Ace2 to activate *HO*, we might identify inhibitory proteins that are important for restricting *HO* expression.

We therefore conducted a series of screens to identify mutants that would allow Ace2 to activate the *HO* gene in the absence of Swi5. The screens revealed a variety of *HO* promoter mutations as well as several protein factors previously shown to have roles in chromatin modification and transcriptional regulation. Combinations of mutants yielded higher levels of *HO* expression than single mutants. Thus, *HO* requires a host of both positive and negative factors that overlap functionally to collectively produce optimal coordination of transcription. Recruitment of Rpd3 appears to be critical, as at least two factors independently bring this protein to the *HO* promoter. The reduced repression observed in the mutants is likely to be the result of unopposed coactivator function. Consistent with this, we demonstrate that the mutants lead to additive increases in the level of association of SWI/SNF with *HO*.

## Materials and Methods

### Strain construction

All yeast strains used are listed in Supplemental Table S1 and are isogenic in the W303 background (*leu2-3,112 trp1-1 can1-100 ura3-1 ade2-1 his3-11,15*) (Thomas and Rothstein 1989). Standard genetic methods were used for strain construction (Rothstein 1991; Sherman 1991; Knop *et al*. 1999; Storici *et al*. 2001). Plasmids are listed in Supplemental Table S2. Oligos and further details regarding plasmid and strain construction are available upon request.

Strain DY10171 for the Ace2 popout genome-wide screen (*HO-ADE2 HO-CAN1 swi5::LEU2 RS::TRP1::ACE2::HIS3::RS ade2::HphMX*) was constructed from DY10085 (*HO-ADE2 HO-CAN1 swi5::LEU2 ade2::HphMX*) in a two-step process using plasmids containing *RS::TRP1::ACE2* (M5011) and *ACE2::HIS3::RS* (M5048). The *RS::TRP1::ACE2 and ACE2::HIS3::RS* plasmids were constructed in yeast using standard PCR and homologous recombination. The *RS::TRP1::ACE2::HIS3::RS* cassette was introduced into additional strains (for the *swi5 ash1* genome-wide screen, as well as the *HO* promoter targeted screens) using standard genetic crosses. Further details on construction of strains for the targeted *HO* promoter screens are provided below in the description of these screens.

The *ho(G-1332A)* strain DY10941 was constructed by first recovering the mutant *ho(G-1332A)* promoter from the screen strain using homologous recombination into linearized plasmid M4915 (*HO* in YCplac22) to generate M5181 (*ho(G-1332A)-ADE2* in YCplac22) and then transforming strain DY7874 *ho*[*URA3(−1496 to −1222 deleted)*]*::KanMX(3’)* to replace the *ho*[*URA3*] with the *ho(G-1332A)* promoter (Rothstein 1991). All other promoter mutants were constructed using a similar replacement strategy. The relevant portion of the *HO* promoter was amplified from each mutant strain and used to replace a *URA3* or *URA3-KanMX* cassette within the *HO* promoter by homologous recombination (see Supplemental Table S1 for strains used; DY14815, DY14198, and DY14840). Some promoter mutant strains had a *KanMX* marker present 3’ to the *HO* gene. Those that lacked this marker were subsequently transformed with a fragment from plasmid M4531 (*HO::KanMX(3’)* for tagging *HO* at the 3’ end with *KanMX*), so that mutant *HO* alleles could be followed in crosses. *HO* promoter mutations in all strains constructed via crosses were confirmed by quantitative PCR (qPCR) melting curve analysis (Wittwer *et al*. 2003) or by sequencing to verify the promoter mutation had not been separated from the 3’ marker by recombination during meiosis. The *ho(G-1332A, G-1142A)* double mutant was constructed using a two-step PCR in which *ho(G-1332A)* and *ho(G-1142A)* were PCR amplified from DY10941 and DY15937, respectively, forming overlapping products that were used for a final full-length PCR with the outermost primers. The PCR product containing both mutations was then transformed into strain DY14839 (*ho*[*URA3(K. lactis)::KanMX(inserted at - 1200)*] *ash1::LEU2 swi5::TRP1*) to replace the *URA3(K. lactis)::KanMX* marker. The strains used in Fig 10 have mutations at all nine SBF sites within URS2 (*9xSBFmut* is the combination of the *LX4* and *RX5* mutations described in Yarrington *et al*. 2016), to eliminate any recruitment of Rpd3 to the URS2 region by SBF.

Standard crosses were used to isolate the *tup1(H575Y)* allele from the strain in which the screen was conducted, using qPCR melt curve analysis (Wittwer *et al*. 2003) to differentiate the *tup1(H575Y)* allele from the wild type *TUP1* allele. Whole gene replacements of *DOT6, TOD6* and *UME6* were constructed in diploids using standard PCR integration methods to amplify markers and replace the coding regions (Longtine *et al*. 1998). Diploids were sporulated, and resulting haploids were used in subsequent standard genetic crosses to obtain all necessary haploid genotypes.

The *DOT6-V5* and *ASH1-V5* alleles were constructed by PCR amplifying and C-terminally integrating a V5 epitope tag with a *HIS3MX* marker from pZC03 (pFA6a-TEV-6xGly-V5-HIS3MX), provided by Zaily Connell and Tim Formosa (Addgene plasmid #44073). The *UME6-FLAG* allele was constructed by PCR amplifying and C-terminally integrating a 3x *FLAG::URA3::3x FLAG* cassette from ZM467 (Moqtaderi and Struhl 2008).

The *TOR1-1 fpr1::NATMX RPL13A-2 x FKBP12::TRP1* strain (Haruki *et al*. 2008) used for constructing Anchor Away strains was obtained from EUROSCARF (Y40343). The strain has two copies of *FKBP12* fused to the C-terminus of *RPL13A*, with a downstream *TRP1* marker, integrated at the *PMA1* locus. Rpl13A is a ribosomal protein that shuttles in and out of the nucleus and is therefore a suitable anchor for nuclear proteins. *TOR1-1* is a dominant allele of *TOR1* that allows growth in the presence of rapamycin. For Anchor Away experiments, the strain was first modified to integrate an *FRB* Rapamycin binding protein at the C-terminus of *CKS1*, *TUP1*, or *UME6* by PCR-amplifying the *FRB* domain and *KanMX* marker from plasmid pFA6a-FRB-KanMX (EUROSCARF P30578). The resulting strain was then used to construct additional genotypes using standard genetic crosses.

### Genetic screens: genome-wide

For each of the *swi5* and *swi5 ash1* Ace2-popout screens (using strains DY10171 and DY13589, respectively), independent cultures were grown to saturation overnight to allow formation of spontaneous mutants. Cells were plated at a low dilution onto synthetic complete medium lacking adenine with 2% dextrose (SD-Ade) to select for those that allowed activation of *HO-ADE2*. After 2 to 3 days at 30°C, Ade+ colonies were streaked to new SD-Ade plates alongside the wild type starting strain. Mutants displaying stronger growth than the wild type were tested for Ace2 dependence in the following manner. Strains were transformed with plasmid M5053, a YCp-*LYS2* plasmid containing the *Zygosaccharomyces rouxii* recombinase under the control of a galactose inducible promoter, and selected on SD-Lys media (Matsuzaki *et al*. 1990). Three transformants from each were patched to SD-Lys and then replica plated to synthetic complete medium lacking lysine with 2% galactose (SG-Lys) to activate transcription of the *Z. rouxii* recombinase. From SG-Lys, clones of each transformant were obtained on synthetic complete medium containing 2% dextrose and 5-fluoroanthranilic acid (SD+5-FAA) (Toyn *et al*. 2000), and colonies were then chosen from SD+5-FAA and tested for growth on SD-Leu (as a control for growth; strains are Leu+ due to either the *swi5::LEU2* or *ash1::LEU2* markers), SD-His (to confirm excision of the *RS::TRP1::ACE2::HIS3* cassette) and SD-Ade. The popout strains lacking Ace2 were then transformed with M2291, a YCp plasmid with *ACE2*, and confirmed to be Ade+ again with the re-introduction of Ace2. Strains that were Ade+ before popout (*RS::TRP1::ACE2::HIS3*), Ade-after popout (*ace2::RS*), and Ade+ after transformation with the *ACE2* plasmid (*ace2::RS*, *ACE2* YCp), were kept for further testing.

Each mutant strain was backcrossed to an appropriate strain marked with a drug-resistance gene at the 3’ end of *HO-ADE2* (strains DY10061 for *swi5* and DY13857 for *swi5 ash1*) to determine whether a single mutation was responsible for the phenotype, to assess possible linkage to *HO* and *ACE2* (using the *TRP1* and *HIS3* markers surrounding *ACE2* and the *HO* 3’ marker), and to generate *MAT***a** and *MAT*α strains for complementation and dominance testing. A cross to an *ace2* strain was used to confirm the Ace2 dependence of each mutation (strains DY10174 for *swi5* and DY13855 for *swi5 ash1*). Strains that were suspected to contain multiple mutations based on the initial backcross underwent additional backcrosses to separate mutations, and any of these that failed to show Ace2 dependence by a cross to an *ace2* strain were discarded. Each mutation was also determined to be either dominant or recessive by crossing to a *MAT* deletion strain (*MATΔ*; DY9341 for *swi5* strains and DY13891 for *swi5 ash1* strains) to produce *MAT***a**/*MATΔ* pseudo-diploids that could be tested for growth on SD-Ade. *HO* is repressed in a *MAT***a**/*MAT*α diploid by the a1/α2 heterodimer, but *HO* is expressed in a *MAT***a**/*MATΔ* pseudodiploid (Jensen *et al*. 1983). To generate strains for complementation testing, a drug resistance gene (*KanMX* or *NatMX*) was PCR amplified from pFA6-KanMX4 or pAG25-NATMX4 (Wach *et al*. 1994; Goldstein and Mccusker 1999) and used to replace the *MAT*α gene in specific mutants by homologous recombination. Complementation testing was performed by scoring the Ade+ phenotype of *MAT***a**/*MATΔ* pseudo-diploids generated by mating pairs of mutants.

The genes responsible for the Ade+ phenotype in mutants from the Ace2-popout screens were determined using multiple methods. *HO* and *ACE2* mutants were identified by linkage analysis, using the *TRP1* and *HIS3* markers surrounding *ACE2* and the drug resistance markers at the 3’ end of *HO*. The *HO* promoter or *ACE2* gene was then recovered from the mutant strain by allele rescue (Rothstein 1991) into a linearized plasmid (M4915, *HO* in YCplac22 and M2291, *ACE2* in YCplac33), followed by Sanger sequencing to determine the exact mutation(s). To identify the gene mutated in the 17 alleles comprising a single complementation group from the *swi5* screen, we first used chromosome loss-based gene mapping (Reid *et al*. 2008) to determine that the relevant gene was located on chromosome XI. Since *ASH1* is located on chromosome XI and was known to be a repressor of *HO* transcription, we then specifically tested *ASH1* by plasmid complementation using a YIp-*URA3* plasmid with *ASH1* (M5404).

Affected genes for eight of the mutants from the *swi5 ash1* screen were determined by Illumina whole-genome sequencing. Yeast DNA was isolated using a combination protocol involving lyticase treatment and precipitation/extraction of proteins with potassium acetate and phenol, followed by further purification using QIAGEN DNeasy columns (69504; Supplementary Protocol for Yeast, starting with addition of AL Buffer). A pellet from a 50 mL culture of cells at approximately OD 1.0 was resuspended in an equal volume of 20 mM Tris, pH 8.0, 100 mM EDTA, 0.5 M 2-mercaptoethanol, and cells were incubated with 500 units of lyticase for 30 min at 37°C, vortexing intermittently. Lysis buffer (100 mM Tris, pH 8.0, 50 mM EDTA, 3% SDS, 1 mg/ml proteinase K) was added, and the mixture was incubated for 15 min at 65°C and cooled. Proteins were precipitated using 5 M potassium acetate, followed by phenol-chloroform extraction and ethanol precipitation. DNA was treated with RNaseA and then purified with a QIAGEN DNeasy Blood and Tissue Kit (69504; see above). Purified DNA was sonicated and used for Illumina library preparation and sequencing with an Illumina GAIIx 75 cycle single-end run (University of Utah High Throughput Genomics Facility).

High throughput sequences from each mutant strain were aligned to the *S. cerevisiae* S288C sequence (SGD release R63-1-1) using Novocraft *novoalign* (http://www.novocraft.com, version 2.7). Variants within each strain were identified using *samtools* (https://github.com/samtools/, release 0.1.19) mpileup function and *bcftools* (part of samtools 0.1.19). Variants were filtered with *vcfutils.pl* (part of samtools 0.1.19). Common and unique variants between the mutant strains were identified using the script *intersect_SNPs.pl* (biotoolbox-legacy, https://github.com/tjparnell/biotoolbox-legacy). Unique variants were annotated with *locate_SNPs.pl* (biotoolbox-legacy), and variants not eliciting a codon change were discarded. Variants present in at least 50% of the reads from a given strain were evaluated for quality by viewing the alignments in a genome browser. Following analysis, each mutant produced a list of one to five candidate genes, for which all high-quality reads showed the mutation. Candidates were then tested by plasmid complementation and/or linkage analysis. Mutants identified by whole genome sequencing include *tup1(H575Y)*, *hda3(T418I)*, *isw2(R250*)*, 3 independent frame-shift mutations within *ITC1*, and 2 mutations within *SFP1* (one in the 5’ UTR and one stop mutation; see also Table 1).

**Table 1.**
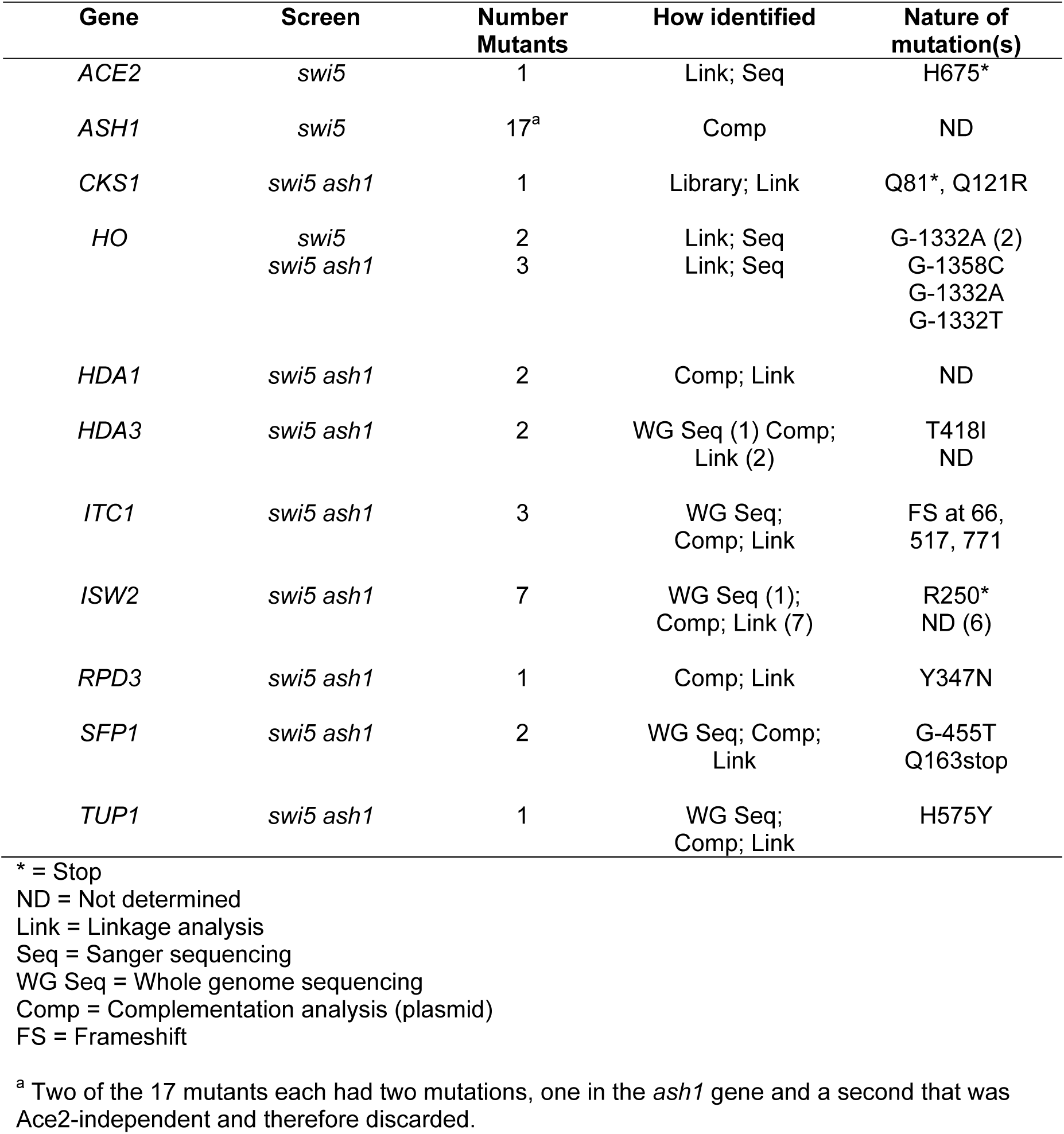
Mutants identified in screens for factors that increase *HO-ADE2* expression in the <u>absence of Swi5.</u>

After mutations in some strains were identified by genomic sequencing, strains with unidentified mutations were tested for alleles of the same genes (*hda1*, *hda3*, *isw2*, *rpd3*) by plasmid complementation testing and linkage analysis (Table 1). For most of these mutants, the gene was not sequenced to determine the precise nature of the mutation. *CKS1* was identified by screening the ATCC library #37415 (15-20 kb inserts in YCp50, *URA3*) for a complementing clone, which was then sequenced to determine the relevant genes. Plasmid complementation then identified *CKS1* as the gene responsible for complementation of the mutant phenotype, and this was confirmed by linkage analysis.

### Genetic screens: HO promoter

Strains for the targeted *HO* promoter screen were designed to allow for integration of a mutagenized *HO* promoter PCR product at the *HO-ADE2* locus. A portion of the *HO* promoter in *HO-ADE2* was replaced or deleted in these strains, rendering the promoter non-functional. Mutagenic PCR of a plasmid containing the *HO* promoter generated fragments that spanned from −1827 to −268, which were used to replace the promoter in the non-functional *HO-ADE2* strains, allowing for a functional *HO-ADE2* to be produced by homologous recombination.

Three strains were constructed and used for screens, one with an internal deletion within the *HO* promoter (*swi5* only; DY13862) and two with a replacement of a portion of the *HO* promoter with the *sup4-o* tRNA gene (*swi5*; DY13863 and *swi5 ash1*; DY13925). The precursor strain for all three was constructed by replacing a portion of the *HO* promoter from −1496 to −1222 with a *URA3* marker (strain DY13832; using M2026, *ho*[*URA3(−1496 to −1222 deleted)*] disruptor). Subsequently, the *ho*[*URA3*] was replaced with either an *HO* promoter with an internal deletion (strain DY13862; using M346, deletion of *HO* promoter from −1496 to −1130) or an *HO* promoter containing a *sup4-o* tRNA gene (strain DY13863; using M3208, where *sup4-o* replaces *HO* sequences from −1496 to −1222). The *sup4-o* is an ochre suppressor tRNA that allows read-through of the *can1-100* nonsense allele, making the strains effectively *CAN1+* (Nasmyth 1985). Can1 is a transporter that confers sensitivity to the drug canavanine, a toxic analog of arginine. Replacement of part of the *HO* promoter with *sup4-o* thereby rendered the strains canavanine sensitive, and subsequent homologous recombination with mutant *ho* promoter PCR sequence to eliminate *sup4-o* allowed the strains to become resistant to canavanine.

For each screen, multiple *HO* promoter PCR reactions were each transformed separately to allow for later determination of mutants that had arisen independently during PCR. Growth on SD-Ade was used to select colonies that displayed stronger activation of *HO-ADE2* than the wild type *HO* promoter. Ade+ colonies from screens with strains containing a *sup4-o* in the *HO-ADE2* promoter were also tested for growth on canavanine to confirm that the promoter had been replaced. Candidates were then tested for Ace2-dependence of the Ade+ phenotype using the strategy for Ace2 popout and add-back described above. For those that showed Ace2-dependence, genomic DNA was isolated, and a region spanning the replaced portion of the *HO* promoter was PCR amplified and subjected to Sanger sequencing to identify mutations. A number of strains had more than one mutation. For most, we were able to deduce the mutation likely causing the Ade+ phenotype by comparison with other strains that contained only a single mutation in either an identical nucleotide or one in close proximity (Table 3). Others were determined by testing single mutations for effects on *HO* expression in the endogenous context (Table 4).

**Table 2.**
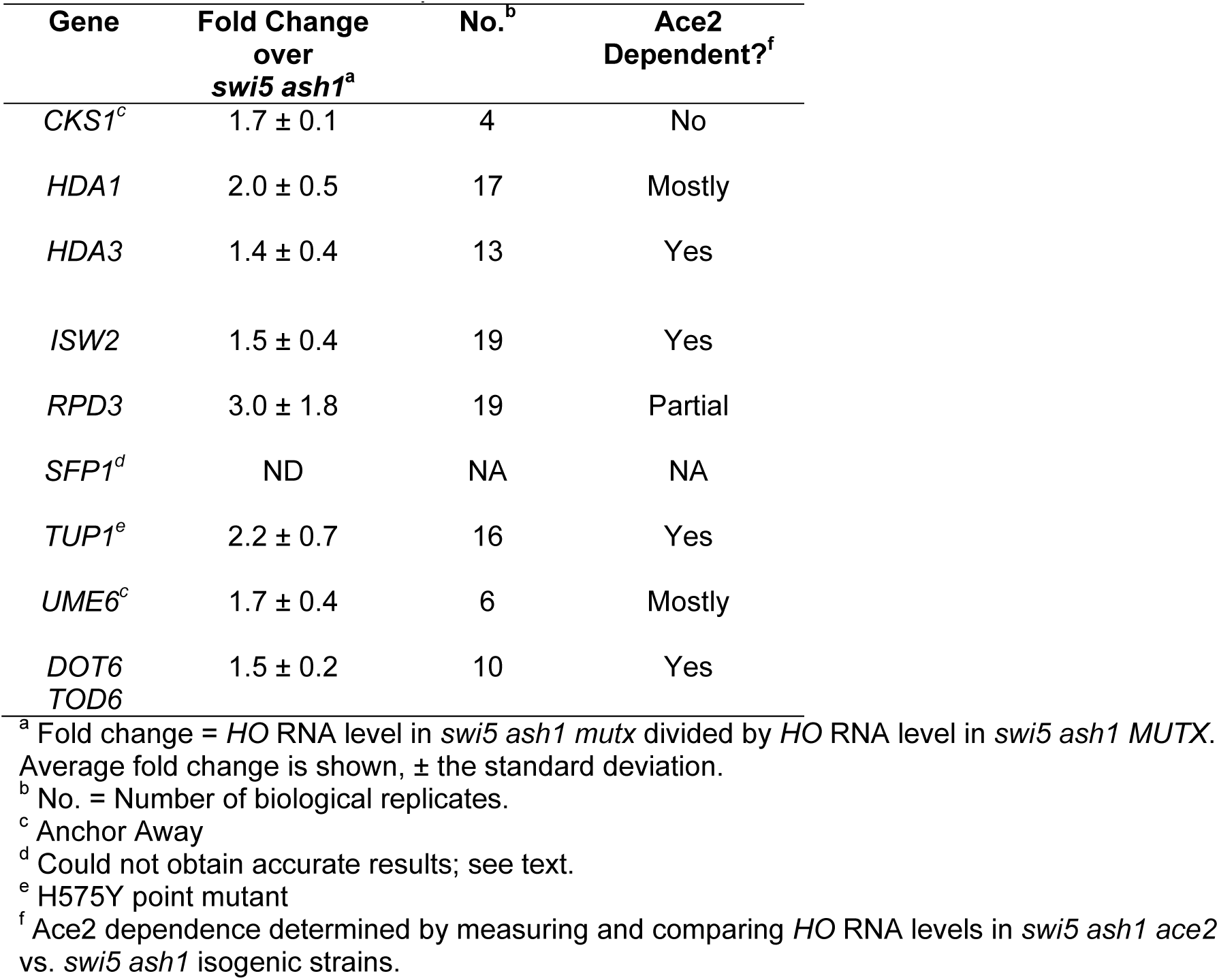
Effect of mutants on expression of *HO* in *swi5 ash1*.

| Gene | Fold Change<br>over<br>*swi5 ash1*<sup>a</sup> | No.<sup>b</sup> | Ace2<br>Dependent?<sup>f</sup> |
| --- | --- | --- | --- |
| *CKS1*<sup>c</sup> | 1.7 ± 0.1 | 4 | No |
| *HDA1* | 2.0 ± 0.5 | 17 | Mostly |
| *HDA3* | 1.4 ± 0.4 | 13 | Yes |
| *ISW2* | 1.5 ± 0.4 | 19 | Yes |
| *RPD3* | 3.0 ± 1.8 | 19 | Partial |
| *SFP1*<sup>d</sup> | ND | NA | NA |
| *TUP1*<sup>e</sup> | 2.2 ± 0.7 | 16 | Yes |
| *UME6*<sup>c</sup> | 1.7 ± 0.4 | 6 | Mostly |
| *DOT6*<br>*TOD6* | 1.5 ± 0.2 | 10 | Yes |

<sup>a</sup> Fold change = *HO* RNA level in *swi5 ash1 mutx* divided by *HO* RNA level in *swi5 ash1 MUTX*. Average fold change is shown, ± the standard deviation.
<sup>b</sup> No. = Number of biological replicates.
<sup>c</sup> Anchor Away
<sup>d</sup> Could not obtain accurate results; see text.
<sup>e</sup> H575Y point mutant
<sup>f</sup> Ace2 dependence determined by measuring and comparing *HO* RNA levels in *swi5 ash1 ace2* vs. *swi5 ash1* isogenic strains.

**Table 3.**
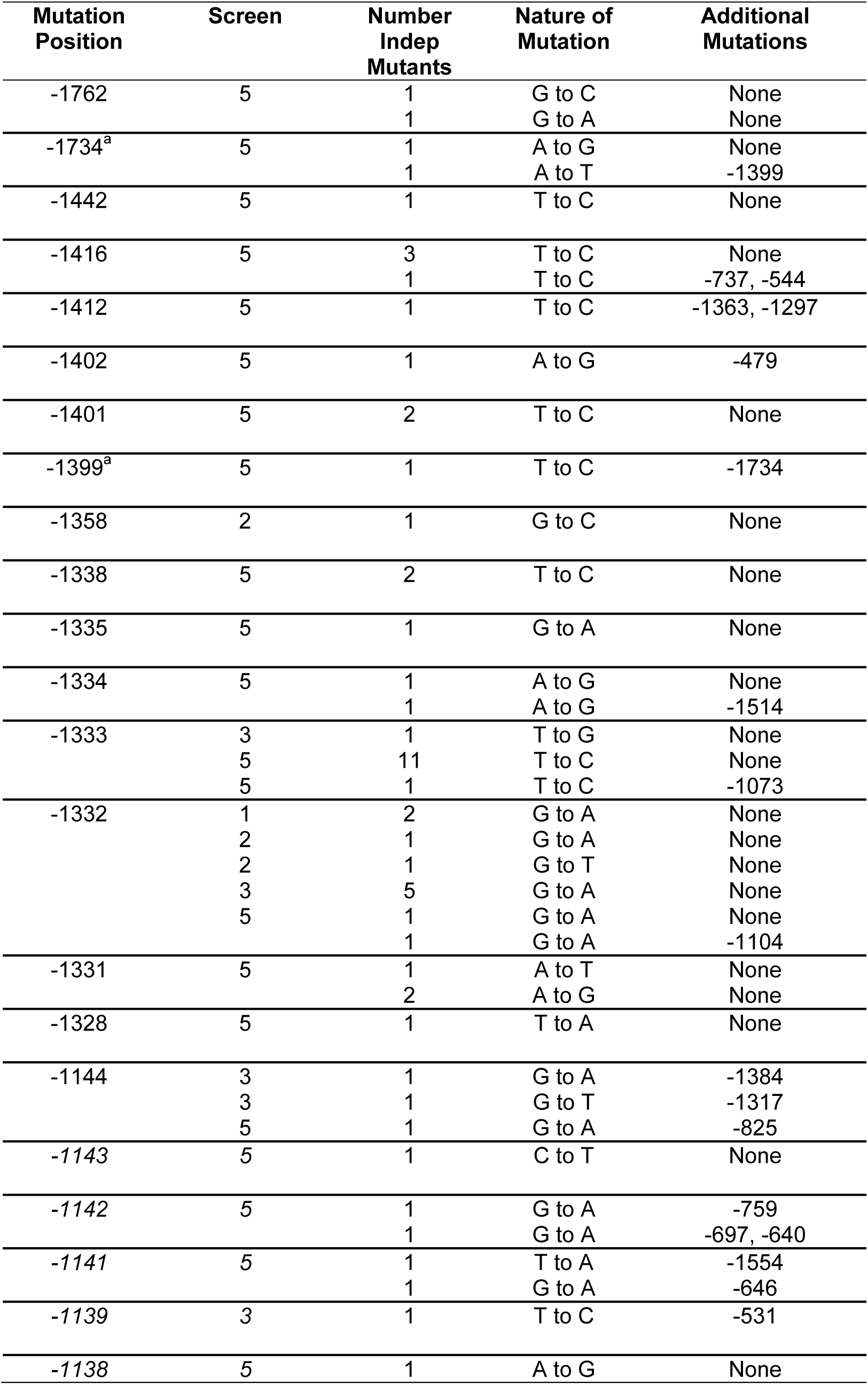
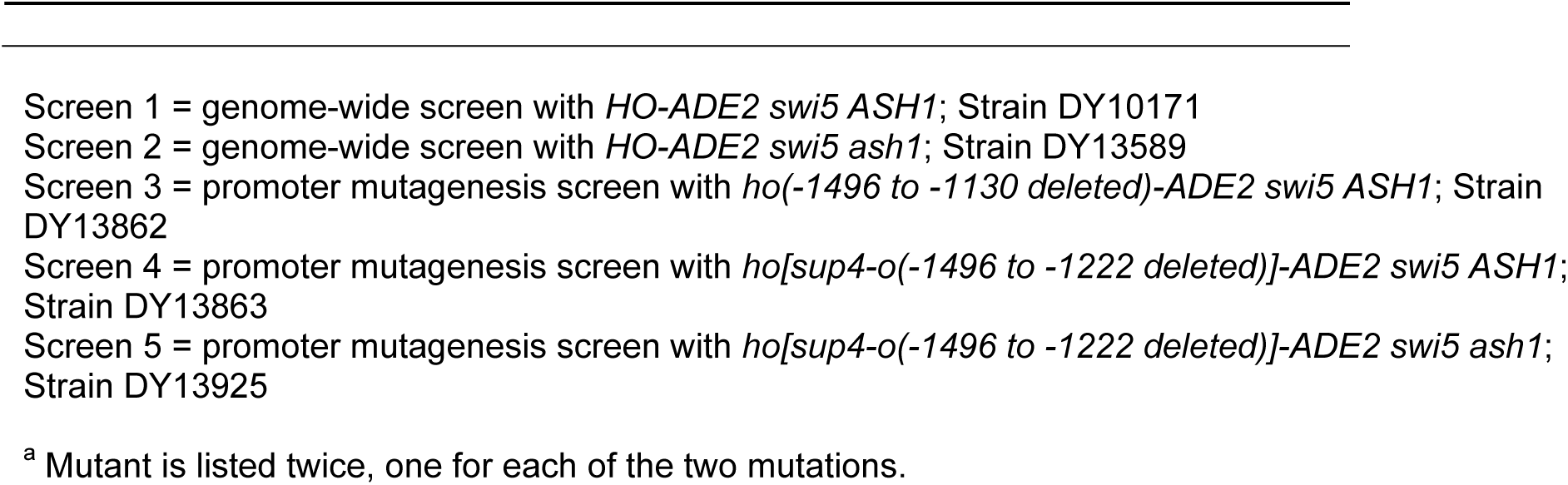
*HO* promoter mutants identified in screens for suppressors of *swi5* or *ash1 swi5*.

| **Mutation Position** | **Screen** | **Number Indep Mutants** | **Nature of Mutation** | **Additional Mutations** |
| --- | --- | --- | --- | --- |
| -1762 | 5 | 1 | G to C | None |
|  |  | 1 | G to A | None |
| -1734<sup>a</sup> | 5 | 1 | A to G | None |
|  |  | 1 | A to T | -1399 |
| -1442 | 5 | 1 | T to C | None |
| -1416 | 5 | 3 | T to C | None |
|  |  | 1 | T to C | -737, -544 |
| -1412 | 5 | 1 | T to C | -1363, -1297 |
| -1402 | 5 | 1 | A to G | -479 |
| -1401 | 5 | 2 | T to C | None |
| -1399<sup>a</sup> | 5 | 1 | T to C | -1734 |
| -1358 | 2 | 1 | G to C | None |
| -1338 | 5 | 2 | T to C | None |
| -1335 | 5 | 1 | G to A | None |
| -1334 | 5 | 1 | A to G | None |
|  |  | 1 | A to G | -1514 |
| -1333 | 3 | 1 | T to G | None |
|  | 5 | 11 | T to C | None |
|  | 5 | 1 | T to C | -1073 |
| -1332 | 1 | 2 | G to A | None |
|  | 2 | 1 | G to A | None |
|  | 2 | 1 | G to T | None |
|  | 3 | 5 | G to A | None |
|  | 5 | 1 | G to A | None |
|  |  | 1 | G to A | -1104 |
| -1331 | 5 | 1 | A to T | None |
|  |  | 2 | A to G | None |
| -1328 | 5 | 1 | T to A | None |
| -1144 | 3 | 1 | G to A | -1384 |
|  | 3 | 1 | G to T | -1317 |
|  | 5 | 1 | G to A | -825 |
| -1143 | 5 | 1 | C to T | None |
| -1142 | 5 | 1 | G to A | -759 |
|  |  | 1 | G to A | -697, -640 |
| -1141 | 5 | 1 | T to A | -1554 |
|  |  | 1 | G to A | -646 |
| -1139 | 3 | 1 | T to C | -531 |
| -1138 | 5 | 1 | A to G | None |

Screen 1 = genome-wide screen with *HO-ADE2 swi5 ASH1*; Strain DY10171 Screen 2 = genome-wide screen with *HO-ADE2 swi5 ash1*; Strain DY13589 Screen 3 = promoter mutagenesis screen with *ho(-1496 to -1130 deleted)-ADE2 swi5 ASH1*; Strain DY13862 Screen 4 = promoter mutagenesis screen with *ho[sup4-o(-1496 to -1222 deleted)]-ADE2 swi5 ASH1*; Strain DY13863 Screen 5 = promoter mutagenesis screen with *ho[sup4-o(-1496 to -1222 deleted)]-ADE2 swi5 ash1*; Strain DY13925
<sup>a</sup> Mutant is listed twice, one for each of the two mutations.

**Table 4.** Effect of *HO* mutants on expression in *swi5 ash1*.

| Mutation Position | Cluster | Nucleotide | Fold Change over *swi5 ash1*<sup>a</sup> | No.<sup>b</sup> | Ace2 Dependent?<sup>c</sup> |
| --- | --- | --- | --- | --- | --- |
| -1762 | NA | G | 1.3 ± 0.0 | 2 | Mostly |
| -1734 | NA | A | 3.6 ± 0.9 | 2 | Yes |
| -1442 | NA | T | 3.0 ± 0.2 | 2 | Mostly |
| -1416 | 1 | T | 3.4 ± 0.9 | 6 | Mostly |
| -1401 | 1 | T | 3.4 ± 1.0 | 8 | Yes |
| -1399 | 1 | T | 1.7 ± 0.1 | 2 | Mostly |
| -1358 | NA | G | 2.2 ± 0.3 | 2 | Mostly |
| -1338 | 2 | T | 3.1 ± 1.2 | 2 | Yes |
| -1335 | 2 | G | 3.4 ± 0.5 | 2 | Mostly |
| -1334 | 2 | A | 5.3 ± 1.4 | 4 | Mostly |
| -1333 | 2 | T | 4.1 ± 0.7 | 4 | Mostly |
| -1332 | 2 | G | 6.4 ± 2.3 | 25 | Mostly |
| -1331 | 2 | A | 4.2 ± 2.4 | 4 | Mostly |
| -1328 | 2 | T | 4.7 ± 1.0 | 2 | Mostly |
| -1144 | 3 | G | 2.4 ± 0.4 | 4 | Yes |
| -1143 | 3 | C | 2.4 ± 0.3 | 4 | Mostly |
| -1142 | 3 | G | 4.3 ± 1.2 | 18 | Mostly |
| -1141 | 3 | G | 2.6 ± 0.5 | 4 | Mostly |
| -1138 | 3 | A | 2.8 ± 0.6 | 2 | Mostly |

<sup>a</sup> Fold change = *HO* RNA level in *swi5 ash1 ho(mut)* divided by *HO* RNA level in *swi5 ash1 HO*. Average fold change is shown, ± the standard deviation.
<sup>b</sup> No. = Number of biological replicates.
<sup>c</sup> Ace2 dependence determined by measuring and comparing *HO* RNA levels in *swi5 ash1 ace2* vs. *swi5 ash1* isogenic strains.

### Growth Assays

For plate spot dilution assays, liquid cultures of the indicated strains were grown to saturation, serially diluted in 10-fold increments, spotted onto SD-Ade or SD Complete media for the number of days indicated, and photographed.

### RNA Expression and Chromatin Immunoprecipitation (ChIP) Analysis

For logarithmic cell collection (OD_660_ of 0.6 to 0.8), cells were grown at 30°C in YPA medium (1% yeast extract, 2% bactopeptone, 0.002% adenine) supplemented with 2% dextrose (Sherman 1991). Cell cycle synchronization was performed by galactose withdrawal and readdition with a *GALp::CDC20* strain grown at 25°C in YPA medium containing 2% galactose and 2% raffinose (Bhoite *et al*. 2001). Synchrony was confirmed by microscopic analysis of budding indices and analysis of cell-cycle regulated mRNAs (data not shown). For Anchor Away experiments, cells were grown to an OD_660_ of 0.3 to 0.4 in YPA media containing 2% glucose at 30°C. The cultures were then split into two, and rapamycin (1μg/mL final concentration, dissolved in ethanol; LC Laboratories; +Rapamycin) or ethanol alone (-Rapamycin) was added to the media for the final two hours of growth prior to collection of cells at an OD_660_ of 0.8 to 1.0.

RNA was isolated from either logarithmically growing cells or synchronized cells, and *HO* mRNA levels were measured by reverse transcription quantitative PCR (RT-qPCR), as described previously (Voth *et al*. 2007). *HO* RNA expression was normalized to that of *RPR1. RPR1* encodes the RNA component of RNase P and is transcribed by RNA polymerase III. Most genetic manipulations that affect RNA Pol II transcription do not affect transcription of *RPR1*. For logarithmic cells, normalized *HO* RNA expression values were graphed relative to wild type (WT) expression.

ChIPs were performed as described (Bhoite *et al*. 2001; Voth *et al*. 2007), using mouse monoclonal antibodies to the V5 epitope (SV5-Pk1; Abcam), the FLAG epitope (M2; Sigma), or the Myc epitope (4A6; Upstate) and antibody-coated magnetic beads (Pan Mouse IgG beads; Life Technologies). Cells from either logarithmically growing cells or synchronized cells were cross-linked in 1% formaldehyde for 20 min at room temperature and quenched with 125 mM glycine. ChIP signals for relevant target genes within each sample were first normalized to either an expected negative reference control or a known positive reference control, and then to their respective input controls. Normalization to the negative reference controls (*IGR-I* intergenic region of chromosome I and *CDC2* coding region) yielded a high degree of variability between ChIP experiments due to normalization to a very small number. However, this normalization was useful for observing the presence of a ChIP signal over the background from a strain lacking the V5 or FLAG epitope tag (Figs 5C-E, 6B, 7, 8C, S1, S2, S4, S5). Normalization to a positive reference resulted in less variability among samples. Positive references used include: *GCD10* for Dot6 (the promoter region of *GCD10*, shared with that of *NOP2*, has two PAC sites), *INO1* for Ume6 and Rpd3 (*INO1* is a well-established target of Ume6; Slekar and Henry 1995; Jackson and Lopes 1996), *CLN3* for Ash1 (*CLN3* is bound by Ash1 by ChIP analysis; Di Talia *et al*. 2009; Zapata *et al*. 2014), *CTS1* for Swi2, and *EGT2* for Swi5 (Figs 5B, 6A, 8A, 8B, 10, 11, S7). For figures using a negative reference control, all values were graphed relative to the No Tag control. For figures using a positive reference control, all values were graphed relative to the wild type control.

Quantitative PCR (qPCR) experiments for both RNA and ChIP analysis were run on a Roche Lightcycler 480, and concentrations were determined using wild type cDNA or ChIP input for in-run standard curves via the E-method (Tellmann 2006). Error bars represent the standard deviation of at least three biological samples. The Student’s t-test was used to determine significance of changes in *HO* expression and factor binding between different genotypes. For all comparisons mentioned in the Results and Discussion, *p*-values are indicated in the figures. Primers for all experiments are available upon request. For ChIP tiling PCR across the *HO* promoter, ChIP time courses with Ash1, Dot6 and Ume6, and time course *HO* expression, a single sample was shown in the main figures for simplicity (Fig 5C, 6B, 7). Triplicate biological samples for the time course ChIP and RNA experiments are shown in Supplemental Figure S4. The time course Dot6 ChIP in wild type and the *ho(G-1332A)* mutant shown in Fig 5D,E was repeated in multiple independent experiments; one is shown for simplicity (note that for wild type, additional experiments are shown in Fig S4).

### Data availability

Strains and plasmids are available upon request. Table S1 lists the strains and Table S2 lists the plasmids used in this study. Seven Supplemental Figures and two Supplemental Tables were uploaded to the GSA Figshare portal.

## Results and Discussion

### Genetic screen to identify mutants that allow Ace2 to activate the HO gene

We used an *HO-ADE2* reporter to perform a genetic screen to identify mutations that would allow Ace2 to activate *HO* expression in the absence of the Swi5 activator. *ADE2* encodes an enzyme necessary for biosynthesis of adenine and has been used previously as a reporter for expression of genes in yeast (Jansen *et al*. 1996). When all *ADE* genes are wild type, the cells biosynthesize adenine and are capable of growth on media that lacks adenine. Cells that have an *ade2* mutation do not synthesize adenine and become dependent on its presence in the media; hence, they fail to grow on media lacking adenine (-Ade). Previous studies have demonstrated that *ADE2* is a useful model for *HO* expression, as modest levels of *HO-ADE2* transcription allow robust growth on -Ade media, while low transcript levels do not confer growth on -Ade, and intermediate levels allow for slow growth (Jansen *et al*. 1996). Therefore, it is possible to observe alterations in expression of *HO-ADE2* by the ability of the strains to grow on media lacking adenine.

The *ADE2* open reading frame was integrated at the genomic *HO* locus, replacing the *HO* ORF, to construct the *HO-ADE2* reporter (Fig 1A). All 5’ and 3’ regulatory sequences necessary for *HO* expression are present, notably the entire 3084 nucleotides of possible promoter region extending to the ORF of the upstream gene. Our screen was performed in an *HO-ADE2 swi5 ACE2* strain. This strain has very low *HO* expression, due to the absence of the Swi5 activator; hence, the cells are phenotypically Ade-(*swi5 ACE2 MUTX*; Fig 1A). We screened for spontaneous mutants that would confer some level of growth on -Ade media (Ade+; *swi5 ACE2 mutx*; Fig 1A).

We anticipated that mutations in a variety of general chromatin and transcription factors could increase *HO* expression by affecting the nucleosome structure at URS2, which could in turn allow premature binding and/or activation by SBF, regardless of whether Ace2 promoted transcription. To avoid obtaining hits in such generic processes, which should be independent of Ace2, we devised a “popout” strategy to quickly remove *ACE2* from the mutants to compare phenotypes in the presence and absence of Ace2 (Fig 1B). The *ACE2* locus was modified by introducing *Z. rouzii* RS recombinase target sites and *TRP1* and *HIS3* marker genes flanking *ACE2*. The 31 nucleotide RS sites are targets for the *Z. rouzii* recombinase, thereby “popping out” the *ACE2* gene as well as the two markers in a small percentage of cells (Matsuzaki *et al*. 1990; Roca *et al*. 1992). Cells lacking the *TRP1-ACE2-HIS3* cassette were selected on 5-fluoroanthranilic acid (5-FAA), which selectively kills cells that express *TRP1* (Toyn *et al*. 2000). The subsequent strains lacking the *TRP1-ACE2-HIS3* cassette were then re-tested for their growth on -Ade to determine the level of *HO-ADE2* expression (*swi5 ace2 mutx*; Fig 1A). This allowed us to identify and discard those whose increased *HO-ADE2* expression was Ace2-independent.

We obtained 124 mutants from the initial genetic screen, 20 of which remained after further testing (Table 1, *swi5* screen). Most were eliminated because their Ade+ phenotype was not dependent on Ace2, as determined either by the popout recombinase test or by a genetic cross to introduce an *ace2* deletion allele. A small subset of the mutants (about 20) were removed because the phenotype of these strains did not remain consistently Ade+ upon repeated further testing. We speculate that these strains became aneuploid due to a mutation in the gene encoding Ume6, a factor that we later found to be involved in negative regulation of *HO* (see below).

Complementation testing revealed that 17 of the 20 desirable mutants likely had alterations in the same gene. We identified the mutated gene in the 17 strains as *ASH1*, the previously known repressor and determinant of mother-cell specific *HO* expression (Table 1, *swi5* screen) (Bobola *et al*. 1996; Sil and Herskowitz 1996). The identification of *ASH1* mutants in our screen was not surprising, given that both Ash1 and Ace2 are largely daughter-specific proteins (Bobola *et al*. 1996; Colman-Lerner *et al*. 2001). The Ash1 protein is translated predominately in daughter cells, as the *ASH1* mRNA is transported through the bud neck and anchored to the distal tip of the daughter cell via the She proteins (Bertrand *et al*. 1998; Munchow *et al*. 1999). Ace2 accumulates selectively in the daughter cell nucleus due to masking of its nuclear export sequence (NES) by the daughter-specific protein Cbk1 (Mazanka *et al*. 2008). The Ash1 protein therefore presents a barrier to Ace2 primarily in daughter cells, so removal of Ash1 could allow Ace2 to inappropriately activate *HO* in daughter cells, increasing the number of cells expressing *HO* without altering the function of the promoter. However, the level of *HO-ADE2* activation in a *swi5 ash1* mutant was still very weak relative to wild type (Fig 1C, Line 3 *ACE2*).

The three remaining Ace2 dependent mutants were identified through linkage analysis; one had a mutation in the *ACE2* gene itself, and two had mutations in the *HO* promoter (Table 1, *swi5* screen). The *ace2(H675*)* allele is a truncation within the C2HC zinc finger of Ace2 and likely reduces its interaction with partner proteins needed for activation of target genes, including *ASH1* (data not shown). The two *HO* promoter mutants each have substitutions of “G” to “A” at position −1332 relative to the *HO* start site of translation. This *ho(G-1332A)* mutation is within a nucleosome depleted region (NDR), 24 nucleotides upstream of the Swi5 site B. The *ho(G-1332A)* promoter mutants had a much stronger Ade+ phenotype than the *ash1* mutants, with a growth level approaching that of wild type (Fig 1C, Line 4 *ACE2*). The *ho(G-1332A)* position could be located within a binding site for Ash1, or for another protein complex that negatively affects *HO* expression.

We next examined whether deletion of *ash1* or the *ho(G-1332A)* mutation could affect endogenous *HO* gene expression in a native context, i.e. without the *ADE2* reporter. We also constructed a *swi5 ash1 ho(G-1332A)* strain to determine whether the *ash1* and *ho(G-1332A)* mutations were additive. If the *ho(G-1332)* position is part of a binding site for Ash1, we expect the effect of the two mutations to be similar, and the two mutations together would not have a stronger effect than either single mutation. *HO* expression measured using RT-qPCR was severely reduced in the *swi5* mutant, to 5% of wild type (Fig 2, Line 2 blue). The *ash1* null increased expression of *HO* above *swi5* by 4-fold, demonstrating suppression, just as had been observed in the *HO-ADE2* strain (compare Lines 2 and 4 blue). The *ho(G-1332A)* mutant showed much more robust suppression, up to approximately half of the normal wild type level of *HO* expression, nearly 10-fold over that of *swi5* alone (compare Lines 2 red and blue). The combination of *ash1* and *ho(G-1332A)* was additive, raising *HO* expression to over 100% (Line 4 red). Thus it is likely that the protein(s) bound at *ho(G-1332)* repress *HO* through a mechanism independent of Ash1. Similar to the spot dilution assays, the suppression by *ash1* and *ho(G-1332A)* is mostly, but not completely, dependent upon Ace2, because *ace2* mutants showed significantly reduced *HO* expression (compare Lines 2 and 3 red and Lines 4 and 5 blue).

**Figure 2.**
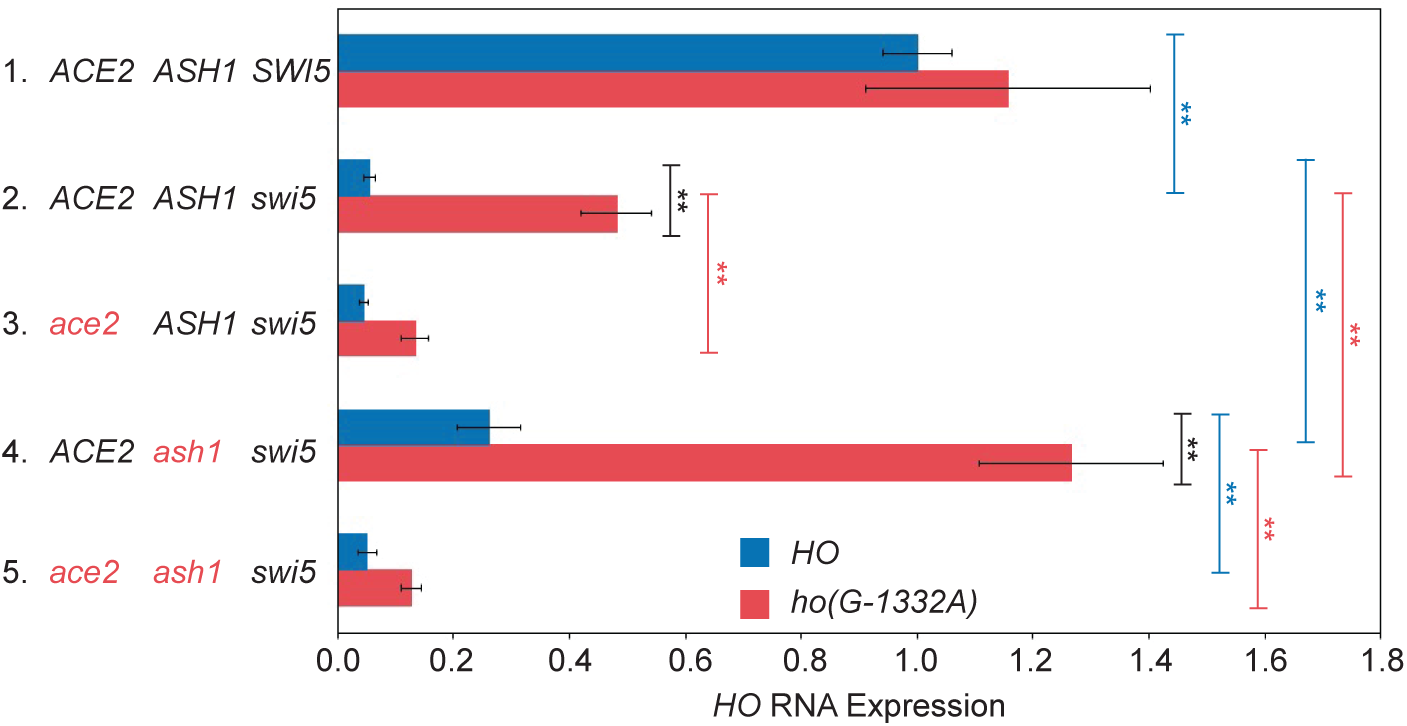
An *ho(G-1332A)* mutant and an *ash1* null additively suppress *swi5* for endogenous *HO* expression. *HO* mRNA levels were measured, normalized to *RPR1*, and expressed relative to wild type, set at 1.0. Data for strains containing wild type *HO* (blue) and *ho(G-1332A)* (red) promoters are shown for each genotype on the left. Error bars represent the standard deviation of 6 biological replicates for each strain genotype. \*\**p* < 0.01.

### Multiple negative regulators contribute to repression of the HO gene

The *ho(G-1332A)* mutants from our first genetic screen suggested that other, Ash1-independent repressors regulate the *HO* promoter. In the context of the initial screen, it may have been difficult to identify these genes, due to the necessity of the weak Ace2 activator to overcome the abundance of Ash1 in daughter cells. We therefore repeated the screen in a *swi5 ash1* mutant. *HO-ADE2* expression in *the swi5 ash1* strain is still much weaker than in wild type (Fig 1C, Lines 1-3 *ACE2*), which allowed us to observe an additive increase in *HO* expression when another negative regulator was removed. From this screen, 152 mutants were obtained, 22 of which remained after further testing. Mutations in a number of repressive chromatin/transcription factors were identified, including Isw2, Itc1, Hda1, Hda3, Rpd3, and Tup1 (Table 1, *swi5 ash1* screen). Ten mutants were in the Isw2 ATP-dependent chromatin remodeling complex, either in the catalytic Isw2 itself or in the Itc1 subunit. Isw2 associates with a large number of sites genome-wide, both via direct recruitment through transcription factors and indirect looping to ectopic sites, and Isw2 has been implicated in repression of many genes (Fazzio *et al*. 2001; Whitehouse *et al*. 2007; Yadon and Tsukiyama 2013). The Isw2 complex appears to localize near NDRs and promotes the movement of nucleosomes into the NDR to repress transcriptional initiation; it also plays a role in repression of non-coding RNAs (Whitehouse *et al*. 2007; Yadon *et al*. 2010). Five mutants were components of histone deacetylase complexes, four from the Hda1 complex (either Hda1 or Hda3) and one in Rpd3. These HDACs are recruited specifically to individual promoters via DNA-binding proteins (e.g. Rpd3 to *HO* via Ash1), and they also act globally in a more non-specific repressive role (Kurdistani and Grunstein 2003). A point mutation in the Tup1 co-repressor protein was also identified in the screen. Tup1 is recruited to a wide variety of promoters in yeast via specific DNA-binding proteins and is thought to inhibit expression by masking the activation domain of transcription factors that recruit SWI/SNF and other coactivators (Smith and Johnson 2000; Malave and Dent 2006; Wong and Struhl 2011).

The mutants identified in this screen displayed substantially stronger growth on -Ade media than *swi5 ash1* alone, similar to the effects of *swi5 ho(G-1332A)* (Figs 1C, Line 4 *ACE2* and 1D, Lines 3-6 *ACE2*). Suppression was Ace2-dependent for most mutants, but *rpd3(Y347N)* showed weak growth even in the absence of Ace2 (Fig 1D, Lines 3-6 *ace2*). This is not surprising, given that Rpd3 is an important component of repression of URS2 at the *HO* promoter, and relief of this repression is not expected to be Ace2-dependent. There were likely a number of additional *rpd3* mutants initially identified in the screen that were eliminated due to their Ace2-independent effect on repression at URS2. The particular *rpd3* point mutant that showed some Ace2-dependence may therefore have an effect on Rpd3 repression at both URS1 and URS2. The observation that *rpd3(Y347N)* is additive with *ash1* for suppression of *swi5* suggests there could be an additional factor that recruits Rpd3 to URS1.

Two surprising mutants were also identified in the screen, *cks1* and *sfp1* (Table 1, *swi5 ash1* screen). The connection between Cks1 and Sfp1 and repression of *HO* transcription is not clear, and it may be indirect by virtue of their effect on other genes that regulate *HO*. Cks1 is a small, regulatory phospho-adaptor protein involved in cell cycle regulation that associates with the cyclin-dependent kinase (the *S. cerevisiae* equivalent of Cdk1) and modulates its activity (Hadwiger *et al*. 1989; Tang and Reed 1993; Reynard *et al*. 2000; Mcgrath *et al*. 2013). In addition, Cks1 associates with the Paf1 complex, serving as an adaptor to allow recruitment of the proteasome during induction of specific genes, including *GAL1* (Pan *et al*. 2013). The 19S proteasome plays a role in evicting nucleosomes from chromatin during the course of *GAL1* induction (Chaves *et al*. 2010). It is possible that Cks1 could play a similar role during the activation of *HO* expression. Alternatively, this *cks1* mutation could alter the cell cycle and thus influence *HO* expression, or it could affect the expression of factors that regulate *HO*.

Sfp1 is an activator of ribosomal protein (RP) and ribosome biogenesis (Ribi) genes and is one of several proteins activated through the TOR pathway that determine cell size and regulate the cellular response to nutrients and stress (Jorgensen *et al*. 2002; Fingerman *et al*. 2003; Jorgensen *et al*. 2004; Marion *et al*. 2004). A recent study demonstrated that Sfp1 also acts as a glucose-regulated repressor at G1/S genes and associates with these promoters via an interaction with Swi4 (Albert *et al*. 2019). We and others were unable to detect binding of Sfp1 at the *HO* promoter in log phase cells (Benjamin Albert and David Shore, personal communication). It is possible that Sfp1 binding is transient during the cell cycle, making detection of binding in log phase cells difficult. Alternatively, the effect of *sfp1* mutants on *HO* expression could be indirect, due to changes in the cell cycle, regulation of factors that influence *HO* expression, or through its ability to influence Sch9, another protein that responds to TOR and phosphorylates Dot6 (Loewith and Hall 2011), a DNA-binding repressor that we later found to be involved in *HO* regulation (see below).

We constructed strains with disruptions for many of the genes that were identified from the *swi5 ash1* screen, and then examined the effect of the null mutations on expression of the endogenous *HO* gene in a *swi5 ash1* mutant. Deletions of *hda1*, *hda3* and *isw2* had a very modest ability to suppress *swi5 ash1*, allowing a small but reproducible increase (1.5 to 2-fold) in *HO* expression (Table 2). Here, the observed effect on *HO-ADE2* expression seems to be greater than that on endogenous *HO* expression, as the growth on -Ade appeared to increase more substantially. One possibility is that there is a non-linear correlation between growth on -Ade and the actual level of *HO-ADE2* expression. Alternatively, these mutants are global regulators of chromatin in yeast and as such, affect the expression of many genes. Thus it is difficult to interpret the effects on *HO* expression if these mutants also affect other regulators of *HO* expression. These pleiotropic effects could dampen the observed effect on *HO* expression, and the effects could be different in observing a growth phenotype versus direct measurement of mRNA levels. Finally, it is possible that the alleles isolated in the screen are not nulls, and thus could show more specific and distinct effects than a null (this was true for *tup1*; see below). Though mutations affecting Hda1 and Isw2 did not show strong effects on *HO* RNA, the modest suppression is consistent with these factors contributing in some way to repression of *HO*. Both have been linked to the Tup1 co-repressor, and it is possible that their effect at *HO* is mediated through Tup1 (Wu *et al*. 2001; Green and Johnson 2004; Zhang and Reese 2004; Fleming *et al*. 2014). Suppression by an *rpd3* null was stronger than that observed for the other chromatin modulators (3-fold above the *swi5 ash1* strain), but, as with the *rpd3* allele isolated in the screen, was only partially dependent on Ace2, likely due to its effects on both Swi5/Ace2-mediated activation at the URS1 region of the promoter and on SBF at the URS2 region.

We attempted to assess the effects of Cks1 and Sfp1 on *HO* expression using null mutations. An *sfp1* null caused changes in expression of a wide variety of genes, making it too difficult to determine an appropriate normalization control for RT-qPCR. The *cks1* null strain was extremely slow growing, and RNA results were inconsistent between experiments. We therefore used Anchor Away as an alternative to measure *HO* transcription while transiently depleting the nucleus of Cks1 (Haruki *et al*. 2008). Using this strategy, we found that loss of Cks1 did increase *HO* expression in a *swi5* mutant, but this effect was not dependent on Ace2 (Table 2). This could suggest a generic role for Cks1 and the proteasome in eviction of nucleosomes at URS2.

We found that *HO* expression was very low in *tup1* null strains, even with WT *SWI5*. Flow cytometry revealed that both the *tup1* null and an Anchor Away depletion caused a delay in G1 prior to the advent of *HO* expression (data not shown), making measurement of *HO* transcription uninterpretable. We therefore used the point mutant isolated in the screen, an H575Y substitution, for endogenous *HO* RNA analysis. The *tup1(H575Y)* mutant displays normal growth, suggesting some critical targets of Tup1 are not affected by this mutation. However, *tup1(H575Y)* increased suppression of *swi5* additively with *ash1*, at a level exceeding that of the null mutants for *hda1*, *hda3* and *isw2* (Table 2; Fig 3, compare Lines 4 and 5). Expression of *HO* in the *tup1(H575Y) ash1 swi5* mutant was Ace2 dependent (Fig 3, compare Lines 5 and 6).

**Figure 3.**
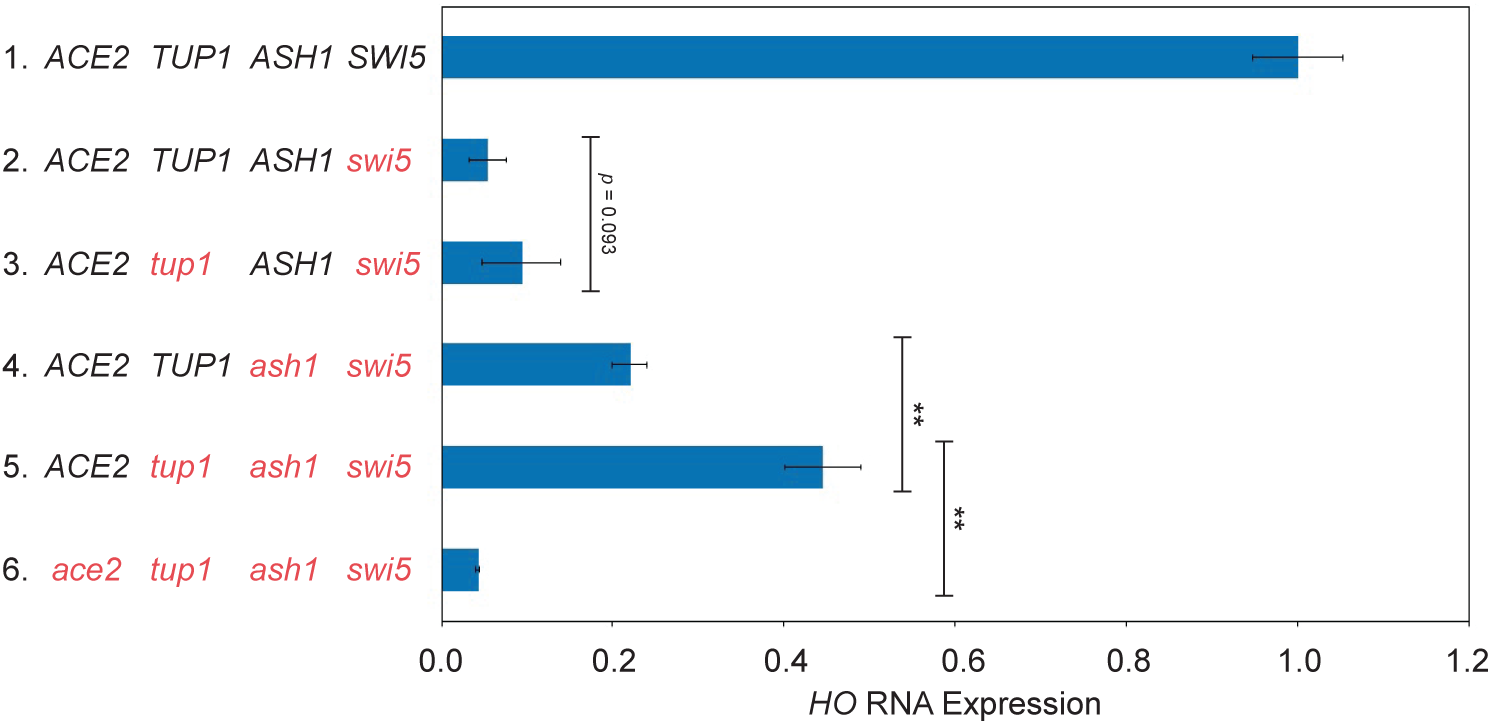
The *tup1(H575Y)* point mutant suppresses an *ash1 swi5* mutant for endogenous *HO* expression. *HO* mRNA levels were measured, normalized to *RPR1*, and expressed relative to wild type. Genotypes are indicated at the left. Error bars represent the standard deviation of 3 biological replicates for each strain. \*\**p* < 0.01.

### Additional HO promoter mutants suggest the presence of binding sites for negative regulators

Three *HO* promoter mutants were also identified from the *swi5 ash1* screen, two changes at −1332 (the position identified in the *swi5* screen) and one at −1358 (Table 3). Since these genome-wide screens may not have been saturated for *HO* promoter mutations, we next performed directed screens in which we mutagenized the region of the *HO* promoter from −1826 to −268, encompassing both URS1 and URS2, and integrated the mutant versions into a complete *HO-ADE2* reporter. Many additional *HO* promoter mutants were obtained (Table 3). All mutations were in the URS1 region of the promoter, which was not surprising, given that we were selecting for mutants that were dependent upon the Ace2 activator and therefore required relief of repression in URS1. A total of seven additional independent *ho(G-1332A)* mutants were identified, highlighting the importance of −1332 in *HO* regulation. Other mutations were identified in nucleotides in close proximity to *ho(G-1332A)*, further suggesting the presence of a binding site for a factor at this position.

Listing the mutants by position within the promoter revealed not only the cluster of mutations encompassing *ho(G-1332A)* (Cluster 2; containing mutations at positions - 1338 to −1328) but two additional clusters, Cluster 1 (with mutations from −1416 to - 1399) and Cluster 3 (with mutations from −1144 to −1138) (Figure 4; Table 3). Only a few mutations fell outside these clusters (positions −1762, −1734, and −1442), and several sites were identified multiple times, suggesting the screen was near saturation. Isolates with multiple mutations were either compared to other hits from the screen or tested further as single mutations to determine the likely cause of suppression (Tables 3, 4). Cluster 1 contains mutations at five positions in the *HO* promoter, from −1416 to −1399, where red indicates evolutionarily conserved nucleotides (Fig 4). We searched for known recognition motifs for DNA binding proteins within this region and tested candidate proteins by ChIP analysis but were unable to identify a protein disrupted by these mutations. Clusters 2 and 3 encompass known regulatory motifs, as detailed below.

**Figure 4.**
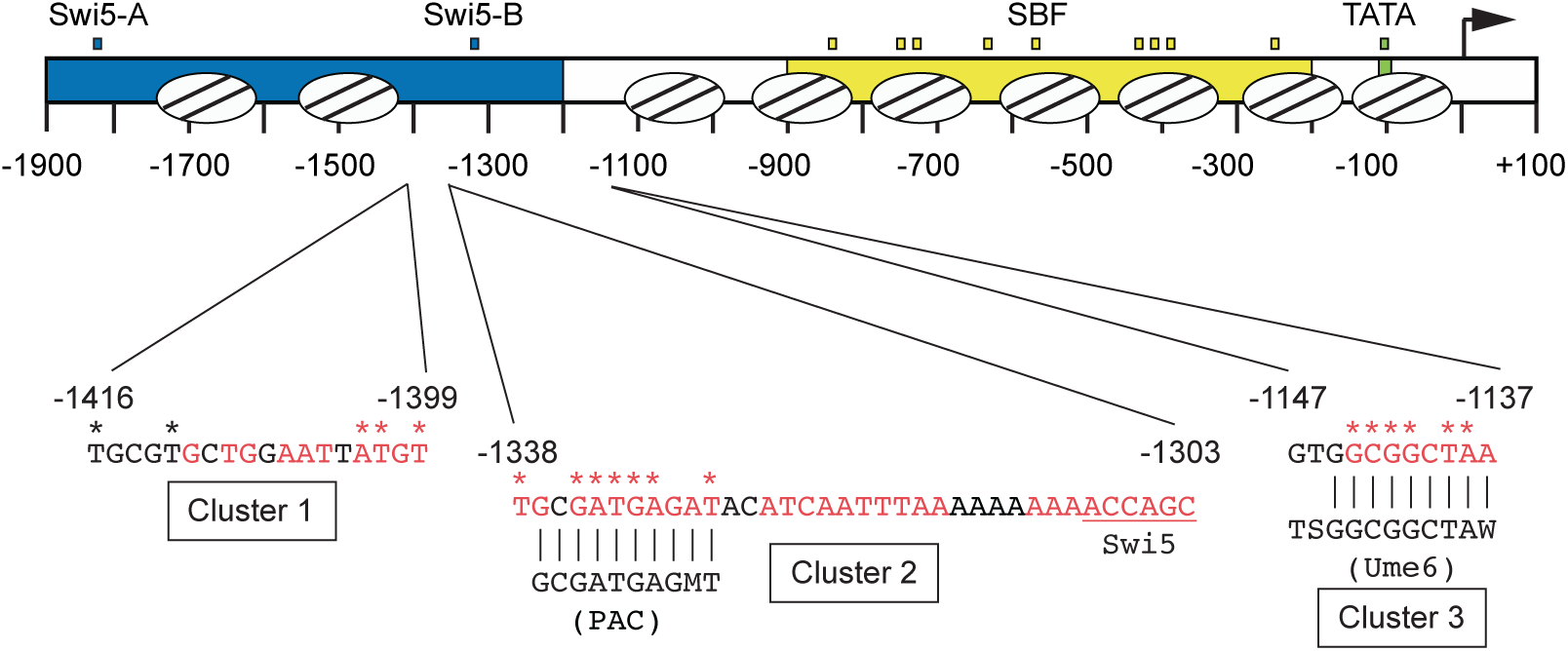
Clusters of mutations within the *HO* promoter that suppress *swi5 ash1*. A schematic of 1900 nucleotides of the *HO* promoter is shown, with nucleosome positions indicated by ovals with lines. URS1 (blue), URS2 (yellow) and TATA (green) are indicated. Small blue boxes = Swi5 binding sites; Small yellow boxes = SBF binding sites; Small green box = TATA. The arrow indicates the position of the ATG (+1). Nucleotide sequences are shown for three clusters of mutations identified in screens for *HO* mutants that allow activation of *HO-ADE2* in a *swi5* or *swi5 ash1* mutant. Red nucleotides are conserved across species of *Saccharomyces*, and stars indicate the positions of the mutations. Cluster 2 contains a PAC motif and Cluster 3 contains a Ume6 binding site, with the sequence of the consensus site shown for each. Vertical lines indicate positions of *HO* sequence that match the PAC or Ume6 consensus sites. The sequence downstream of Cluster 2 was extended to show the relative position of the −1300 Swi5 binding site, underlined in red.

Strains were constructed with individual *HO* promoter mutations in the native *HO* context, and RNA expression was measured. These *cis* mutations had a stronger effect on *HO* expression than the *trans* mutants identified in the previous screens, with the promoter mutations generally causing a 2 to 6-fold increase in expression compared to the *swi5 ash1* control (Table 4). Additionally, these *HO* promoter mutations were mostly Ace2-dependent. We speculate that off-target pleiotropic effects caused by eliminating important chromatin/transcription regulatory factors diminished the specific effect of these mutations at the *HO* locus, while promoter mutations had only local effects.

### The HO promoter contains a PAC motif that is bound by the Dot6 protein

Mutations within Cluster 2 were the most frequently identified in our screens and displayed the strongest effects on *HO* expression in a *swi5 ash1* mutant (Fig 4; Table 4). The altered nucleotides in Cluster 2 mutants are all conserved among species of *Saccharomyces*. Cluster 2 includes a perfect match to a polymerase A/C (PAC) motif (GCGATGAGMT), and 7 of our mutations are in or adjacent to this motif. PAC sites were first identified within promoters of ribosome biogenesis (Ribi) genes, a group of approximately 300 genes that encode various factors required for ribosome assembly and translation (Hughes *et al*. 2000; Jorgensen *et al*. 2004; Wade *et al*. 2006). Two paralogous repressive factors, Dot6 and Tod6, bind to the PAC sites in Ribi genes and reduce transcription via recruitment of the Rpd3 complex (Huber *et al*. 2011).

Association of Dot6/Tod6 with Ribi promoters is modulated via phosphorylation by Sch9 in response to environmental conditions through the TOR and PKA pathways (Lippman and Broach 2009; Huber *et al*. 2011). A number of non-Ribi genes also have PAC sites within their promoters, raising the possibility that Dot6 and/or Tod6 may be general factors that are employed to repress other genes in different contexts (Lippman and Broach 2009). Since the *HO* promoter has a perfect match to the PAC motif, and the strongest mutations we identified are within the core PAC site, we investigated whether Dot6 and/or Tod6 might play a role in regulation of *HO* expression.

We constructed a *dot6 tod6 swi5 ash1* strain to determine whether loss of Dot6 and Tod6 could increase *HO* expression above the level observed in a *swi5 ash1* mutant. Expression of *HO* in the *dot6 tod6 swi5 ash1* mutant was approximately 30% of wild type, an increase of 1.5 to 1.7-fold above that of *swi5 ash1* (Table 2; Fig 5A, compare Lines 3 and 4). This is a very modest effect, although similar to what was observed with *hda1*, *hda3* and *isw2*, and could suggest that like these other global transcription and chromatin factors, null mutations in *DOT6* and *TOD6* do not recapitulate the effect of an *HO* promoter mutation, possibly due to additional global effects. While the RNA results are not striking, they are consistent with the possibility that Dot6/Tod6 may bind to *HO* and influence its expression.

**Figure 5.**
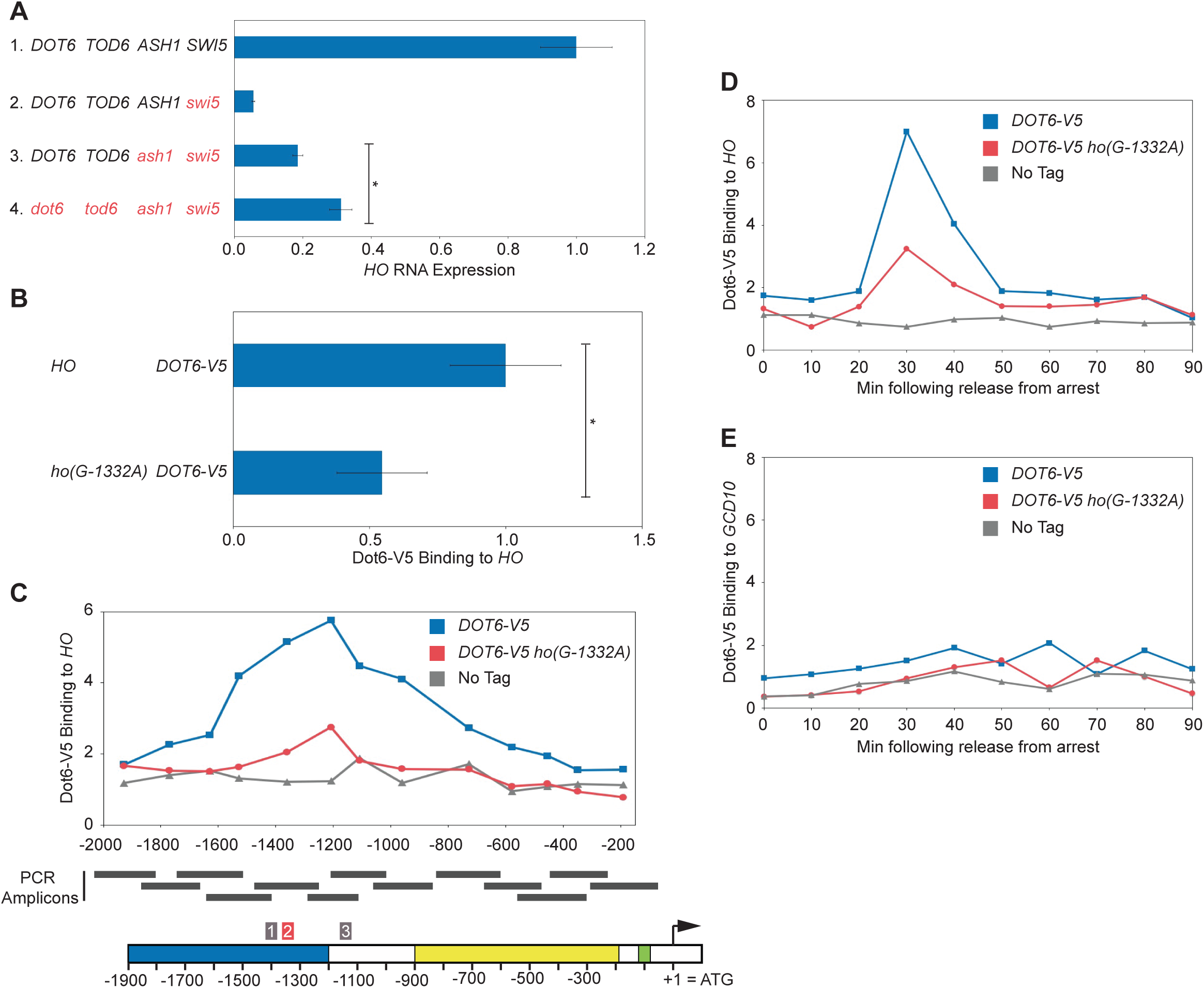
Dot6 associates with the PAC motif in the *HO* promoter and may influence *HO* expression. (A) *HO* mRNA levels were measured, normalized to *RPR1*, and expressed relative to wild type. Genotypes are indicated at the left. Error bars represent the standard deviation of 3 to 4 biological replicates for each strain. \**p* < 0.05. (B) Binding of Dot6-V5 to HO wild type and *ho(G-1332A)* promoters from heat shocked cells was determined by ChIP, followed by qPCR with primers that amplify *HO* sequence from −1471 to −1250. Dot6-V5 enrichment at *HO* was normalized to enrichment at the *GCD10* promoter and to the corresponding input sample and then graphed relative to wild type. Error bars show the standard deviation of 3 biological replicates. \**p* < 0.05. (C) Single samples of *HO* wild type (blue) and *ho(G-1332A)* (red) were chosen from (B) and used for qPCR with primers that span the length of URS1 and URS2 in approximately 150 bp increments. Positions of the PCR amplicons are indicated by the gray bars. Points on the graph correspond to the midpoints of these amplicons, with the x-axis showing position across the *HO* promoter. A schematic of the *HO* promoter is shown at the bottom, with positions of Clusters 1, 2, and 3 indicated. Cluster 2 contains the PAC motif and is highlighted red. For each primer set, Dot6 enrichment was normalized to an intergenic region on chromosome I (*IGR-I*) and to input. A “No Tag” control is shown to indicate the background level of binding (gray). (D) Binding of Dot6-V5 to the *HO* wild type (blue) and *ho(G-1332A)* (red) promoters was measured in cells containing the *GALp::CDC20* allele that had been synchronized by galactose withdrawal and re-addition. The 0 min time point represents the G2/M arrest, before release with galactose addition. Cells were harvested at the indicated time points following release (x-axis), and samples were processed for ChIP analysis. Dot6 enrichment at the *HO* promoter was measured using primers that span from −1295 to - 1121 and was normalized to *IGR-I* and to input. Note that *HO* expression begins to rise at approximately 30 min after release, and peaks at 50 min (see Figure 7). (E) Same as (D), using primers to the *GCD10* promoter, normalized to *IGR-I* and to input.

To determine whether Dot6 and/or Tod6 associate with the *HO* promoter, we performed ChIP analysis using Dot6-V5 and Tod6-V5 strains both in normal growth conditions and in conditions of heat shock stress. The heat shock conditions allowed us to use a Ribi gene as a positive control to demonstrate that the ChIP was successful. We were unable to detect a reliable ChIP signal for either Tod6-V5 or Tod6-HA under normal growth conditions, either at Ribi targets or at *HO*. However, we detected Dot6-V5 bound to multiple Ribi targets as well as to *HO* under heat shock conditions (Fig 5B; Figs S1A,B). Using primers that span the URS1 and URS2 regions of the *HO* promoter in approximately 150 bp increments, we observed a broad peak of Dot6 binding centered over the region of the promoter that contains the PAC site (Fig 5C). This binding was significantly diminished, but not completely eliminated, by the *ho(G-1332A)* mutation (Figs 5, S1), suggesting that Dot6 is capable of binding to *HO* at the PAC site and that the *ho(G-1332A)* mutation affects its binding. However, we were unable to reproducibly detect binding of Dot6 to *HO* under the conditions in which we typically monitor *HO* expression, i.e. non-heat shock stress. This suggests that Dot6 may bind to *HO* only transiently at a specific point in the cell cycle or that Dot6 does not tightly associate with the promoter and can therefore only reliably be detected when Dot6 has an increased capacity to bind chromatin during stressful conditions.

We performed Dot6-V5 ChIP in cells synchronized using a *GALp::CDC20* arrest and release protocol to investigate the possibility that Dot6 binds to *HO* at a specific time in the cell cycle. This type of cell synchrony has been used extensively to monitor sequential factor binding and expression at the *HO* promoter during the cell cycle (Bhoite *et al*. 2001; Takahata *et al*. 2009b). The Swi5 transcription factor binds at 20 min following release from G2/M arrest, and *HO* expression occurs at 50 min. We observed a strong peak of Dot6 binding at 30 min following release from the arrest, a time that is coincident with binding of the Ash1 repressor (Fig 5D; see also Fig 7). Similar to the ChIP results in heat shock conditions, the *ho(G-1332A)* mutation diminished but did not eliminate binding of Dot6. Importantly, a peak was not observed during the cell cycle time course at the Ribi target genes we tested (Fig 5E; the *GCD10* promoter is shown as an example). These results support the idea that Dot6 can bind to the *HO* promoter in the absence of an environmental stress and that it could be the factor responsible for the change in *HO* expression observed in the *ho(G-1332A)* mutant. It remains an open possibility that Tod6 could also associate with *HO* and influence its expression. The fact that we did not obtain mutations in *DOT6* as part of either screen may suggest that both *DOT6* and *TOD6* need to be mutated in order to observe an effect on *HO-ADE2* expression.

**Figure 6.**
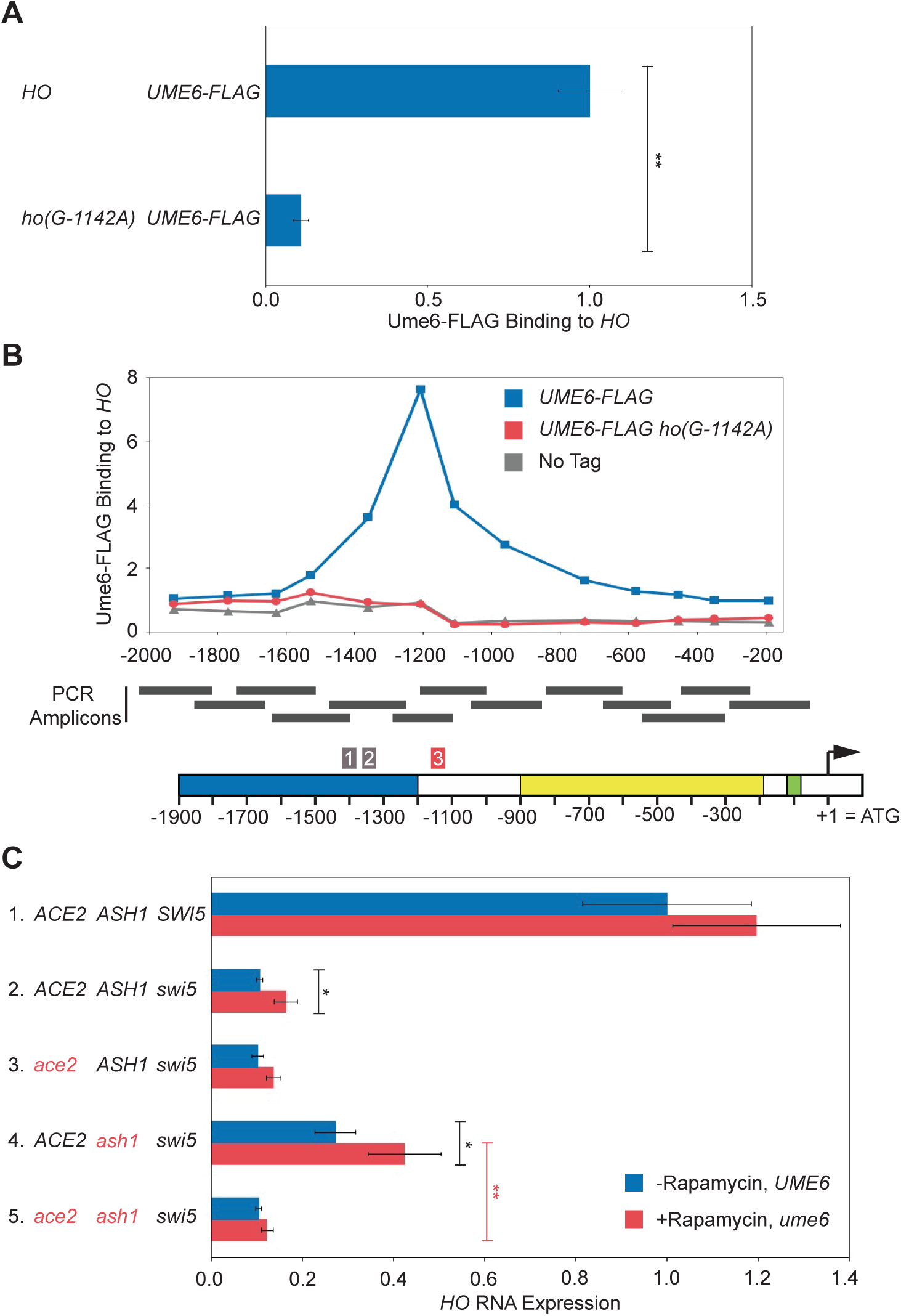
Ume6 associates with its consensus motif in the *HO* promoter and influences *HO* expression. (A) Binding of Ume6-FLAG to *HO* wild type and *ho(G-1142A)* promoters was determined by ChIP, followed by qPCR with primers that amplify *HO* sequence from - 1295 to −1121. Ume6-FLAG enrichment for each sample was normalized to enrichment at the promoter of *INO1* and to the corresponding input sample and then graphed relative to wild type. Error bars show the standard deviation of 3 biological replicates. \*\**p* < 0.01. (B) Single samples of *HO* wild type (blue) and *ho(G-1142A)* (red) were chosen from (A) and used for qPCR with primers that span the length of URS1 and URS2 in approximately 150 bp increments. Positions of the PCR amplicons are indicated by the gray bars. Points on the graph correspond to the midpoints of these amplicons, with the x-axis showing position across the *HO* promoter. A schematic of the *HO* promoter is shown at the bottom, with positions of Clusters 1, 2, and 3 indicated. Cluster 3 contains the Ume6 consensus motif and is highlighted red. For each primer set, Ume6 enrichment was normalized to an intergenic region on chromosome I (*IGR-I*) and to input. A “No Tag” control is shown to indicate the background level of binding (gray). (C) *HO* mRNA levels were measured, normalized to *RPR1*, and expressed relative to wild type. Error bars represent the standard deviation of at least 3 biological replicates for each strain. See Materials and Methods section for a full description of genotypes and growth conditions. Cultures were split into two, and one half was untreated (-Rapamycin, *UME6*; blue bars), while the other half was treated with 1μg/ml rapamycin for 2 hours (+Rapamycin, *ume6*; red bars). \**p* < 0.05, \*\**p* < 0.01.

**Figure 7.**
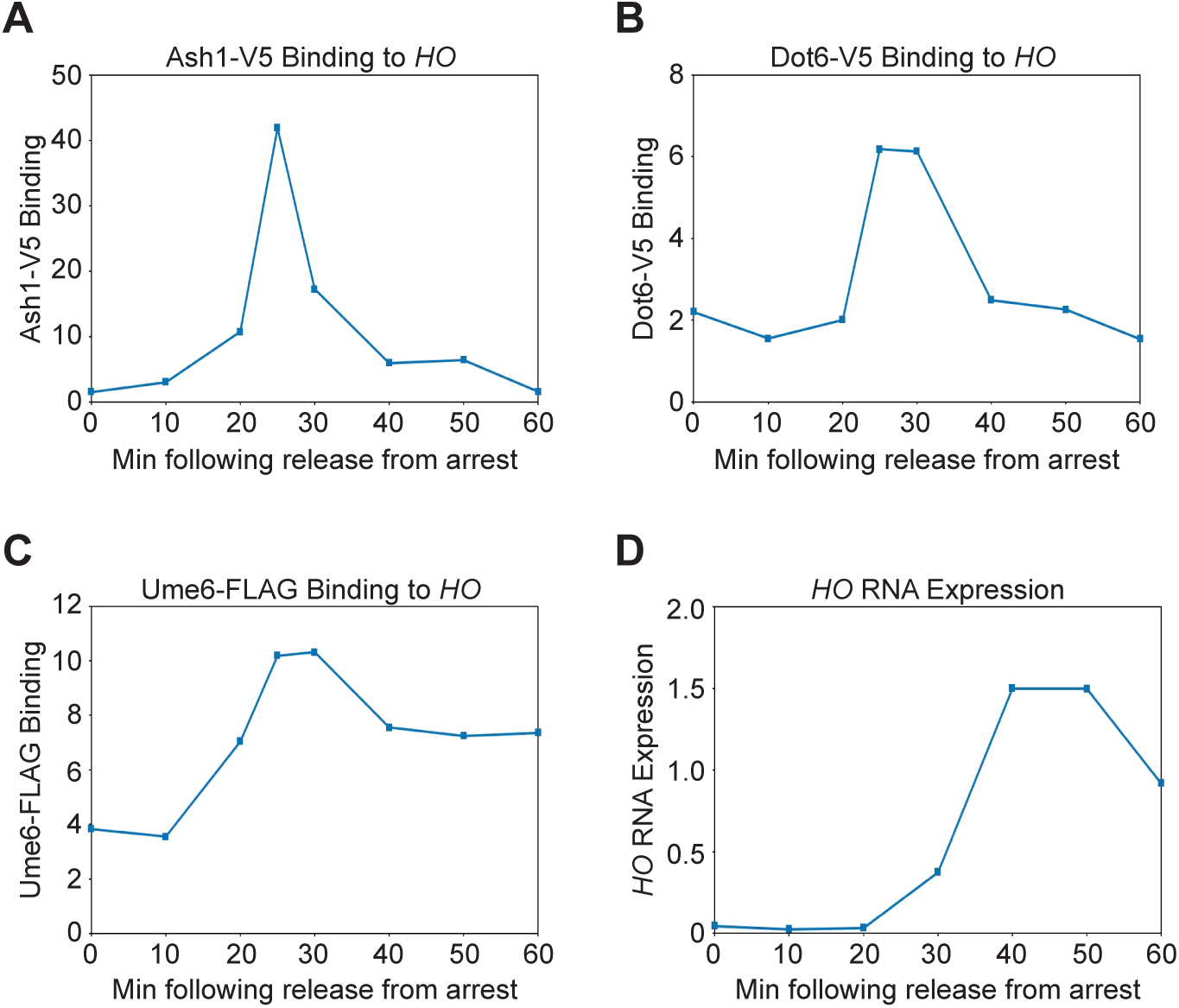
The Ash1, Dot6 and Ume6 repressors associate with the *HO* promoter before the peak of HO expression. (A, B, C) Binding of Ash1-V5 (A), Dot6-V5 (B), Ume6-FLAG (C) to the *HO* promoter during a cell cycle arrest and release experiment. Cells containing the *GALp::CDC20* allele were synchronized by galactose withdrawal and re-addition. The 0 min time point represents the G2/M arrest, before release by addition of galactose. Cells were harvested at the indicated time points following release (x-axis), and samples were processed for ChIP analysis. Enrichment at the *HO* promoter was measured using primers that span from −1295 to −1121 and was normalized to *IGR-I* and to input. Graphs show one representative experiment from each strain for comparison. Supplemental Figure S4 shows three replicates for each strain. (D) A representative example of *HO* mRNA expression measured over the course of the synchrony experiment and normalized to *RPR1* (using a Dot6-V5 strain).

### Ume6 binds to the HO promoter and negatively regulates its expression

Cluster 3 mutations (−1144 to −1138) also have strong effects on *HO* expression (Table 4). This region of the promoter contains a match to the consensus Ume6 binding site (Park *et al*. 1992; Williams *et al*. 2002; Doniger *et al*. 2005). Ume6 is a DNA-binding factor that was originally identified as a repressor of meiotic genes (Strich *et al*. 1994). Significantly, this screen also identified *SIN3* and *RPD3*, which repress *HO* expression (Dorland *et al*. 2000). Ume6 localizes to many sites genome-wide and recruits both the Rpd3 histone deacetylase complex and the Isw2 chromatin remodeling complex (Kadosh and Struhl 1997; Rundlett *et al*. 1998; Goldmark *et al*. 2000). We examined whether Ume6 binds to the *HO* promoter using a Ume6-FLAG tagged strain. Strong binding of Ume6 to the *HO* promoter was observed, and this binding was eliminated by the *ho(G-1142A)* mutation, one of the mutations identified in the screen (Figs 6A, S2A). Examination of binding across the *HO* promoter revealed a very distinct peak of Ume6 binding centered at −1140, as expected based on the position of the Ume6 consensus site (Fig 6B).

No mutations in the *UME6* gene were identified in either of our suppressor screens. However, in both screens, we eliminated a number of Ade+ mutants whose Ade+ phenotype dissipated with continued passage, and we now suggest these strains could have had a mutation in *UME6*. Yeast cells with a *ume6* mutation have a strong growth defect, and in many cases this defect can be suppressed by chromosome duplication (Fazzio *et al*. 2001). Hence, *ume6* mutants could have been initially Ade+ due to the loss of a negative regulator that inhibits *HO-ADE2* transcription, but then became Ade-due to the aneuploidy. Many of the mutants we eliminated also displayed multiple colony sizes and variable Ade+ phenotypes, consistent with some type of genomic instability.

To test this, we disrupted *UME6* in diploid cells and then sporulated to isolate fresh haploid *ume6* mutants. However, it was problematic to directly examine the effect of a *ume6* null on *HO* expression, as we had inconsistent results for biological replicates within an experiment and between experiments. There was also variation in colony size in the haploid *ume6* strains, suggesting rapid aneuploidy. We therefore used Anchor Away as an alternate strategy to deplete the nucleus of Ume6. *INO1*, a known Ume6 target gene (Slekar and Henry 1995; Jackson and Lopes 1996), displayed greatly increased expression after Ume6 was depleted by addition of rapamycin to the medium (Fig S3). Under these conditions, we also observed suppression of both *swi5* and *swi5 ash1* for *HO* expression (Fig 6C; compare Lines 2 red vs. blue and 4 red vs. blue). Consistent with the effects of other global negative regulators on *HO* expression, the 1.6-fold increase in transcription was very modest but reproducible. Importantly, the effect of Ume6 depletion in the *ash1 swi5* strain was Ace2-dependent (compare Lines 4 and 5 red). We conclude that Ume6 is an important repressor of *HO* expression, but binding site mutations more accurately demonstrate its significance than loss of the protein, which causes global effects on transcription.

### Ash1, Dot6 and Ume6 associate with the HO promoter largely independently, but with similar timing

Our screens identified three DNA-binding repressor proteins that influence *HO* expression: Ash1, Dot6, and Ume6. All of these factors are sub-stoichiometric members of the Rpd3 complex and have been demonstrated to recruit Rpd3 to target genes. At the *HO* promoter, Ash1, Dot6 and Ume6 could be redundant with one another, such that loss of one would not have a substantial effect on expression. Alternatively, they could all be necessary for complete repression, either acting independently or in an interdependent fashion.

To begin to determine whether Ash1, Dot6 and Ume6 play similar or distinct roles in repression at *HO*, we examined the timing of binding of each protein to the *HO* promoter in cells synchronized by *GALp::CDC20* arrest and release. All three proteins bound to *HO* at approximately 25 to 30 min following release from the arrest, a time point following Swi5 activator binding (20 min) but preceding *HO* expression (50 min) (Fig 7). Ash1 and Dot6 binding peaked at 25 min and then diminished back to their original levels before *HO* expression started. While Ume6 associated with the promoter at approximately the same time as Ash1 and Dot6, binding did not diminish as much as for Ash1 and Dot6, and significant amounts of Ume6 remained bound beyond the peak of *HO* expression.

The similar timing of association of Ash1, Dot6 and Ume6 with the *HO* promoter in synchronized cells suggested the possibility that one or more of these factors might be inter-dependent upon one another for binding. We therefore examined binding of each protein in the absence of the other two factors by ChIP analysis. We used an *ash1* null to examine the effect of loss of Ash1 and promoter mutants *ho(G-1332A)* and *ho(G-1142A)* to cause the specific reduction or loss of Dot6 and Ume6, respectively, at the *HO* promoter.

Ash1-V5 association with the *HO* promoter did not decrease in either an *ho(G-1332A)* or in an *ho(G-1142A)* mutant, suggesting that Ash1 binding does not require Dot6 or Ume6 (Fig 8A). Instead, we observed a small increase in binding of Ash1 in these mutant strains. This slight additional binding could suggest that Dot6 and Ume6 binding are very weakly inhibitory to association of Ash1. Ume6-FLAG association with the *HO* promoter did not change significantly in the *ho(G-1332A)* mutant, and was reduced modestly by an *ash1* mutation (Fig 8B). Thus Ume6 binding is completely independent of Dot6 and largely independent of Ash1.

**Figure 8.**
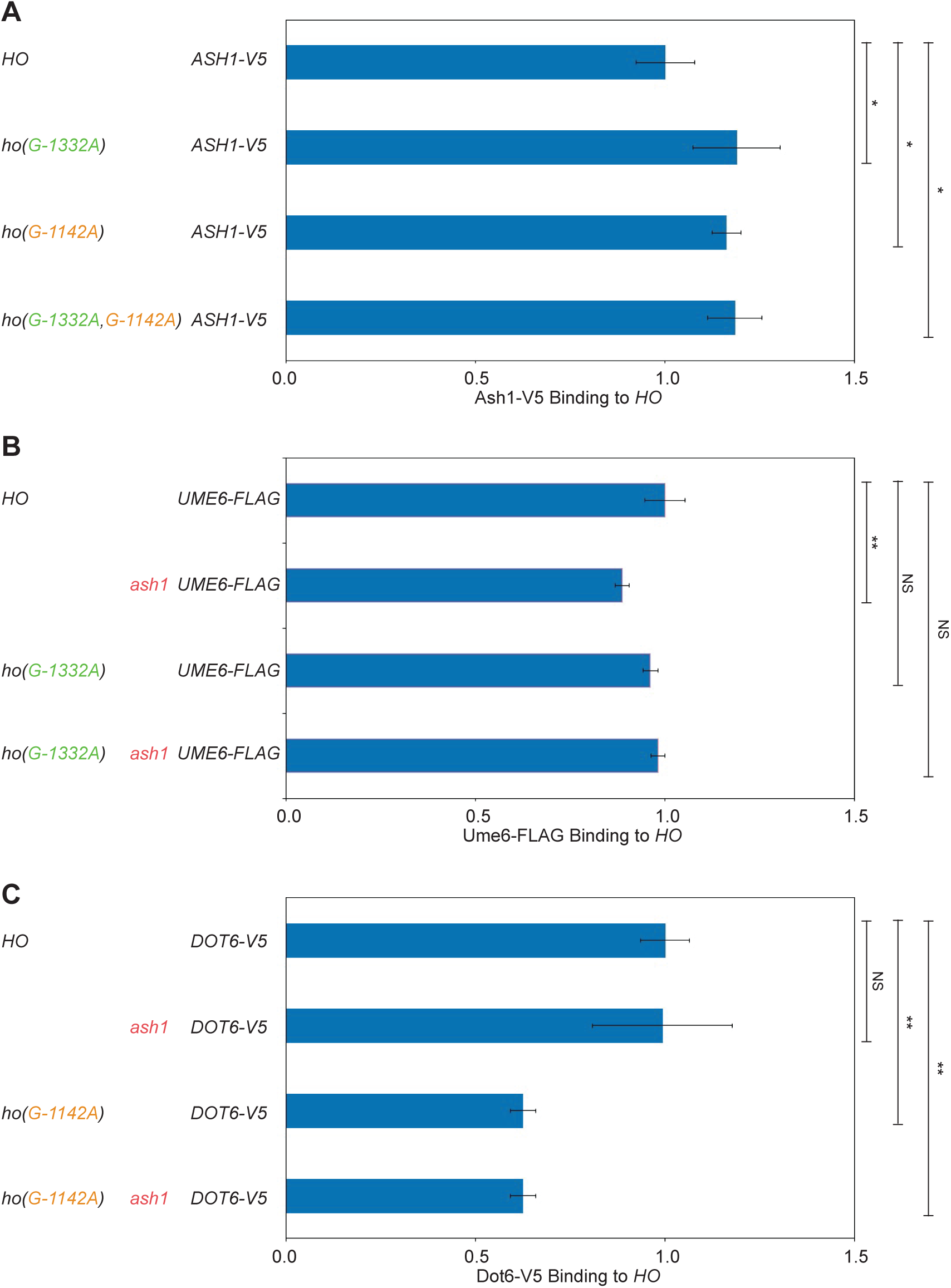
Ash1, Dot6, and Ume6 are largely independent of one another for association with the *HO* promoter. Binding of Ash1-V5, Dot6-V5 and Ume6-FLAG to the *HO* promoter in wild type and in the absence of one or both of the other factors or sites was determined by ChIP. Each sample was normalized to a reference control and to its corresponding input and graphed relative to wild type enrichment. Strain genotypes are indicated at the left. Error bars show the standard deviation of 4 biological replicates. \**p* < 0.05, \*\**p* < 0.01. (A) Ash1-V5 enrichment at *HO* −1295 to −1121 is shown normalized to binding at the promoter of the *CLN3* gene. (B) Dot6-V5 enrichment at *HO* −1295 to −1121 in heat-shocked cells is shown normalized to binding at the intergenic region *IGR-I*. (C) Ume6-FLAG enrichment at *HO* −1295 to −1121 is shown normalized to binding at the promoter of the *INO1* gene.

The enrichment of Dot6 was decreased in both the *ho(G-1142A)* and *ash1 ho(G-1142A)* mutants, suggesting that the presence of Ume6 may have some influence on the level of association of Dot6 with the *HO* promoter (Fig 8C). Some Dot6 could be brought to *HO* via Ume6 if Dot6 comes along as part of the Ume6-containing Rpd3 complex (Carrozza *et al*. 2005), and this may explain the very broad peak of Dot6 binding across the *HO* promoter (Fig 5C) in contrast to the narrower peak of Ume6 binding (Fig 6B). This possibility is supported by the observation that the *ho(G-1332A)* mutation, which reduced but did not eliminate Dot6 binding, shifted residual Dot6 binding toward the transcription start site, near the Ume6 site (Fig 5C). Similar results were observed when we mutated the entire PAC site (data not shown). Alternatively, perhaps the presence of Ume6 affects the positioning of nucleosomes in such a way that Dot6 association is altered in the absence of Ume6. This possibility could explain how Ume6 could affect Dot6 binding at its PAC site despite the nearly two hundred nucleotides between the binding sites for these two factors. Overall, these results suggest that while all three proteins are capable of some level of association with the *HO* promoter independent of the others, the presence of Ume6 does appear to affect binding of Dot6 through a mechanism that is currently not understood.

### The ash1, ho(G-1332A) and ho(G-1142A) mutations are additive for effects on HO expression but not for loss of Rpd3 recruitment

The lack of strong dependency of Ash1, Dot6, and Ume6 upon one another for binding to the *HO* promoter suggests they may each be required for full repression of *HO* expression. We therefore examined whether the absence of each of these factors was additive for their relief of *HO* repression at URS1. For these experiments, we continued to use the *ash1* null mutant, along with *ho(G-1332A)* for reduction in Dot6 (and possibly Tod6) and *ho(G-1142A)* for elimination of Ume6.

We measured *HO* expression in all combinations of *ash1*, *ho(G-1332A)*, and *ho(G-1142A)*, both in *SWI5*+ (Fig 9, Left graphs) and in *swi5* strains (Fig 9, Right graphs). All three single mutants caused an increase in *HO* expression in otherwise wild type cells (Fig 9, Lines 2,3 blue and Line 1 red). The *ash1* mutant had the strongest effect, with an approximately 2-fold increase, likely due to a gain of expression of *HO* in daughter cells (Line 1 red). The *ho(G-1332A)* (Dot6) and *ho(G-1142A)* (Ume6) mutants were additive with one another, increasing *HO* expression to 2.6-fold higher than wild type (compare Line 4 blue to Lines 2 and 3 blue), which was increased even higher in the triple *ash1 ho(G-1332A, G-1142A)* mutant (4.3-fold higher than wild type; compare Line 4 red to Line 4 blue). The *ho(G-1142A)* mutant showed significant additivity with *ash1*, while the additivity of *ho(G-1332A)* and *ash1* was less pronounced (compare Lines 2 and 3 red to Line 1 red).

**Figure 9.**
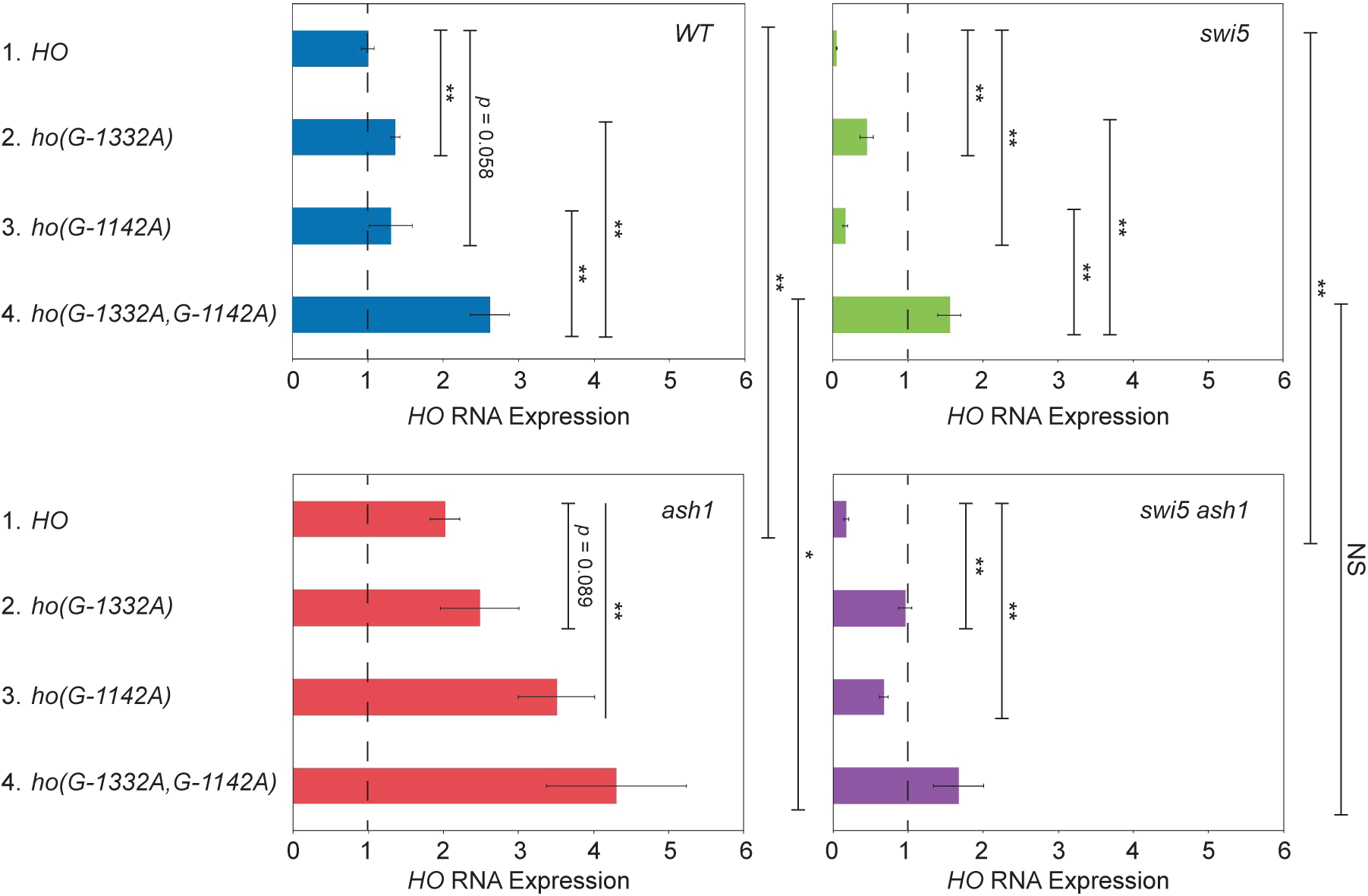
Combining the *ash1*, *ho(G-1332A)*, and *ho(G-1142A)* mutations shows additive suppression of *ash1 swi5*. *HO* mRNA levels were measured, normalized to *RPR1*, and expressed relative to wild type. Each graph shows results for the genotype indicated (WT in blue, *swi5* in green, *ash1* in red, *ash1 swi5* in purple). The type of *HO* promoter for each bar is shown at the left (*HO* wt, *ho(G-1332A)*, *ho(G-1142A)* and *ho(G-1332A, G-1142A)*. The dotted line indicates the level of *HO* expression in a *SWI5 ASH1* strain with the wild type *HO* promoter. Error bars represent the standard deviation of 4 biological replicates for each strain. \**p* < 0.05, \*\**p* < 0.01.

All three single mutants suppressed *swi5* for expression of *HO* (Fig 9, Right graphs). The strongest was *ho(G-1332A)*, which increased expression 9-fold over that of a *swi5* mutant (45% of wild type; compare Lines 1 and 2 green); *ho(G-1142A)* and *ash1* had more modest effects (16% and 18% of wild type; compare Lines 3 green and Line 1 purple to Line 1 green). Each of the *ho* promoter mutants was additive with *ash1* for suppression of *swi5* (compare Lines 2 and 3 purple to Line 1 purple). Interestingly, the *ho(G-1332A)* and *ho(G-1142A)* mutants were also strongly additive with one another for suppression of *swi5*, with a value 150% of wild type (compare Line 4 green with Lines 2 and 3 green); this value did not increase substantially with the addition of the *ash1* mutant (Line 4 purple). These results demonstrate that mutations that reduce or eliminate Ash1, Dot6 and Ume6 localization are generally additive for relief of *HO* repression, suggesting the three proteins and their associated complexes act independently. However, in the absence of the Swi5 activator, loss of Ash1 is additive with either single promoter mutant but not with the double promoter mutant. This suggests that if the proteins at the PAC site and at the Ume6 site have been reduced or removed, elimination of Ash1 has little additional effect on increasing *HO* expression in a *swi5* mutant. The same is not true for loss of Ash1 and the PAC-associated proteins or Ash1 and the Ume6 site-associated proteins; in both of these cases, removal of the third complex further relieves repression.

The additivity of the Ash1, PAC and Ume6 site mutants thus appears to be different in the presence and absence of Swi5, for reasons that are not clear. Mother and daughter cells could differ in their responses to loss of the repressors. Ash1 protein levels are vastly different in the two cell types, and the effect of no Ash1 in mother cells may be different than in daughter cells. Swi5 has the potential to activate *HO* in both mother and daughter cells, whereas in the absence of Swi5, Ace2 can activate *HO* only in daughter cells. Perhaps the additivity of all three mutants (Ash1, PAC, Ume6) is only seen in a *SWI5+* strain because Swi5 normally activates *ASH1* expression (Bobola *et al*. 1996).

Interpretation of these results is also somewhat complicated by the observation that Ume6 appears to contribute positively to Dot6 binding at the *HO* promoter (Fig 8C). The additivity of the *ho(G-1332A)* and *ho(G-1142A)* mutants could partly be explained if the amount of Dot6 association with *HO* in the *ho(G-1332A, G-1142A)* double mutant is lower than in the *ho(G-1332A)* mutant alone. However, Ume6 binding becomes undetectable in the *ho(G-1142A)* mutant, while Dot6 binding is simply reduced, suggesting that the effect of *ho(G-1142A)* is likely mostly due to the absence of Ume6.

The additivity of *ash1*, *ho(G-1332A)* and *ho(G-1142A)* mutants for relief of *HO* repression suggests that Ash1, Dot6, and Ume6 may independently recruit the Rpd3 complex to contribute to repression. To examine this possibility, we performed Rpd3-V5 ChIP in strains with single mutants and mutant combinations to determine whether recruitment of Rpd3 by these factors is redundant or additive (Figs 10, S5A). At the URS2 region of *HO*, SBF recruits Whi5 and Stb1, which in turn recruit Rpd3 (Takahata *et al*. 2009a), raising the possibility that Rpd3 bound at URS2 could affect the results of our Rpd3 ChIP assay at URS1. To circumvent this problem, we used strains with mutations that disrupt binding at all nine SBF sites within URS2. Both *ash1* and *ho(G-1142A)* single mutants diminished recruitment of Rpd3 at URS1, but neither alone eliminated binding (compare Lines 2 and 4 to Line 1). When combined, *ash1* and *ho(G-1142A)* reduced Rpd3 binding further (compare Line 6 to Lines 2 and 4), suggesting Ash1 and Ume6 recruit Rpd3 independently, with each contributing separately to its full recruitment and repression. In contrast, the *ho(G-1332A)* mutant did not appear to substantially affect Rpd3 binding either on its own or in combination with *ash1*, *ho(G-1142A)* or both (compare Lines 3 vs. 1, 5 vs. 2, 7 vs. 4, 8 vs. 6). Thus, Ash1 and Ume6 are important for recruitment of Rpd3 to URS1 of the *HO* promoter, but surprisingly, Dot6 does not appear to contribute in a significant way. The observation that *ho(G-1142A)* and *ho(G-1332A)* do not have the same effect on Rpd3 binding at *HO* supports the suggestion that the effect of *ho(G-1142A)* on *HO* expression (Fig 9) is not mediated by the partial loss of Dot6 but rather by the substantial loss of Ume6 binding. If the predominant effect of both mutations was partial loss of Dot6, both should display the same consequence for Rpd3 binding.

**Figure 10.**
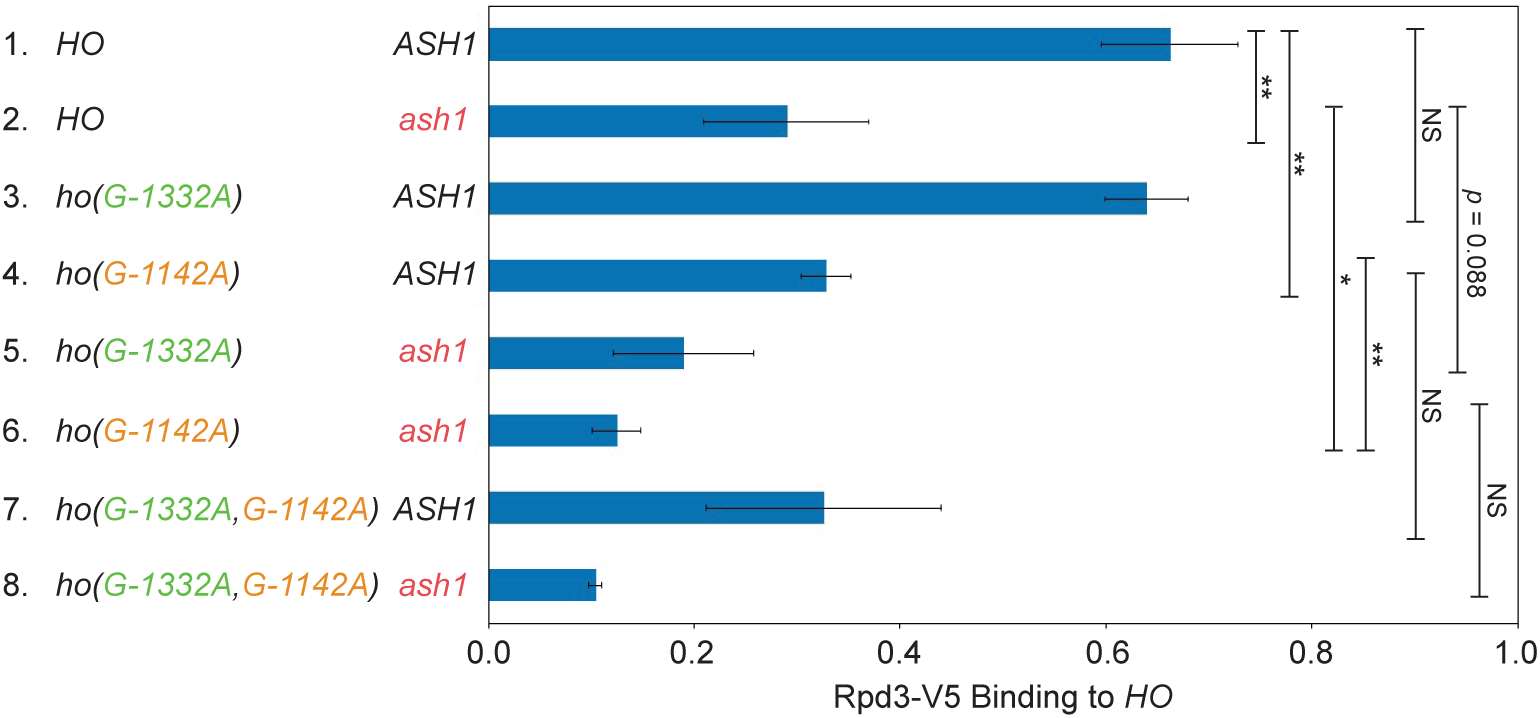
Rpd3 binding is diminished by *ash1* and *ho(G-1142A)* mutants. Binding of Rpd3-V5 to the *HO* promoter in single and combination mutant strains was determined by ChIP, followed by qPCR with primers that amplify *HO* sequence from −1295 to −1121. The strains in this experiment have mutations at all nine SBF sites within URS2 (*9xSBFmut*), to eliminate any recruitment of Rpd3 to the URS2 region by Rpd3. Rpd3-V5 enrichment for each sample was normalized to that of the promoter of *INO1*, and to the corresponding input sample and graphed relative to wild type enrichment. Strain genotypes are indicated on the left. Error bars show the standard deviation of 3 biological replicates. \**p* < 0.05, \*\**p* < 0.01.

The discrepancy between the strong effect on *HO* expression observed in an *ho(G-1332A)* mutant (Table 4, Fig 9) and the apparent negligible effect on Rpd3 recruitment could suggest that Dot6 plays a separate, yet undetermined, repressive role at *HO*. Dot6 and/or Tod6 could repress non-Ribi genes such as *HO* using an Rpd3-independent mechanism. Alternatively, the Rpd3 that associates with *HO* via Dot6 may be difficult to measure at *HO* if the Dot6 interaction itself is more transient than Ash1 and Ume6, as suggested by the lack of consistent detection of Dot6 by ChIP in asynchronous, non-heat shocked cells.

### Negative regulators additively contribute to repression of HO by decreasing SWI/SNF recruitment

The crucial importance of URS1 for *HO* expression is recruitment of the SWI/SNF remodeling complex, which causes waves of nucleosome eviction along the promoter, uncovering the SBF sites within URS2 and ultimately allowing recruitment of RNA polymerase and transcription of the gene. We therefore examined whether mutation of each of these repressors would additively increase SWI/SNF recruitment in the context of a *swi5* mutant. ChIP assays were performed to measure Swi2-V5 recruitment to URS1 in a set of mutant strains, and cells were collected from the same cultures for parallel analysis of *HO* RNA expression (Fig 11). Swi2-V5 recruitment and *HO* expression were both very low in a *swi5* strain, relative to wild type (compare Line 2 to Line 1), suggesting that the Ace2 activator is unable to promote high *HO* expression because it does not recruit sufficient SWI/SNF for nucleosome eviction to uncover the SBF sites in URS2. Each single mutant representing reduction or loss of Ash1, Dot6 and Ume6 increased SWI/SNF recruitment to a level that closely matched the corresponding level of *HO* expression increase (compare Lines 3-5 to Line 2). Binding of Swi2-V5 to URS2 was weaker than at URS1 in all strains(Fig S6A), as expected (Takahata *et al*. 2009b). Similar trends of changes in Swi2-V5 binding between each mutant and wild type were observed at URS1 and URS2 (Fig S6B), as expected if SWI/SNF is recruited to URS1 and spreads downstream to URS2. Ace2 is thus more capable of recruiting SWI/SNF to URS1 in the absence of the repressors, restoring the cascade of nucleosome eviction that extends into URS2 to allow SBF binding and *HO* transcription.

**Figure 11.**
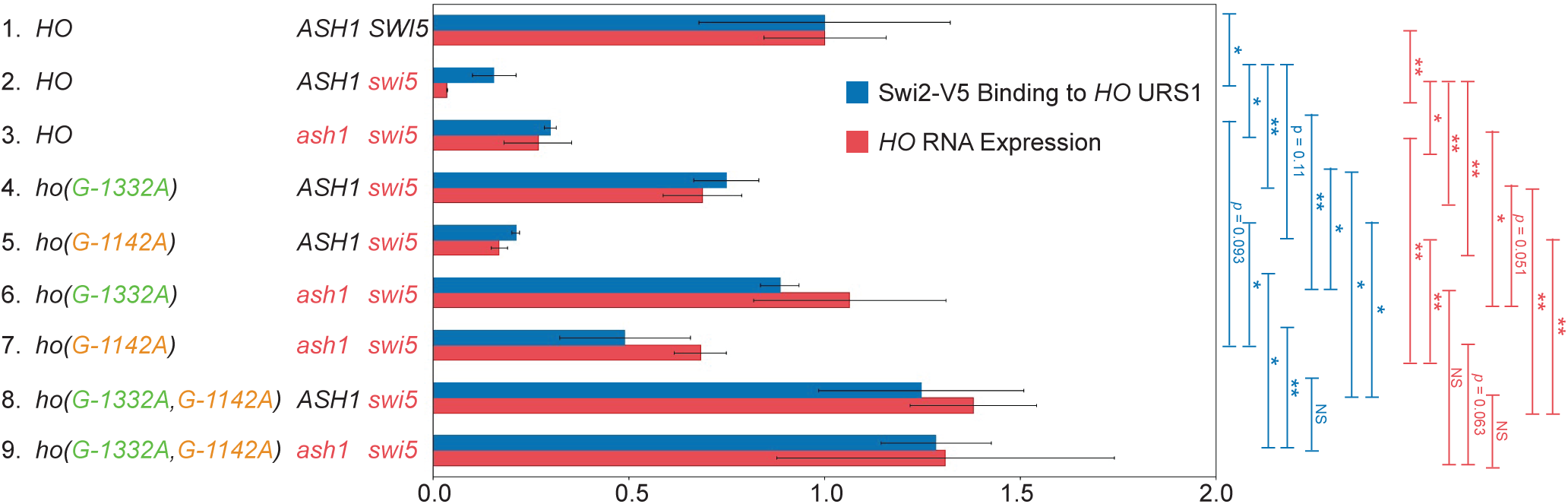
Swi2 binding is restored in *ash1*, *ho(G-1332A)*, and *ho(G-1142A)* mutants. Shown is a comparison of Swi2-V5 binding at the *HO* promoter as determined by ChIP (blue) with *HO* RNA expression (red) in mutant strains. Cells for ChIP and RNA originated from the same cultures that were split upon harvesting. Strain genotypes are indicated on the left. Error bars show the standard deviation of 3 biological replicates. The qPCR following ChIP was performed using primers that amplify *HO* sequence from −1429 to −1158. Swi2-V5 enrichment for each sample was normalized to that of the promoter of *CTS1*, and to the corresponding input sample and graphed relative to wild type enrichment. *HO* expression was normalized to *RPR1* and expressed relative to wild type. \**p* < 0.05, \*\**p* < 0.01.

**Figure 12.**
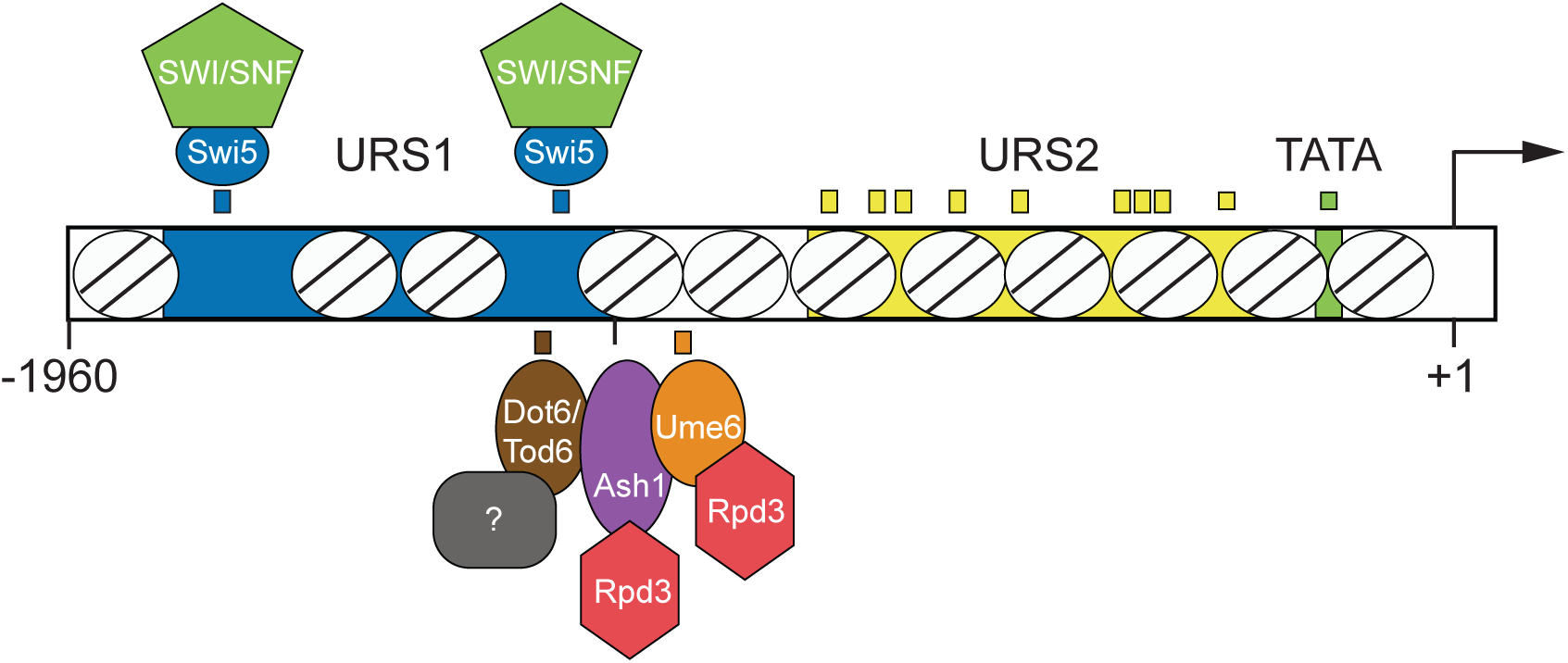

The *ho(G-1332A)* and *ho(G-1142A)* mutations were each additive with *ash1* for recruitment of Swi2-V5 and *HO* expression (Fig 11, compare Line 6 to lines 3 and 4 and Line 7 to Lines 3 and 5). The *ho(G-1332A, G-1142A)* double mutant was also strongly additive for an increase in Swi2-V5 binding, but Swi2-V5 binding did not increase further in the *ho(G-1332A, G-1142A) ash1* triple mutant (compare Line 8 to Lines 4 and 5 and Line 9 to Line 8). These SWI/SNF binding and *HO* expression results are consistent with the *HO* expression data in Fig 9, and further illustrate that if both complexes at the PAC site and Ume6 site are reduced or eliminated in the absence of Swi5, *HO* expression is increased to a point in which an *ash1* mutation has no further effect. Additionally, the observation that *ho(G-1332A)* had strong effects on Swi2-V5 recruitment but not on Rpd3-V5 recruitment are consistent with the idea that proteins bound at the PAC site employ an Rpd3-independent mechanism for repression.

### Concluding remarks: A large number of different repressors is necessary for complex regulation

Our Swi2 ChIP results suggest that the purpose of the Ash1, Ume6 and PAC site complexes is to limit the spread of SWI/SNF-mediated nucleosome eviction from URS1 to URS2. This could ensure that the URS2 and TATA regions of the promoter are not opened prematurely or kept open longer than necessary to achieve the appropriate number of *HO* transcripts. The observation that Ash1, Dot6, and Ume6 associate with the promoter after the initiating event of Swi5 binding, but before the rise in *HO* RNA levels, suggests that their effect is likely on recruitment of SWI/SNF itself rather than on recruitment of the Swi5 activator, which would in turn affect the level of SWI/SNF. Consistent with this idea, we have found that Swi5 levels at the *HO* promoter do not change in response to Ash1 levels or *HO* promoter mutants (Fig S7).

Extending this view to the larger picture of repression at the *HO* promoter, we suggest that the primary role of the cohort of negative regulators is to restrict the recruitment and/or action of coactivator complexes. The number and diversity of regulators suggests there are multiple mechanisms to accomplish this goal, targeting different coactivators at a range of locations along the promoter. The Rpd3 complex brought by Ash1 and Ume6 would limit the action of the Gcn5 histone acetyltransferase within the SAGA coactivator. The Hda1 complex, for which we also identified mutants in our screen, likely has a similar function. Because SAGA levels affect recruitment of SWI/SNF, these histone deacetylase complexes could, by extension, also affect the levels of SWI/SNF, as we observed in our Swi2 ChIP experiments. The Isw2 chromatin remodeler (also likely to play a role at *HO* since mutants were identified in our screen) may more directly oppose SWI/SNF action. Ume6 is known to associate with Isw2 throughout the genome, and thus Ume6 could serve as a recruitment site for both Rpd3 and Isw2. Tup1 could also fit within this model, as studies have demonstrated connections between Tup1 and both Hda1 and Rpd3, and it has been shown that Tup1 inhibits SWI/SNF recruitment (Wong and Struhl 2011). Additionally, Tup1 could affect recruitment or function of the Mediator complex, a coactivator that is important at *HO,* but for which a clear connection to any of the other repressors identified in our screens has not been established.

The majority of Ash1 binding within the *HO* promoter is within the nucleosome just upstream of the linker containing the Ume6 site (data not shown; manuscript in preparation). The position of Ash1 and Ume6 at the right end of URS1 is consistent with a role in limiting the spread of nucleosome eviction into URS2. Why would *HO* require two methods of recruiting Rpd3 to the promoter at similar locations? One possible reason could be the differential levels of repression in mother versus daughter cells. In daughter cells, strong repression is necessary to keep the gene off. However, in mother cells, a more nuanced mechanism of repression is required to ensure an adequate but not excessive level of nucleosome eviction spreading into URS2. Since Ash1 and Ume6 associate with *HO* after Swi5 has bound, it seems likely that the Swi5 activator binds to the *HO* promoter in both mothers and daughters. Recruitment of coactivators to URS1 and subsequent nucleosome eviction at URS1 would thereby occur in both cell types, but the much higher concentration of Ash1-Rpd3 complex in daughters would prevent nucleosome eviction from ultimately spreading into URS2 and promoting gene activation. In contrast, the small amount of Ash1 in mother cells, in combination with Ume6, would serve as an effective, but not absolute, restraint on the ability of SWI/SNF and SAGA to evict nucleosomes.

The PAC site is located near a Swi5 binding site within the NDR upstream of the Ash1/Ume6 sites. The observation that this site does not affect Rpd3 levels suggests that it limits coactivator recruitment by a currently unknown mechanism, different from that of Ash1 and Ume6. The PAC site is particularly intriguing, since it is highly conserved and substitutions here have the strongest effects of all of our mutations on *HO* expression. There is much more to explore related to the PAC site and its influence on *HO* expression, including whether Tod6 and/or other proteins also bind to this site and what types of repressor complexes may be recruited there. The observation that Dot6 binds at *HO* at a time during the cell cycle when it is not bound to Ribi genes suggests there may be a mechanism that operates specifically at *HO* (or *HO* and a class of other genes) to allow Dot6 binding at this time. Dot6 association with chromatin is suggested to be influenced by its phosphorylation (Huber *et al*. 2011). One possible idea is that a phosphatase recruited to *HO* may transiently dephosphorylate Dot6, allowing Dot6 binding at a time when it would not associate with Ribi genes.

Our screens thus identified many contributors to repression at *HO*: the DNA-binding proteins Ash1, Ume6 and Dot6, as well as other regulatory complexes that oppose transcription, including the Rpd3, Hda1, and Isw2 complexes, the Tup1 corepressor and a possible additional complex located at the position of the Cluster 1 mutations. These proteins appear to associate with a variety of locations within the *HO* promoter, including the nucleosome and linker downstream of Swi5 Site B at the right end of URS1 (Ash1 and Ume6), within the NDR that includes Swi5 Site B (Dot6), and within the NDR that includes Swi5 site A (putative complex at Cluster 1) (Fig 12). The question then becomes why the *HO* promoter requires such a large quantity and diversity of repressors. Many complex genes, such as *HO*, require a series of coactivators with different activities to achieve the appropriate level of transcription. It is not surprising that a parallel set of corepressors could be needed to provide balance, restricting the action of coactivators both temporally and by cell type. While corepressor complexes were initially thought to be exclusively negative in terms of their effect on transcription, increasing evidence suggests they can be required for resetting the promoter for subsequent expression and are therefore an integral part of the activation process (Wang *et al*. 2009; Dovey *et al*. 2010; Wong and Struhl 2011). A delicate balance as well as coordination between activating and repressing factors and processes is a necessary part of complex gene regulation. Further investigation of the repressor complexes at the *HO* promoter may serve to clarify and understand their specific roles and their relationship to one another and to the coactivator complexes.

## Acknowledgements

We thank Tim Formosa and Laura McCullough for comments on the manuscript, Tim Parnell for help with the bioinformatics, and Tim Formosa and members of the Stillman lab for helpful advice throughout the course of this project. We thank Zaily Connell, Tim Formosa, Mark Gartenberg, and Kevin Struhl for plasmids, and Benjamin Albert and David Shore for personal communication. We thank undergraduate students Tom Engar and Jon Navar who contributed to development of the genetic screen. This work was supported by National Institutes of Health grant GM121079, awarded to D.J.S.

