## Supplemental Information for "Genetic screens identify multiple negative regulators that additively restrict recruitment of the SWI/SNF chromatin remodeler to the *HO* promoter in *Saccharomyces cerevisiae*"

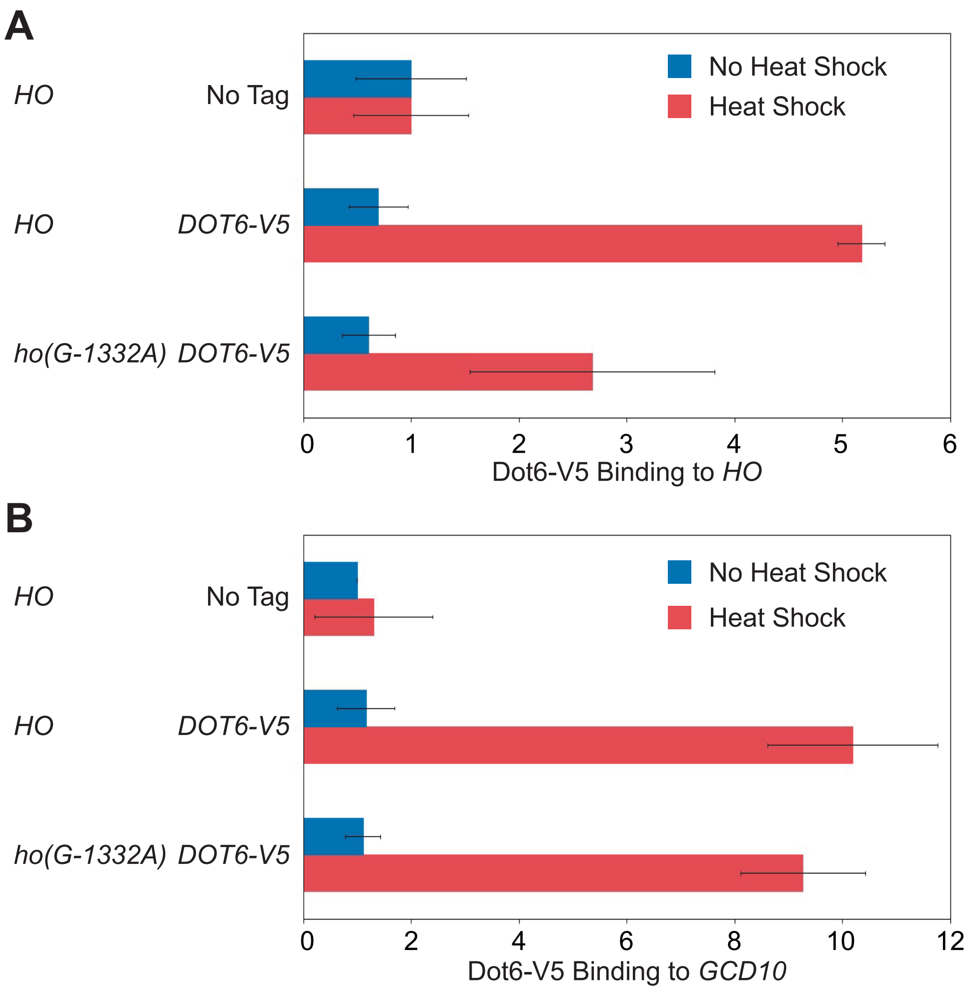


**Supplemental Figure S1. Dot6 binding to the *HO* and *GCD10* promoters by ChIP analysis.**

Binding of Dot6-V5 to *HO* wild type and *ho(G-1332A)* promoters in heat shocked cells was determined by ChIP. This is the same data presented in Figure 5B, with each primer set used in Figure 5B now normalized to a negative reference control, the intergenic region *IGR-I*. Each was also normalized to the corresponding input sample and then graphed relative to the "No Tag" control. Error bars show the standard deviation of 3 biological replicates.

(A) Primers that amplify *HO* sequence from -1471 to -1250 were used.

(B) Primers that amplify *GCD10* promoter sequence were used.

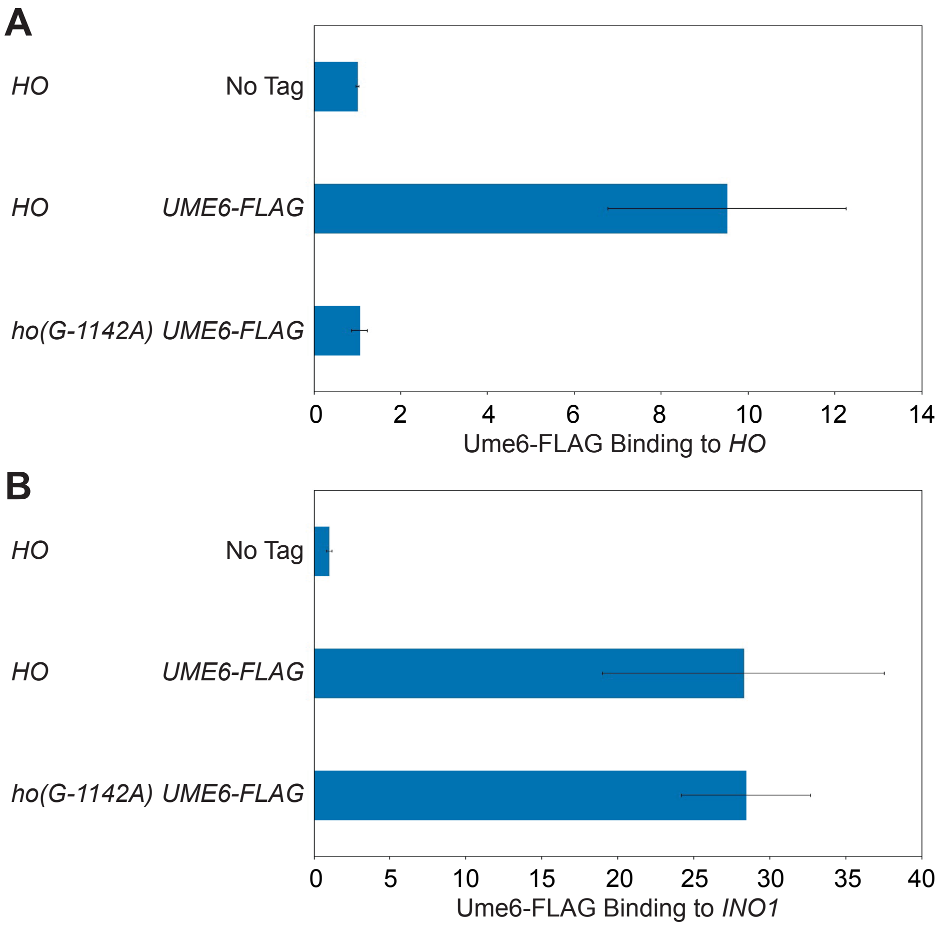


**Supplemental Figure S2. Ume6 binding to the *HO* and *INO1* promoters by ChIP analysis.**

Binding of Ume6-FLAG to *HO* wild type and *ho(G-1142A)* promoters was determined by ChIP. This is the same data presented in Figure 6A, with each primer set used in Figure 6A now normalized to a negative reference control, the coding region of *CDC2*. Each was also normalized to the corresponding input sample and then graphed relative to the "No Tag" control. Error bars show the standard deviation of 3 biological replicates.

(A) Primers that amplify *HO* sequence from -1471 to -1250 were used.

(B) Primers that amplify *INO1* promoter sequence were used.

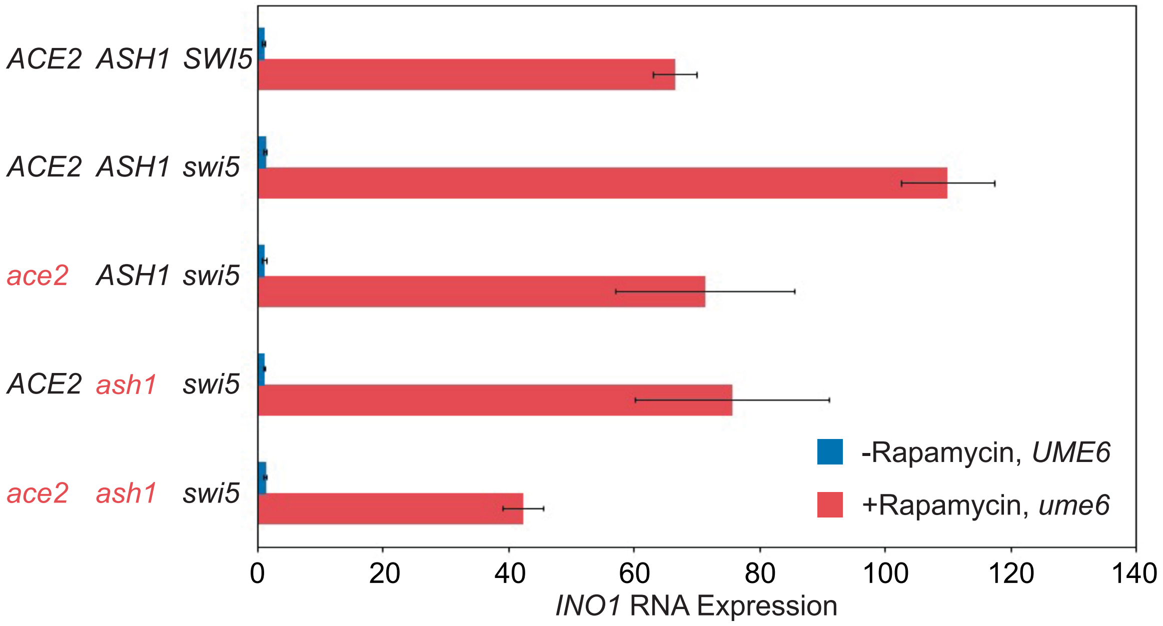


**Supplemental Figure S3. *INO1* expression increases dramatically upon Anchor Away of Ume6.**

This data is a corollary to Figure 6C, which shows *HO* RNA expression in Ume6 Anchor Away conditions. *INO1* mRNA levels were measured, normalized to *RPR1*, and expressed relative to wild type. Error bars represent the standard deviation of at least 3 biological replicates for each strain. See Materials and Methods section for a full description of genotypes and growth conditions. Cultures were split into two, and one half was untreated (-Rapamycin, *UME6*; blue bars), while the other half was treated with 1μg/ml rapamycin for 2 hours (+Rapamycin, *ume6*; red bars).

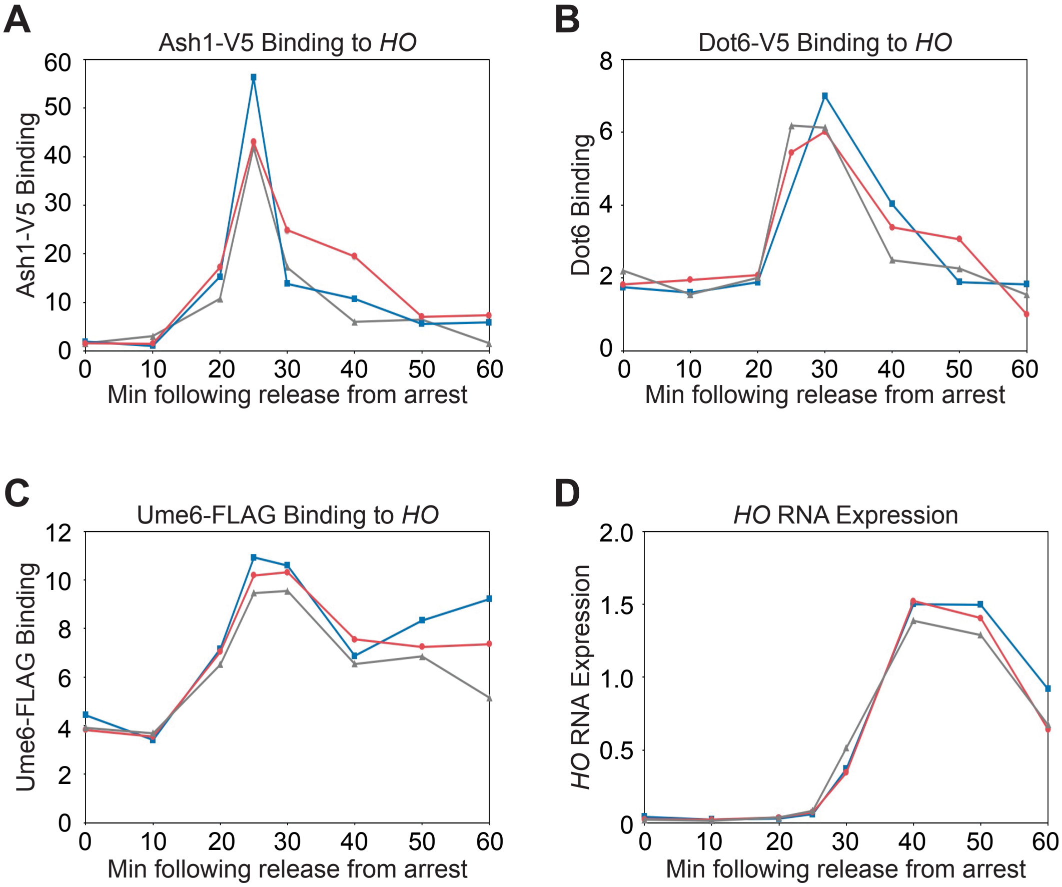


**Supplemental Figure S4. The Ash1, Dot6 and Ume6 repressors associate with the *HO* promoter before the peak of HO expression.**

This figure shows three biological replicate experiments for each panel in Figure 7.

(A, B, C) Binding of Ash1-V5 (A), Dot6-V5 (B), Ume6-FLAG (C) to the *HO* promoter during a cell cycle arrest and release experiment. Cells containing the *GALp::CDC20* allele were synchronized by galactose withdrawal and re-addition. The 0 min time point represents the G2/M arrest, before release with galactose addition. Cells were harvested at the indicated time points following release (x-axis), and samples were processed for ChIP analysis. Enrichment at the *HO* promoter was measured using primers that span from -1295 to -1121 and was normalized to *IGR-I* and to input.

(D) *HO* mRNA expression measured over the course of synchrony experiments (using a Dot6-V5 strain as an example) and normalized to *RPR1*.

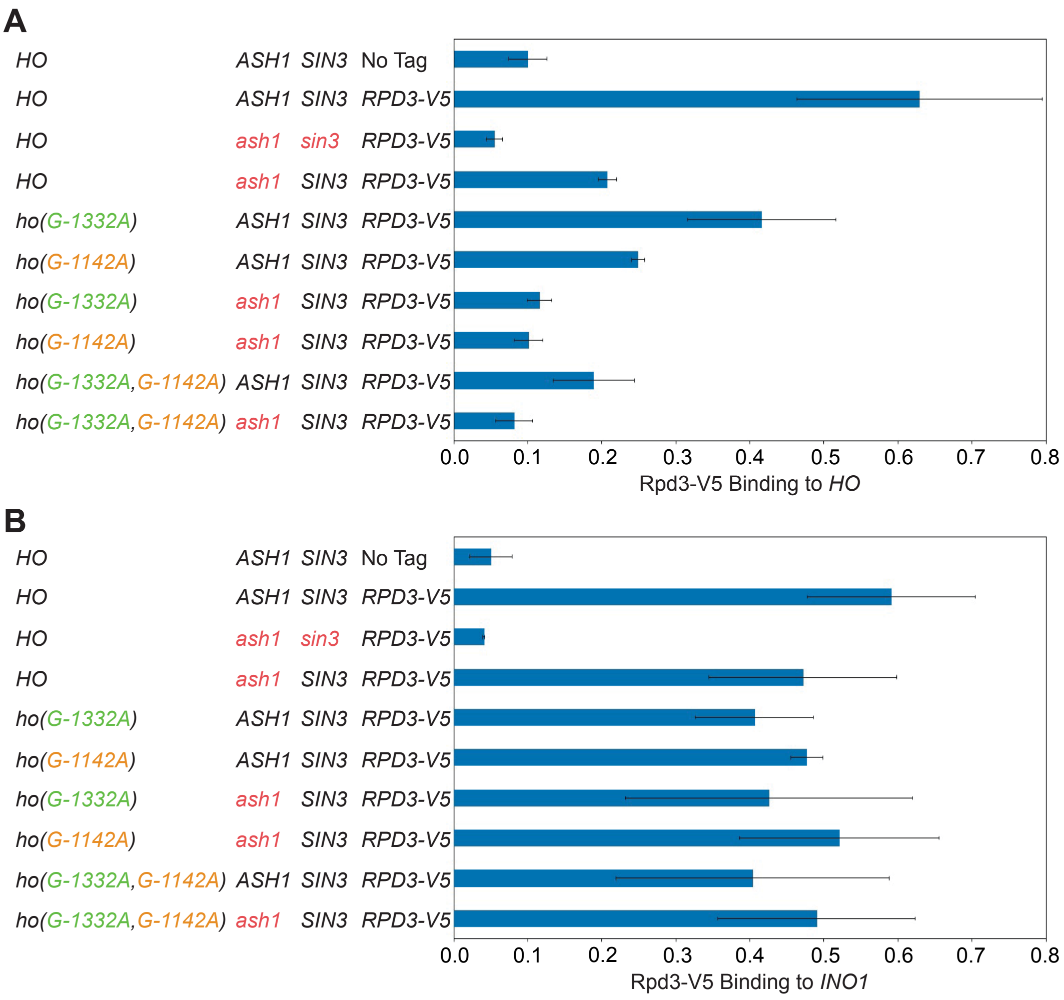


**Supplemental Figure S5. Rpd3 binding to the *HO* and *INO1* promoters by ChIP analysis.**

Binding of Rpd3-V5 in single and combination mutant strains was determined by ChIP. This is the same data presented in Figure 10, with each primer set used in Figure 10 now normalized to a negative reference control, the intergenic region *IGR-I*. Each was also normalized to the corresponding input sample and then graphed relative to the “No Tag” control. Error bars show the standard deviation of 3 biological replicates.

(A) Primers that amplify *HO* sequence from -1471 to -1250 were used.

(B) Primers that amplify *INO1* promoter sequence were used.

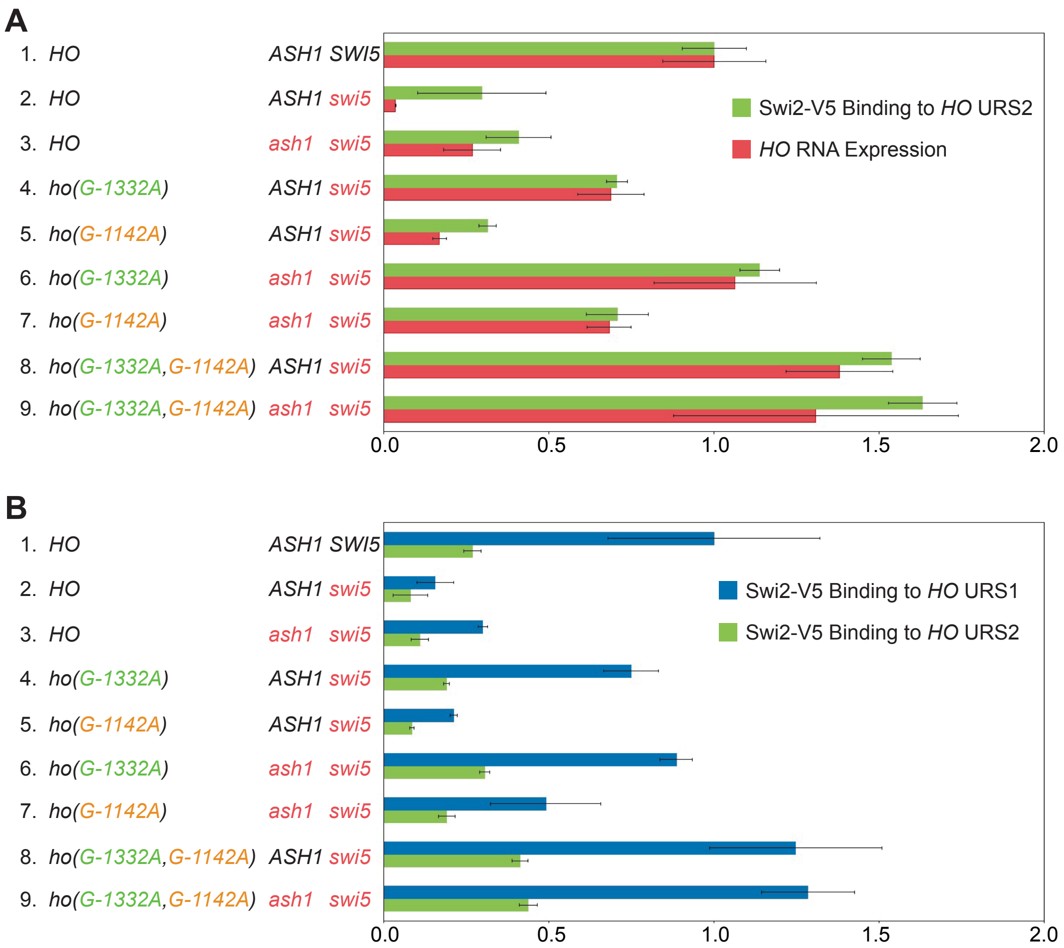


**Supplemental Figure S6. Swi2 binding to *HO* URS2 by ChIP analysis.**

(A) Binding of Swi2-V5 to *HO* URS2 by ChIP analysis (green), compared with *HO* RNA expression levels (red) in mutant strains. Cells for ChIP and RNA originated from the same cultures that were split upon harvesting; these samples are identical to those used in Fig 11. Strain genotypes are indicated on the left. Error bars show the standard deviation of 3 biological replicates. The qPCR following ChIP was performed using primers that amplify *HO* sequence from -897 to -798. Swi2-V5 enrichment for each sample was normalized to that of the promoter of *CTS1*, and to the corresponding input sample and graphed relative to wild type enrichment. *HO* expression was normalized to *RPR1* and expressed relative to wild type.

(B) Comparison of binding of Swi2-V5 to *HO* URS1 (blue) vs. *HO* URS2 (green), using the same data presented in Figs 11 and S6A. Swi2-V5 enrichment for each sample was normalized to that of the promoter of *CTS1*, and to the corresponding input sample and graphed relative to wild type enrichment at URS1.

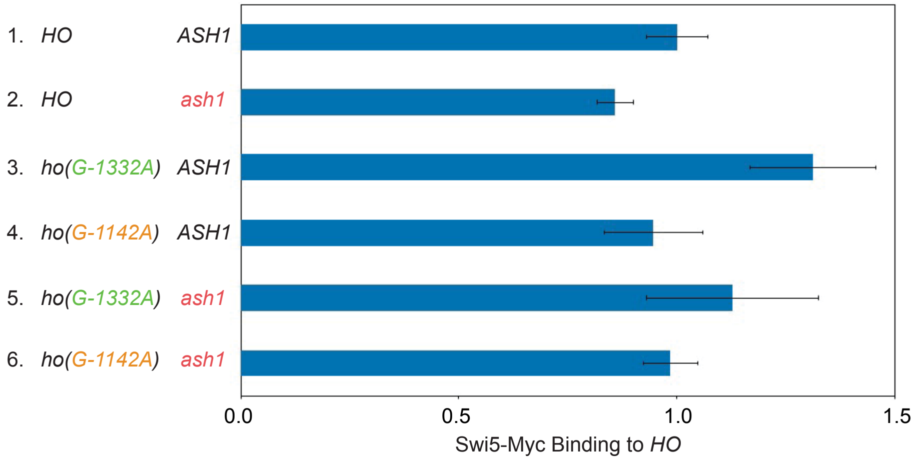


**Supplemental Figure S7. Swi5 binding to *HO* URS1 in *ash1* and *ho* promoter mutants.**

Binding of Swi5-Myc to *HO* URS1 was determined by ChIP, followed by qPCR with primers that amplify *HO* sequence from -1429 to -1158. Swi5-Myc enrichment for each sample was normalized to enrichment at the promoter of *EGT2* and to the corresponding input sample and then graphed relative to wild type. Error bars show the standard deviation of 4 biological replicates.

**Supplemental Table S1. Strains used in this study.**

| | |
|---|---|
| Figure 1C | |
| DY9484 | *MATα HO-ADE2 HO-CAN1 ade2::HphMX his3 leu2 lys2 met15 trp1 ura3* |
| DY10171 | *MATα HO-ADE2 HO-CAN1 swi5::LEU2 RS::TRP1::ACE2::HIS3::RS ade2::HphMX his3 leu2 lys2 met15 trp1 ura3* |
| DY13589 | *MATα HO-ADE2 HO-CAN1 swi5::KanMX RS::TRP1::ACE2::HIS3::RS ash1::LEU2 ade2::HphMX his3 leu2 lys2 met15 trp1 ura3* |
| EY81 | *MATα ho(G-1332A)-ADE2 HO-CAN1 swi5::LEU2 RS::TRP1::ACE2::HIS3::RS ade2::HphMX his3 leu2 lys2 met15 trp1 ura3* |
| DY10239 | *MATα HO-ADE2 HO-CAN1 ace2::HIS3 ade2::HphMX his3 leu2 lys2 met15 trp1 ura3* |
| EY106 | *MATα HO-ADE2 HO-CAN1 swi5::LEU2 ace2::RS ade2::HphMX his3 leu2 lys2 met15 trp1 ura3* |
| EY455 | *MATα HO-ADE2 HO-CAN1 swi5::KanMX ash1::LEU2 ace2::RS ade2::HphMX his3 leu2 lys2 met15 trp1 ura3* |
| EY320 | *MATα ho(G-1332A)-ADE2 HO-CAN1 swi5::LEU2 ace2::RS ade2::HphMX his3 leu2 lys2 met15 trp1 ura3* |
| Figure 1D | |
| DY9484 | *MATα HO-ADE2 HO-CAN1 ade2::HphMX his3 leu2 lys2 met15 trp1 ura3* |
| DY13589 | *MATα HO-ADE2 HO-CAN1 swi5::KanMX ash1::LEU2 RS::TRP1::ACE2::HIS3::RS ade2::HphMX his3 leu2 lys2 met15 trp1 ura3* |
| EY424 | *MATα HO-ADE2 HO-CAN1 swi5::KanMX ash1::LEU2 hda3(T418I) RS::TRP1::ACE2::HIS3::RS ade2::HphMX his3 leu2 lys2 met15 trp1 ura3* |
| EY474 | *MATα HO-ADE2 HO-CAN1 swi5::KanMX ash1::LEU2 isw2(R250I) RS::TRP1::ACE2::HIS3::RS ade2::HphMX his3 leu2 lys2 met15 trp1 ura3* |
| EY468 | *MATα HO-ADE2 HO-CAN1 swi5::KanMX ash1::LEU2 rpd3(Y347N) RS::TRP1::ACE2::HIS3::RS ade2::HphMX his3 leu2 lys2 met15 trp1 ura3* |

| | |
|---|---|
| EY421 | *MATα HO-ADE2 HO-CAN1 swi5::KanMX ash1::LEU2 tup(H575Y) RS::TRP1::ACE2::HIS3::RS ade2::HphMX his3 leu2 lys2 met15 trp1 ura3* |
| DY10239 | *MATα HO-ADE2 HO-CAN1 ace2::HIS3 ade2::HphMX ade2 his3 leu2 lys2 met15 trp1 ura3* |
| EY455 | *MATα HO-ADE2 HO-CAN1 swi5::KanMX ash1::LEU2 ace2::RS ade2::HphMX his3 leu2 lys2 met15 trp1 ura3* |
| EY445 | *MATα HO-ADE2 HO-CAN1 swi5::KanMX ash1::LEU2 hda3(T418I) ace2::RS ade2::HphMX his3 leu2 lys2 met15 trp1 ura3* |
| EY495 | *MATα HO-ADE2 HO-CAN1 swi5::KanMX ash1::LEU2 isw2(R250I) ace2::RS ade2::HphMX his3 leu2 lys2 met15 trp1 ura3* |
| EY489 | *MATα HO-ADE2 HO-CAN1 swi5::KanMX ash1::LEU2 rpd3(Y347N) ace2::RS ade2::HphMX his3 leu2 lys2 met15 trp1 ura3* |
| EY442 | *MATα HO-ADE2 HO-CAN1 swi5::KanMX ash1::LEU2 tup(H575Y) ace2::RS ade2::HphMX his3 leu2 lys2 met15 trp1 ura3* |
| Figure 2 | |
| DY150 | *MAT***a** *ade2 can1 his3 leu2 trp1 ura3* |
| DY408 | *MAT***a** *swi5::hisG-URA3-hisG ade2 can1 his3 leu2 trp1 ura3* |
| DY1137 | *MAT***a** *swi5::hisG-URA3-hisG ace2::HIS3 ade2 can1 his3 leu2 trp1 ura3* |
| DY4846 | *MAT***a** *swi5::hisG-URA3-hisG ash1::LEU2 ade2 can1 his3 leu2 trp1 ura3* |
| DY5431 | *MATα swi5::hisG-URA3-hisG ash1::LEU2 ace2::HIS3 ade2 can1 his3 leu2 lys2 trp1 ura3* |
| DY10941 | *MAT***a** *ho(G-1332A)::KanMX(3') ade2 can1 his3 leu2 trp1 ura3* |
| DY11305 | *MAT***a** *ho(G-1332A)::KanMX(3') swi5::LEU2 ade2 can1 his3 leu2 met15 trp1 ura3* |
| DY11307 | *MAT***a** *ho(G-1332A)::KanMX(3') swi5::LEU2 ace2::HIS3 ade2 can1 his3 leu2 met15 trp1 ura3* |
| DY12656 | *MAT***a** *ho(G-1332A)::KanMX(3') swi5::TRP1 ash1::LEU2 ade2 can1 his3 leu2 trp1 ura3* |

| | |
|---|---|
| DY12658 | *MAT***a** *ho(G-1332A)::KanMX(3') swi5::TRP1 ash1::LEU2 ace2::HIS3 ade2 can1 his3 leu2 lys2 trp1 ura3* |
| Table 2 | |
| DY150 | *MAT***a** *ade2 can1 his3 leu2 trp1 ura3* |
| DY4846 | *MAT***a** *swi5::hisG-URA3-hisG ash1::LEU2 ade2 can1 his3 leu2 trp1 ura3* |
| DY14347 | *MAT***a** *swi5::TRP1 ash1::LEU2 ace2::HIS3 ade2 can1 his3 leu2 lys2 trp1 ura3* |
| DY15413 | *MAT*α *TOR1-1 fpr1::NatMX RPL13A-2xFKBP12::TRP1 CKS1::FRB::KanMX ade2 can1 his3 leu2 trp1 ura3* |
| DY15492 | *MAT*α *TOR1-1 fpr1::NatMX RPL13A- 2xFKBP12::TRP1 CKS1::FRB::KanMX swi5::hisG-URA3-hisG ash1::LEU2 ade2 can1 his3 leu2 trp1 ura3* |
| DY15489 | *MAT***a** *TOR1-1 fpr1::NatMX RPL13A- 2xFKBP12::TRP1 CKS1::FRB::KanMX swi5::hisG-URA3-hisG ash1::LEU2 ace2::HIS3 ade2 can1 his3 leu2 trp1 ura3* |
| DY14851 | *MAT***a** *hda1::URA3 swi5::TRP1 ash1::LEU2 ade2 can1 his3 leu2 lys2 trp1 ura3* |
| DY14849 | *MAT***a** *hda1::URA3 swi5::TRP1 ash1::LEU2 ace2::HIS3 ade2 can1 his3 leu2 lys2 trp1 ura3* |
| DY14496 | *MAT***a** *hda3::KanMX swi5::TRP1 ash1::LEU2 ade2 can1 his3 leu2 trp1 ura3* |
| DY14492 | *MAT***a** *hda3::KanMX swi5::TRP1 ash1::LEU2 ace2::HIS3 ade2 can1 his3 leu2 trp1 ura3* |
| DY14237 | *MAT***a** *isw2::LEU2 swi5::hisG-URA3-hisG ash1::TRP1 ade2 can1 his3 leu2 trp1 ura3* |
| DY14241 | *MAT***a** *isw2::LEU2 swi5::hisG-URA3-hisG ash1::TRP1 ace2::HIS3 ade2 can1 his3 leu2 trp1 ura3* |
| DY14865 | *MAT***a** *rpd3::LEU2 swi5::hisG-URA3-hisG ash1::TRP1 ade2 can1 his3 leu2 lys2 trp1 ura3* |
| DY14862 | *MAT***a** *rpd3::LEU2 swi5::hisG-URA3-hisG ash1::TRP1 ace2::HIS3 ade2 can1 his3 leu2 trp1 ura3* |

| | |
|---|---|
| DY16723 | *MAT***a** *sfp1::HphMX swi5::TRP1 ash1::LEU2 ade2 can1 his3 leu2 trp1 ura3* |
| DY16708 | *MAT***a** *sfp1::HphMX swi5::KanMX ash1::LEU2 ace2::HIS3 ade2 can1 his3 leu2 trp1 ura3* |
| DY16764 | *MAT***a** *tup1(H575Y) swi5::KanMX ash1::LEU2 ade2 can1 his3 leu2 met15 trp1 ura3* |
| DY16762 | *MAT***a** *tup1(H575Y) swi5::KanMX ash1::LEU2 ace2::HIS3 ade2 can1 his3 leu2 trp1 ura3* |
| DY15373 | *MAT***a** *TOR1-1 fpr1::NatMX RPL13A-2xFKBP12::TRP1 UME6::FRB::KanMX ade2 can1 his3 leu2 trp1 ura3* |
| DY15361 | *MAT***a** *TOR1-1 fpr1::NatMX RPL13A-2xFKBP12::TRP1 UME6::FRB::KanMX swi5::hisG-URA3-hisG ash1::LEU2 ade2 can1 his3 leu2 trp1 ura3* |
| DY15359 | *MAT***a** *TOR1-1 fpr1::NatMX RPL13A-2xFKBP12::TRP1 UME6::FRB::KanMX swi5::hisG-URA3-hisG ash1::LEU2 ace2::HIS3 ade2 can1 his3 leu2 trp1 ura3* |
| DY14784 | *MAT***a** *dot6::NatMX tod6::KanMX swi5::hisG-URA3-hisG ash1::TRP1 ade2 can1 his3 leu2 trp1 ura3* |
| DY14782 | *MAT***a** *dot6::NatMX tod6::KanMX swi5::hisG-URA3-hisG ash1::TRP1 ace2::HIS3 ade2 can1 his3 leu2 trp1 ura3* |
| Figure 3 | |
| DY150 | *MAT***a** *ade2 can1 his3 leu2 trp1 ura3* |
| DY408 | *MAT***a** *swi5::hisG-URA3-hisG ade2 can1 his3 leu2 trp1 ura3* |
| DY16768 | *MAT***a** *tup1(H575Y) swi5::KanMX ade2 can1 his3 leu2 met15 trp1 ura3* |
| DY4846 | *MAT***a** *swi5::hisG-URA3-hisG ash1::LEU2 ade2 can1 his3 leu2 trp1 ura3* |
| DY16764 | *MAT***a** *tup1(H575Y) swi5::KanMX ash1::LEU2 ade2 can1 his3 leu2 met15 trp1 ura3* |
| DY16762 | *MAT***a** *tup1(H575Y) swi5::KanMX ash1::LEU2 ace2::HIS3 ade2 can1 his3 leu2 trp1 ura3* |

| Table 4 | |
|---|---|
| DY150 | *MAT***a** *ade2 can1 his3 leu2 trp1 ura3* |
| DY408 | *MAT***a** *swi5::hisG-URA3-hisG ade2 can1 his3 leu2 trp1 ura3* |
| DY4846 | *MAT***a** *swi5::hisG-URA3-hisG ash1::LEU2 ade2 can1 his3 leu2 trp1 ura3* |
| DY14347 | *MAT***a** *swi5::TRP1 ash1::LEU2 ace2::HIS3 ade2 can1 his3 leu2 lys2 trp1 ura3* |
| EY811 | *MAT***a** *ho(G-1762C)::KanMX(3') swi5::URA3 ash1::LEU2 ade2 can1 his3 leu2 trp1 ura3* |
| EY807 | *MAT***a** *ho(G-1762C)::KanMX(3') swi5::TRP1 ash1::LEU2 ace2::HIS3 ade2 can1 his3 leu2 trp1 ura3* |
| EY826 | *MAT***a** *ho(A-1734G)::KanMX(3') swi5::TRP1 ash1::LEU2 ade2 can1 his3 leu2 trp1 ura3* |
| EY818 | *MAT***a** *ho(A-1734G)::KanMX(3') swi5::TRP1 ash1::LEU2 ace2::HIS3 ade2 can1 his3 leu2 trp1 ura3* |
| EY745 | *MAT***a** *ho(C-1442T)::HphMX(3') swi5::TRP1ash1::LEU2 ade2 can1 his3 leu2 trp1 ura3* |
| EY744 | *MAT***a** *ho(C-1442T)::HphMX(3') swi5::TRP1 ash1::LEU2 ace2::HIS3 ade2 can1 his3 leu2 trp1 ura3* |
| EY740 | *MAT***a** *ho(C-1416T)::HphMX(3') swi5::TRP1 ash1::LEU2 ade2 can1 his3 leu2 trp1 ura3* |
| EY735 | *MAT***a** *ho(C-1416T)::HphMX(3') swi5::hisG-URA3-hisG ash1::LEU2 ace2::HIS3 ade2 can1 his3 leu2 trp1 ura3* |
| EY663 | *MAT***a** *ho(T-1401C)::KanMX(3') swi5::hisG-URA3-hisG ash1::LEU2 ade2 can1 his3 leu2 trp1 ura3* |
| EY667 | *MAT***a** *ho(T-1401C)::KanMX(3') swi5::hisG-URA3-hisG ash1::LEU2 ace2::HIS3 ade2 can1 his3 leu2 trp1 ura3* |

| | |
|---|---|
| EY983 | *MAT***a** *ho(T-1399C)::KanMX(3') swi5::TRP1 ash1::LEU2 ade2 can1 his3 leu2 trp1 ura3* |
| EY979 | *MAT***a** *ho(T-1399C)::KanMX(3') swi5::TRP1 ash1::LEU2 ace2::HIS3 ade2 can1 his3 leu2 trp1 ura3* |
| EY966 | *MAT***a** *ho(G-1358C)::KanMX(3') swi5::hisG-URA3-hisG ash1::LEU2 ade2 can1 his3 leu2 trp1 ura3* |
| EY962 | *MAT***a** *ho(G-1358C)::KanMX(3') swi5::hisG-URA3-hisG ash1::LEU2 ace2::HIS3 ade2 can1 his3 leu2 trp1 ura3* |
| EY936 | *MAT***a** *ho(T-1338C)::KanMX(3') swi5::hisG-URA3-hisG ash1::LEU2 ade2 can1 his3 leu2 trp1 ura3* |
| EY929 | *MAT***a** *ho(T-1338C)::KanMX(3') swi5::hisG-URA3-hisG ash1::LEU2 ace2::HIS3 ade2 can1 his3 leu2 trp1 ura3* |
| EY871 | *MAT***a** *ho(G-1335A)::HphMX(3') swi5::TRP1 ash1::LEU2 ade2 can1 his3 leu2 trp1 ura3* |
| EY868 | *MAT***a** *ho(G-1335A)::HphMX(3') swi5::TRP1 ash1::LEU2 ace2::HIS3 ade2 can1 his3 leu2 trp1 ura3* |
| EY849 | *MAT***a** *ho(A-1334G)::KanMX(3') swi5::URA3 ash1::LEU2 ade2 can1 his3 leu2 trp1 ura3* |
| EY847 | *MAT***a** *ho(A-1334G)::KanMX(3') swi5::TRP1 ash1::LEU2 ace2::HIS3 ade2 can1 his3 leu2 trp1 ura3* |
| EY951 | *MAT***a** *ho(T-1333C)::KanMX(3') swi5::hisG-URA3-hisG ash1::LEU2 ade2 can1 his3 leu2 trp1 ura3* |
| EY947 | *MAT***a** *ho(T-1333C)::KanMX(3') swi5::hisG-URA3-hisG ash1::LEU2 ace2::HIS3 ade2 can1 his3 leu2 trp1 ura3* |
| DY12656 | *MAT***a** *ho(G-1332A)::KanMX(3') swi5::TRP1 ash1::LEU2 ade2 can1 his3 leu2 trp1 ura3* |

| | |
|---|---|
| DY12658 | *MAT***a** *ho(G-1332A)::KanMX(3') swi5::TRP1 ash1::LEU2 ace2::HIS3 ade2 can1 his3 leu2 lys2 trp1 ura3* |
| EY775 | *MAT***a** *ho(A-1331T)::KanMX(3') swi5::hisG-URA3-hisG ash1::LEU2 ade2 can1 his3 leu2 trp1 ura3* |
| EY769 | *MAT***a** *ho(A-1331T)::KanMX(3') swi5::hisG-URA3-hisG ash1::LEU2 ace2::HIS3 ade2 can1 his3 leu2 trp1 ura3* |
| EY672 | *MAT***a** *ho(T-1328A)::KanMX(3') swi5::TRP1 ash1::LEU2 ade2 can1 his3 leu2 trp1 ura3* |
| EY676 | *MAT***a** *ho(T-1328A)::KanMX(3') swi5::TRP1 ash1::LEU2 ace2::HIS3 ade2 can1 his3 leu2 lys2 trp1 ura3* |
| EY922 | *MAT***a** *ho(G-1144A)::KanMX(3') swi5::TRP1 ash1::LEU2 ade2 can1 his3 leu2 trp1 ura3* |
| EY914 | *MAT***a** *ho(G-1144A)::KanMX(3') swi5::TRP1 ash1::LEU2 ace2::HIS3 ade2 can1 his3 leu2 trp1 ura3* |
| EY696 | *MAT***a** *ho(C-1143T)::KanMX(3') swi5::TRP1 ash1::LEU2 ade2 can1 his3 leu2 trp1 ura3* |
| EY725 | *MAT***a** *ho(C-1143T)::HphMX(3') swi5::TRP1 ash1::LEU2 ace2::HIS3 ade2 can1 his3 leu2 trp1 ura3* |
| EY906 | *MAT***a** *ho(G-1142A)::KanMX(3') swi5::TRP1 ash1::LEU2 ade2 can1 his3 leu2 trp1 ura3* |
| EY900 | *MAT***a** *ho(G-1142A)::KanMX(3') swi5::TRP1 ash1::LEU2 ace2::HIS3 ade2 can1 his3 leu2 trp1 ura3* |
| EY890 | *MAT***a** *ho(G-1141A)::KanMX(3') swi5::TRP1 ash1::LEU2 ade2 can1 his3 leu2 trp1 ura3* |
| EY883 | *MAT***a** *ho(G-1141A)::KanMX(3') swi5::TRP1 ash1::LEU2 ace2::HIS3 ade2 can1 his3 leu2 trp1 ura3* |

| | |
|---|---|
| EY840 | *MAT***a** *ho(A-1138G)::KanMX(3') swi5::TRP1 ash1::LEU2 ade2 can1 his3 leu2 trp1 ura3* |
| EY832 | *MAT***a** *ho(A-1138G)::KanMX(3') swi5::TRP1 ash1::LEU2 ace2::HIS3 ade2 can1 his3 leu2 trp1 ura3* |
| Figure 5A | |
| DY150 | *MAT***a** *ade2 can1 his3 leu2 trp1 ura3* |
| DY408 | *MAT***a** *swi5::hisG-URA3-hisG ade2 can1 his3 leu2 trp1 ura3* |
| DY4846 | *MAT***a** *swi5::hisG-URA3-hisG ash1::LEU2 ade2 can1 his3 leu2 trp1 ura3* |
| DY14784 | *MAT***a** *dot6::NatMX tod6::KanMX swi5::hisG-URA3-hisG ash1::TRP1 ade2 can1 his3 leu2 trp1 ura3* |
| Figure 5B | |
| DY17613 | *MAT***a** *DOT6-V5::HIS3MX ade2 can1 his3 leu2 trp1 ura3* |
| DY17611 | *MAT***a** *DOT6-V5::HIS3MX ho(G-1332A)::KanMX(3') ade2 can1 his3 leu2 trp1 ura3* |
| Figure 5C | |
| DY17613 | *MAT***a** *DOT6-V5::HIS3MX ade2 can1 his3 leu2 trp1 ura3* |
| DY17611 | *MAT***a** *DOT6-V5::HIS3MX ho(G-1332A)::KanMX(3') ade2 can1 his3 leu2 trp1 ura3* |
| DY150 | *MAT***a** *ade2 can1 his3 leu2 trp1 ura3* |
| Figure 5D, E | |
| DY6669 | *MAT***a** *GALp::CDC20::ADE2 ade2 can1 his3 leu2 trp1 ura3* |
| DY17662 | *MAT***a** *GALp::CDC20::ADE2 DOT6-V5::HIS3MX ade2 can1 his3 leu2 trp1 ura3* |
| DY17660 | *MAT***a** *GALp::CDC20::ADE2 DOT6-V5::HIS3MX ho(G-1332A)::KanMX(3') ade2 can1 his3 leu2 trp1 ura3* |
| Figure 6A | |
| DY13782 | *MAT***a** *UME6-Flag(3)::URA3::Flag(3) ade2 can1 his3 leu2 trp1 ura3* |
| DY16059 | *MAT***a** *UME6-Flag(3)::URA3::Flag(3) ho(G-1142A)::KanMX(3') ade2 can1 his3 leu2 trp1 ura3* |

| | |
|---|---|
| Figure 6B | |
| DY13782 | *MAT***a** *UME6-Flag(3)::URA3::Flag(3) ade2 can1 his3 leu2 trp1 ura3* |
| DY16059 | *MAT***a** *UME6-Flag(3)::URA3::Flag(3) ho(G-1142A)::KanMX(3') ade2 can1 his3 leu2 trp1 ura3* |
| DY150 | *MAT***a** *ade2 can1 his3 leu2 trp1 ura3* |
| Figure 6C | |
| DY15373 | *MAT***a** *TOR1-1 fpr1::NatMX RPL13A-2xFKBP12::TRP1 UME6::FRB::KanMX ade2 can1 his3 leu2 trp1 ura3* |
| DY15365 | *MAT***a** *TOR1-1 fpr1::NatMX RPL13A-2xFKBP12::TRP1 UME6::FRB::KanMX swi5::hisG-URA3-hisG ade2 can1 his3 leu2 trp1 ura3* |
| DY15363 | *MAT***a** *TOR1-1 fpr1::NatMX RPL13A-2xFKBP12::::TRP1 UME6::FRB::KanMX swi5::hisG-URA3-hisG ace2::HIS3 ade2 can1 his3 leu2 trp1 ura3* |
| DY15361 | *MAT***a** *TOR1-1 fpr1::NatMX RPL13A-2xFKBP12::::TRP1 UME6::FRB::KanMX swi5::hisG-URA3-hisG ash1::LEU2 ade2 can1 his3 leu2 trp1 ura3* |
| DY15359 | *MAT***a** *TOR1-1 fpr1::NatMX RPL13A-2xFKBP12::::TRP1 UME6::FRB::KanMX swi5::hisG-URA3-hisG ash1::LEU2 ace2::HIS3 ade2 can1 his3 leu2 trp1 ura3* |
| Figure 7 | |
| DY18522 | *MAT***a** *GALp::CDC20::ADE2 ASH1-V5::HIS3MX ade2 can1 his3 leu2 trp1 ura3* |
| DY17658 | *MAT***a** *GALp::CDC20::ADE2 UME6-Flag(3)::URA3::Flag(3) ade2 can1 his3 leu2 trp1 ura3* |
| DY17662 | *MAT***a** *GALp::CDC20::ADE2 DOT6-V5::HIS3MX ade2 can1 his3 leu2 trp1 ura3* |
| Figure 8A | |
| DY18407 | *MAT***a** *ASH1-V5::HIS3MX ade2 can1 his3 leu2 trp1 ura3* |
| DY19541 | *MAT***a** *ASH1-V5::HIS3MX ho(G-1332A)::KanMX(3') ade2 can1 his3 leu2 trp1 ura3* |
| DY18708 | *MAT***a** *ASH1-V5::HIS3MX ho(G-1142A)::KanMX(3') ade2 can1 his3 leu2 trp1 ura3* |

| | |
|---|---|
| DY19543 | *MAT***a** *ASH1-V5::HIS3MX ho(G-1332A, G-1142A)::KanMX(3') ade2 can1 his3 leu2 trp1 ura3* |
| Figure 8B | |
| DY17613 | *MAT***a** *DOT6-V5::HIS3MX ade2 can1 his3 leu2 trp1 ura3* |
| DY19537 | *MAT***a** *DOT6-V5::HIS3MX ash1::LEU2 ade2 can1 his3 leu2 trp1 ura3* |
| DY19535 | *MAT***a** *DOT6-V5::HIS3MX ho(G-1142A)::HphMX(3') ade2 can1 his3 leu2 trp1 ura3* |
| DY19533 | *MAT***a** *DOT6-V5::HIS3MX ho(G-1142A)::HphMX(3') ash1::LEU2 ade2 can1 his3 leu2 trp1 ura3* |
| Figure 8C | |
| DY13782 | *MAT***a** *UME6-Flag(3)::URA3::Flag(3) ade2 can1 his3 leu2 trp1 ura3* |
| DY14354 | *MAT***a** *UME6-Flag(3)::URA3::Flag(3) ash1::LEU2 ade2 can1 his3 leu2 trp1 ura3* |
| DY13781 | *MAT***a** *UME6-Flag(3)::URA3::Flag(3) ho(G-1332A)::KanMX(3') ade2 can1 his3 leu2 trp1 ura3* |
| DY19539 | *MAT***a** *UME6-Flag(3)::URA3::Flag(3) ho(G-1332A)::KanMX(3') ash1::LEU2 ade2 can1 his3 leu2 trp1 ura3* |
| Figure 9 | |
| DY150 | *MAT***a** *ade2 can1 his3 leu2 trp1 ura3* |
| DY10941 | *MAT***a** *ho(G-1332A)::KanMX(3') ade2 can1 his3 leu2 trp1 ura3* |
| DY16057 | *MAT***a** *ho(G-1142A)::KanMX(3') ade2 can1 his3 leu2 trp1 ura3* |
| DY16968 | *MAT***a** *ho(G-1332A, G-1142A)::KanMX(3') ade2 can1 his3 leu2 trp1 ura3* |
| DY4394 | *MAT***a** *ash1::LEU2 ade2 can1 his3 leu2 trp1 ura3* |
| DY12581 | *MAT***a** *ho(G-1332A)::KanMX(3') ash1::LEU2 ade2 can1 his3 leu2 trp1 ura3* |
| DY16610 | *MAT***a** *ho(G-1142A)::HphMX(3') ash1::LEU2 ade2 can1 his3 leu2 trp1 ura3* |
| DY16974 | *MAT***a** *ho(G-1332A, G-1142A)::KanMX(3') ash1::LEU2 ade2 can1 his3 leu2 trp1 ura3* |
| DY408 | *MAT***a** *swi5::hisG-URA3-hisG ade2 can1 his3 leu2 trp1 ura3* |

| | |
|---|---|
| DY15892 | *MAT***a** *ho(G-1332A)::KanMX(3') swi5::hisG-URA3-hisG ade2 can1 his3 leu2 trp1 ura3* |
| DY15940 | *MAT***a** *ho(G-1142A)::KanMX(3') swi5::hisG-URA3-hisG ade2 can1 his3 leu2 trp1 ura3* |
| DY16965 | *MAT***a** *ho(G-1332A, G-1142A)::KanMX(3') swi5::TRP1 ade2 can1 his3 leu2 trp1 ura3* |
| DY4846 | *MAT***a** *swi5::hisG-URA3-hisG ash1::LEU2 ade2 can1 his3 leu2 trp1 ura3* |
| DY12656 | *MAT***a** *ho(G-1332A)::KanMX(3') swi5::TRP1 ash1::LEU2 ade2 can1 his3 leu2 trp1 ura3* |
| DY15937 | *MAT***a** *ho(G-1142A)::KanMX(3') swi5::TRP1 ash1::LEU2 ade2 can1 his3 leu2 trp1 ura3* |
| DY16963 | *MAT***a** *ho(G-1332A, G-1142A)::KanMX(3') swi5::TRP1 ash1::LEU2 ade2 can1 his3 leu2 trp1 ura3* |
| Figure 10 | |
| DY17742 | *MAT***a** *RPD3-V5::HIS3MX ho(9xSBFmut) ade2 can1 his3 leu2 trp1 ura3* |
| DY17693 | *MAT***a** *RPD3-V5::HIS3MX ho(9xSBFmut) ash1:LEU2 ade2 can1 his3 leu2 trp1 ura3* |
| DY17744 | *MAT***a** *RPD3-V5::HIS3MX ho(G-1332A, 9xSBFmut)::KanMX(3') ade2 can1 his3 leu2 trp1 ura3* |
| DY18666 | *MAT***a** *RPD3-V5::HIS3MX ho(G-1142A, 9xSBFmut)::KanMX(3') ade2 can1 his3 leu2 trp1 ura3* |
| DY17748 | *MAT***a** *RPD3-V5::HIS3MX ho(G-1332A, 9xSBFmut)::KanMX(3') ash1:LEU2 ade2 can1 his3 leu2 trp1 ura3* |
| DY17752 | *MAT***a** *RPD3-V5::HIS3MX ho(G-1142A, 9xSBFmut) ash1:LEU2 ade2 can1 his3 leu2 trp1 ura3* |
| DY17682 | *MAT***a** *RPD3-V5::HIS3MX ho(G-1332A, G-1142A, 9xSBFmut)::KanMX(3') ade2 can1 his3 leu2 trp1 ura3* |

| | |
|---|---|
| DY17750 | *MAT***a** *RPD3-V5::HIS3MX ho(G-1332A, G-1142A, 9xSBFmut)::KanMX(3') ash1:LEU2 ade2 can1 his3 leu2 trp1 ura3* |
| Figure 11 | |
| DY19207 | *MAT***a** *SWI2-V5::HphMX ade2 can1 his3 leu2 trp1 ura3* |
| DY19250 | *MAT***a** *SWI2-V5::HphMX swi5::TRP1 ade2 can1 his3 leu2 trp1 ura3* |
| DY19330 | *MAT***a** *SWI2-V5::HphMX swi5::TRP1 ash1::LEU2 ade2 can1 his3 leu2 trp1 ura3* |
| DY19247 | *MAT***a** *SWI2-V5::HphMX ho(G-1332A)::KanMX(3') swi5::TRP1 ade2 can1 his3 leu2 trp1 ura3* |
| DY19251 | *MAT***a** *SWI2-V5::HphMX ho(G-1142A):KanMX(3') swi5::TRP1 ade2 can1 his3 leu2 trp1 ura3* |
| DY19491 | *MAT***a** *SWI2-V5::HphMX ho(G-1332A)::KanMX(3') swi5::TRP1 ash1::LEU2 ade2 can1 his3 leu2 trp1 ura3* |
| DY19493 | *MAT***a** *SWI2-V5::HphMX ho(G-1142A):KanMX(3') swi5::TRP1 ash1::LEU2 ade2 can1 his3 leu2 trp1 ura3* |
| DY19326 | *MAT***a** *SWI2-V5::HphMX ho(G-1332A, G-1142A)::KanMX(3') swi5::TRP1 ade2 can1 his3 leu2 trp1 ura3* |
| DY19323 | *MAT***a** *SWI2-V5::HphMX ho(G-1332A, G-1142A)::KanMX(3') swi5::TRP1 ash1::LEU2 ade2 can1 his3 leu2 trp1 ura3* |
| Figure S1A, B | |
| DY17613 | *MAT***a** *DOT6-V5::HIS3MX ade2 can1 his3 leu2 trp1 ura3* |
| DY17611 | *MAT***a** *DOT6-V5::HIS3MX ho(G-1332A)::KanMX(3') ade2 can1 his3 leu2 trp1 ura3* |
| DY150 | *MAT***a** *ade2 can1 his3 leu2 trp1 ura3* |
| Figure S2A, B | |
| DY13782 | *MAT***a** *UME6-Flag(3)::URA3::Flag(3) ade2 can1 his3 leu2 trp1 ura3* |
| DY16059 | *MAT***a** *UME6-Flag(3)::URA3::Flag(3) ho(G-1142A)::KanMX(3') ade2 can1 his3 leu2 trp1 ura3* |
| DY150 | *MAT***a** *ade2 can1 his3 leu2 trp1 ura3* |

| | |
|---|---|
| Figure S3 | |
| DY15373 | *MAT***a** *TOR1-1 fpr1::NatMX RPL13A-2xFKBP12::TRP1 UME6::FRB::KanMX ade2 can1 his3 leu2 trp1 ura3* |
| DY15365 | *MAT***a** *TOR1-1 fpr1::NatMX RPL13A-2xFKBP12::TRP1 UME6::FRB::KanMX swi5::hisG-URA3-hisG ade2 can1 his3 leu2 trp1 ura3* |
| DY15363 | *MAT***a** *TOR1-1 fpr1::NatMX RPL13A-2xFKBP12::::TRP1 UME6::FRB::KanMX swi5::hisG-URA3-hisG ace2::HIS3 ade2 can1 his3 leu2 trp1 ura3* |
| DY15361 | *MAT***a** *TOR1-1 fpr1::NatMX RPL13A-2xFKBP12::::TRP1 UME6::FRB::KanMX swi5::hisG-URA3-hisG ash1::LEU2 ade2 can1 his3 leu2 trp1 ura3* |
| DY15359 | *MAT***a** *TOR1-1 fpr1::NatMX RPL13A-2xFKBP12::::TRP1 UME6::FRB::KanMX swi5::hisG-URA3-hisG ash1::LEU2 ace2::HIS3 ade2 can1 his3 leu2 trp1 ura3* |
| Figure S4 | |
| DY18522 | *MAT***a** *GALp::CDC20::ADE2 ASH1-V5::HIS3MX ade2 can1 his3 leu2 trp1 ura3* |
| DY17658 | *MAT***a** *GALp::CDC20::ADE2 UME6-Flag(3)::URA3::Flag(3) ade2 can1 his3 leu2 trp1 ura3* |
| DY17662 | *MAT***a** *GALp::CDC20::ADE2 DOT6-V5::HIS3MX ade2 can1 his3 leu2 trp1 ura3* |
| Figure S5A, B | |
| DY17742 | *MAT***a** *RPD3-V5::HIS3MX ho(9xSBFmut) ade2 can1 his3 leu2 trp1 ura3* |
| DY17693 | *MAT***a** *RPD3-V5::HIS3MX ho(9xSBFmut) ash1:LEU2 ade2 can1 his3 leu2 trp1 ura3* |
| DY17744 | *MAT***a** *RPD3-V5::HIS3MX ho(G-1332A, 9xSBFmut)::KanMX(3') ade2 can1 his3 leu2 trp1 ura3* |
| DY18666 | *MAT***a** *RPD3-V5::HIS3MX ho(G-1142A, 9xSBFmut)::KanMX(3') ade2 can1 his3 leu2 trp1 ura3* |
| DY17748 | *MAT***a** *RPD3-V5::HIS3MX ho(G-1332A, 9xSBFmut)::KanMX(3') ash1:LEU2 ade2 can1 his3 leu2 trp1 ura3* |

| | |
|---|---|
| DY17752 | *MAT***a** *RPD3-V5::HIS3MX ho(G-1142A, 9xSBFmut) ash1:LEU2 ade2 can1 his3 leu2 trp1 ura3* |
| DY17682 | *MAT***a** *RPD3-V5::HIS3MX ho(G-1332A, G-1142A, 9xSBFmut)::KanMX(3') ade2 can1 his3 leu2 trp1 ura3* |
| DY17750 | *MAT***a** *RPD3-V5::HIS3MX ho(G-1332A, G-1142A, 9xSBFmut)::KanMX(3') ash1:LEU2 ade2 can1 his3 leu2 trp1 ura3* |
| DY17746 | *MAT***a** *RPD3-V5::HIS3MX ho(9xSBFmut) ash1:LEU2 sin3:hisG ade2 can1 his3 leu2 trp1 ura3* |
| DY17753 | *MAT***a** *ho(9xSBFmut) ade2 can1 his3 leu2 trp1 ura3* |
| Figure S6A, B | |
| DY19207 | *MAT***a** *SWI2-V5::HphMX ade2 can1 his3 leu2 trp1 ura3* |
| DY19250 | *MAT***a** *SWI2-V5::HphMX swi5::TRP1 ade2 can1 his3 leu2 trp1 ura3* |
| DY19330 | *MAT***a** *SWI2-V5::HphMX swi5::TRP1 ash1::LEU2 ade2 can1 his3 leu2 trp1 ura3* |
| DY19247 | *MAT***a** *SWI2-V5::HphMX ho(G-1332A)::KanMX(3') swi5::TRP1 ade2 can1 his3 leu2 trp1 ura3* |
| DY19251 | *MAT***a** *SWI2-V5::HphMX ho(G-1142A):KanMX(3') swi5::TRP1 ade2 can1 his3 leu2 trp1 ura3* |
| DY19491 | *MAT***a** *SWI2-V5::HphMX ho(G-1332A)::KanMX(3') swi5::TRP1 ash1::LEU2 ade2 can1 his3 leu2 trp1 ura3* |
| DY19493 | *MAT***a** *SWI2-V5::HphMX ho(G-1142A):KanMX(3') swi5::TRP1 ash1::LEU2 ade2 can1 his3 leu2 trp1 ura3* |
| DY19326 | *MAT***a** *SWI2-V5::HphMX ho(G-1332A, G-1142A)::KanMX(3') swi5::TRP1 ade2 can1 his3 leu2 trp1 ura3* |
| DY19323 | *MAT***a** *SWI2-V5::HphMX ho(G-1332A, G-1142A)::KanMX(3') swi5::TRP1 ash1::LEU2 ade2 can1 his3 leu2 trp1 ura3* |
| Figure S7 | |
| DY5832 | *MAT***a** *SWI5-Myc(8)::KanMX ade2 can1 his3 leu2 trp1 ura3* |

| | |
|---|---|
| DY8749 | *MAT***a** *SWI5-Myc(8)::KanMX ash1::TRP1 ade2 can1 his3 leu2 trp1* |
| DY17188 | *MAT***a** *SWI5-Myc(8)::KanMX ho(G-1332A)::HphMX(3') ade2 can1 his3 leu2 trp1* |
| DY18096 | *MAT***a** *SWI5-Myc(8)::KanMX ho(G-1142A)::HphMX(3') ade2 can1 his3 leu2 trp1* |
| DY17186 | *MAT***a** *SWI5-Myc(8)::KanMX ho(G-1332A)::HphMX(3') ash1::TRP1 ade2 can1 his3 leu2 trp1* |
| DY18098 | *MAT***a** *SWI5-Myc(8)::KanMX ho(G-1142A)::HphMX(3') ash1::TRP1 ade2 can1 his3 leu2 trp1* |
| Materials and Methods | |
| DY10171 | *MATα HO-ADE2 HO-CAN1 swi5::LEU2 RS::TRP1::ACE2::HIS3::RS ade2::HphMX his3 leu2 lys2 met15 trp1 ura3* |
| DY10085 | *MATα HO-ADE2 HO-CAN1 swi5::LEU2 ade2::HphMX his3 leu2 lys2 met15 trp1 ura3* |
| DY10941 | *MAT***a** *ho(G-1332A)::KanMX(3') ade2 can1 his3 leu2 trp1 ura3* |
| DY7874 | *MAT***a** *ho[URA3(-1496 to -1222 deleted)]::KanMX(3') ade2 can1 his3 leu2 trp1 ura3* |
| DY14815 | *MAT***a** *ho[URA3(K. lactis)::KanMX(inserted at -1750)] swi5::TRP1 ash1::LEU2 ade2 can1 his3 leu2 trp1 ura3* |
| DY14198 | *MAT***a** *ho[URA3(-1496 to -1222 deleted)]::KanMX(3') swi5::TRP1 ash1::LEU2 ade2 can1 his3 leu2 trp1 ura3* |
| DY14840 | *MAT***a** *ho[URA3(K. lactis)::KanMX(inserted at -1200)] swi5::TRP1 ash1::LEU2 ade2 can1 his3 leu2 trp1 ura3* |
| DY15937 | *MAT***a** *ho(G-1142A)::KanMX(3') swi5:TRP1 ash1:LEU2 ade2 can1 his3 leu2 trp1 ura3* |
| DY14839 | *MAT***a** *ho[URA3(K. lactis)::KanMX(inserted at -1200)] swi5::TRP1 ash1::LEU2 ade2 can1 his3 leu2 trp1 ura3* |
| Y40343 | *MATα TOR1-1 fpr1::NatMX RPL13A-2xFKBP12::TRP1 ade2 can1 his3 leu2 trp1 ura3* |

| | |
|---|---|
| DY13589 | *MATα HO-ADE2 HO-CAN1 swi5::KanMX ash1::LEU2 RS::TRP1::ACE2::HIS3::RS ade2::HphMX his3 leu2 lys2 met15 trp1 ura3* |
| DY10061 | *MAT***a** *HO-ADE2::KanMX(3') HO-CAN1 swi5::LEU2 ade2 his3 leu2 met15 trp1 ura3* |
| DY13857 | *MAT***a** *HO-ADE2::NatMX(3') HO-CAN1 ash1::LEU2 swi5::KanMX ade2 his3 leu2 met15 trp1 ura3* |
| DY10174 | *MAT***a** *HO-ADE2 HO-CAN1 swi5::LEU2 ace2::HIS3 ade2 his3 leu2 met15 trp1 ura3* |
| DY13855 | *MAT***a** *HO-ADE2::NatMX(3') HO-CAN1 swi5::KanMX ash1::LEU2 ace2::HIS3 ade2 his3 leu2 met15 trp1 ura3* |
| DY9341 | *MATΔ::LEU2 HO-ADE2 HO-CAN1 swi5::KanMX ade2 his3 leu2 met15 ura3* |
| DY13891 | *MATΔ::NATMX HO-ADE2 HO-CAN1 swi5::KanMX ash1::LEU2 ade2 his3 leu2 trp1 ura3* |
| DY13862 | *MATα ho(-1496 to -1130 deleted)-ADE2::KanMX(3') swi5::LEU2 RS::TRP1::ACE2::HIS3::RS ade2::HphMX can1 his3 leu2 lys2 trp1 ura3* |
| DY13863 | *MATα ho[sup4-o(-1496 to -1222 deleted)]-ADE2::KanMX swi5::LEU2 RS::TRP1::ACE2::HIS3::RS ade2::HphMX can1 his3 leu2 lys2 trp1 ura3* |
| DY13925 | *MATα ho[sup4-o(-1496 to -1222 deleted)]-ADE2::KanMX swi5::hisG-URA3-hisG ash1::LEU2 RS::TRP1::ACE2::HIS3::RS ade2::HphMX can1 his3 leu2 lys2 trp1 ura3* |
| DY13832 | *MATα ho[URA3(-1496 to -1222 deleted)]::KanMX(3') swi5::LEU2 RS::TRP1::ACE2::HIS3::RS ade2::HphMX can1 his3 leu2 lys2 trp1 ura3* |

**Supplemental Table S2. Plasmids used in this study.**

| | |
|---|---|
| M5011 | *RS::TRP1::ACE2* in YEp24 (YEp-*URA3*) |
| M5048 | *ACE2::HIS3::RS* in YEp24 (YEp-*URA3*) |
| M4915 | *HO* in YCplac22 (YCp-*TRP1*) |
| M5181 | *ho(G-1332A)-ADE2* in YCplac22 (YCp-*TRP1*) |
| M4531 | *KanMX* inserted 3' to *HO* gene in pGEM7 |
| pZC03 | pFA6a-TEV-6xGly-V5-HIS3MX tagging cassette in pFA6 |
| ZM467 | *Flag(3)::URA3::Flag(3)* tagging cassette in pBluescript SK (Moqtaderi and Struhl 2008) |
| P30578 | pFA6a-FRB-KanMX tagging cassette in pFA6 (Haruki *et al.* 2008) |
| M5053 | pGAL::Z. rouxii recombinase in pRS317 (YCp-*LYS2*) |
| M2291 | *ACE2* in YCplac33 (YCp-*URA3*) |
| pFA6-KanMX4 | KanMX4 disruptor cassette in pFA6 (Wach *et al.* 1994) |
| pAG25-NatMX4 | NatMX4 disruptor cassette in pFA6 (Goldstein and Mccusker 1999) |
| M2291 | *ACE2* in YCplac33 (YCp-*URA3*) |
| M5404 | *ASH1* in pRS306 (YIp-*URA3*) |
| M2026 | *ho[URA3(-1496 to -1222 deleted)]* in pIC19R |
| M346 | *ho(-1496 to -1130 deleted)* in pBR322 |
| M3208 | *ho[sup4-o(-1496 to -1222 deleted)]* in pGEM7zf |
